# The intrinsically disordered transcriptional activation domain of CIITA is functionally tuneable by single substitutions: An exception or a new paradigm?

**DOI:** 10.1101/2023.11.02.565367

**Authors:** Shwetha Sreenivasan, Paul Heffren, Kyung-Shin Suh, Mykola V. Rodnin, Edina Kosa, Aron W. Fenton, Alexey S. Ladokhin, Paul E. Smith, Joseph D. Fontes, Liskin Swint-Kruse

## Abstract

During protein evolution, some amino acid substitutions modulate protein function (“tuneability”). In most proteins, the tuneable range is wide and can be sampled by a set of protein variants that each contains multiple amino acid substitutions. In other proteins, the full tuneable range can be accessed by a set of variants that each contains a single substitution. Indeed, in some globular proteins, the full tuneable range can be accessed by the set of site-saturating substitutions at an individual “rheostat” position. However, in proteins with intrinsically disordered regions (IDRs), most functional studies – which would also detect tuneability – used multiple substitutions or small deletions. In disordered transcriptional activation domains (ADs), studies with multiple substitutions led to the "acidic exposure" model, which does not anticipate the existence of rheostat positions. In the few studies that did assess effects of single substitutions on AD function, results were mixed: The ADs of two full-length transcription factors did *not* show tuneability, whereas a fragment of a third AD was tuneable by single substitutions. Here, we tested tuneability in the AD of full-length human class II transactivator (CIITA). Sequence analyses and experiments showed that CIITA’s AD is an IDR. Functional assays of singly-substituted AD variants showed that CIITA’s function was highly tuneable, with outcomes not predicted by the acidic exposure model. Four tested positions showed rheostat behaviour for transcriptional activation. Thus, tuneability of different IDRs can vary widely. Future studies are needed to illuminate the biophysical features that govern whether an IDR is tuneable by single substitutions.

## INTRODUCTION

Amino acid substitutions enable protein evolution and serve as a means for bioengineering functional variation. Among their outcomes, many amino acid substitutions tune protein function. A function’s full tuneable range spans from wild-type (WT), to partially-impaired, to “dead”; gain-of-function outcomes (better than WT) can also occur. To access the full tuneable range, some proteins require variants with multiple substitutions. In other proteins, the full tuneable range can be accessed by a variety of single substitutions (*e.g.,* ^1^). Furthermore, in some globular proteins, 50-100% of the tuneable functional range can be sampled by a set of substitutions at an individual “rheostat” position (*e.g.,* ^2–9^). It is still unknown whether the existence of rheostat positions requires a classically-folded protein structure or whether they can also be present in intrinsically disordered regions (IDRs).

Indeed, most historical studies of IDRs, including those found in many eukaryotic transcription factors, have assessed functional tuning *via* multiple, simultaneous substitutions (*e.g.,* ^1; 10–16^) or deleting small regions (*e.g.,* ^17–19^). This approach was further justified by studies of the IDR transcriptional activation domains in p53^20^ and peroxisome proliferator–activated receptor ɣ (PPARɣ)^21^, in which all possible single substitutions were created at each position (site-saturating mutagenesis, “SSM”). When their functional outcomes were assessed, single substitutions in the IDRs showed few significant effects,^20; 21^ and examination of SSM results did not identify rheostat positions in these IDRs. These SSM studies, along with observations that IDR activation domains are rich in acidic residues (*e.g.,*^20; 21^), engendered a “rational mutagenesis” approach for further exploring the sequence/function relationship in transcriptional activation domains.

In the rational mutagenesis studies, sets of substitutions were designed to alter the balance of acidic and aromatic/leucine residues and effects on function were monitored.^13–16; 22–24^ The resulting “acidic exposure model” proposed that interspersing acidic residues among hydrophobic motifs allows the IDR to be in equilibrium between at least two conformations: (i) the collapsed conformation, mediated through interactions of the hydrophobic motifs and (ii) the expanded conformation, mediated through repulsion of the acidic residues that make the hydrophobic motifs available for binding partner proteins.^13; 15^ As such, one way for tuning transcriptional activation is to sufficiently alter the balance of acidic/hydrophobic residues so that the equilibrium between the conformations is changed.

Although dominated by multiple substitutions, the data sets arising from rational mutagenesis of various transcriptional activation domains also contained a handful of singly-substituted variants.^13; 15^ In contrast to the SSM studies of p53^20^ and PPARɣ^21^, the set of single substitutions for the Hifα-derived activation domain^15^ showed broad functional tuning (Supplementary Figure 1). The existence of single substitutions with partially-impaired or enhanced function suggests that (i) the IDR activation domains from different transcription factors can have different substitution sensitivities and that (ii) rheostat positions might be present in some IDRs. These hypotheses are further supported by earlier, albeit limited, alanine scans of IDRs, which also revealed singly-substituted variants with partially-diminished or enhanced function ^18; 25–31^.

To test these hypotheses, we chose a transcription factor with different domain arrangements than p53/PPARɣ – the class II major histocompatibility complex transactivator protein (CIITA) from vertebrates.^32; 33^ CIITA differs from other transcription factors in that it lacks a DNA binding domain. Nevertheless, CIITA contains an acidic activation domain at its N-terminus that is critical for transcriptional activation (positions 1-161)^34; 35^ and appears to have the same characteristics as other acidic activation domains.^36^ To carry out its function, CIITA interacts with multiple DNA-bound complexes at the promoters of genes encoding the major histocompatibility class II proteins; CIITA also recruits other factors necessary to modulate chromatin structure and initiate transcription.^37^ Several of these binding interactions are mediated through CIITA’s activation domain. Furthermore, the G58P and A94P variants of CIITA retained 71% and 43%, respectively, of WT’s transcriptional activation^19^; these partially-diminished activities suggest that these two positions could be rheostat positions.

Here, we first showed that the transcriptional activation domain of CIITA lacked persistent secondary structure using sequence analyses, circular dichroism spectropolarimetry and molecular dynamics simulations. Using full-length CIITA, we next selected seven positions, sampling regions with both high and low disorder probabilities, and generated 10-14 single amino acid substitutions at each position. This semi-saturating mutagenesis approach (semiSM) was previously demonstrated to be sufficient for identifying rheostat positions.^38^ Importantly, this study design used unbiased selection of amino acid side chains and thus expands beyond the rational mutagenesis approach used to formulate the acidic exposure model. The seven sets of CIITA variants were then assessed for functional changes using cell-based assays that measured effects on cellular protein concentrations and normalized transcriptional activation.

Results showed that transcriptional activation could be tuned across the full functional range by single substitutions; gain-of-function substitutions (better than WT) were also observed. All seven tested positions were sensitive to individual substitutions; *i.e.,* no position was neutral. Of these, four positions acted as rheostat positions for transcriptional activation. Strikingly, we did not observe any functional trends related to amino acid side chain chemistries, including outcomes expected from the acidic exposure model. However, for two positions, transcriptional activation did correlate with amino acid structural propensities, suggesting that substitutions in this region of CIITA alters transient helical content. Overall, modulating structural change could be one means by which the various functions of some IDRs are tuned by single amino acid substitutions.

## RESULTS

### Sequence and structure analyses indicate that the activation domain of CIITA is intrinsically disordered

Most transcriptional activation domains contain intrinsically disordered regions (IDRs). IDRs were first identified by their unique sequence features,^39^ which include higher fractions of charged residues and structure-breakers (proline and glycine) and fewer hydrophobic residues than globular proteins (reviewed in ^40^). Since these early studies, several computational algorithms have been developed to predict disorder from sequence. For the transcriptional activation domain of CIITA (canonical isoform I; accession NM_001286402; 1130 amino acids; 123.5 kDa), we obtained its disorder probabilities using Predictor of Natural Disordered regions (PONDR).^41^ Much of the N-terminus, which contains the activation domain (residues 1 to 161), has a high likelihood (>0.5) for intrinsic disorder (Figure 1; Supplementary figure 2a; Supplementary table 1). Results from alternative IDR predictors^42–44^ show similar trends across the whole protein (Supplementary figure 2).

**Figure 1.**
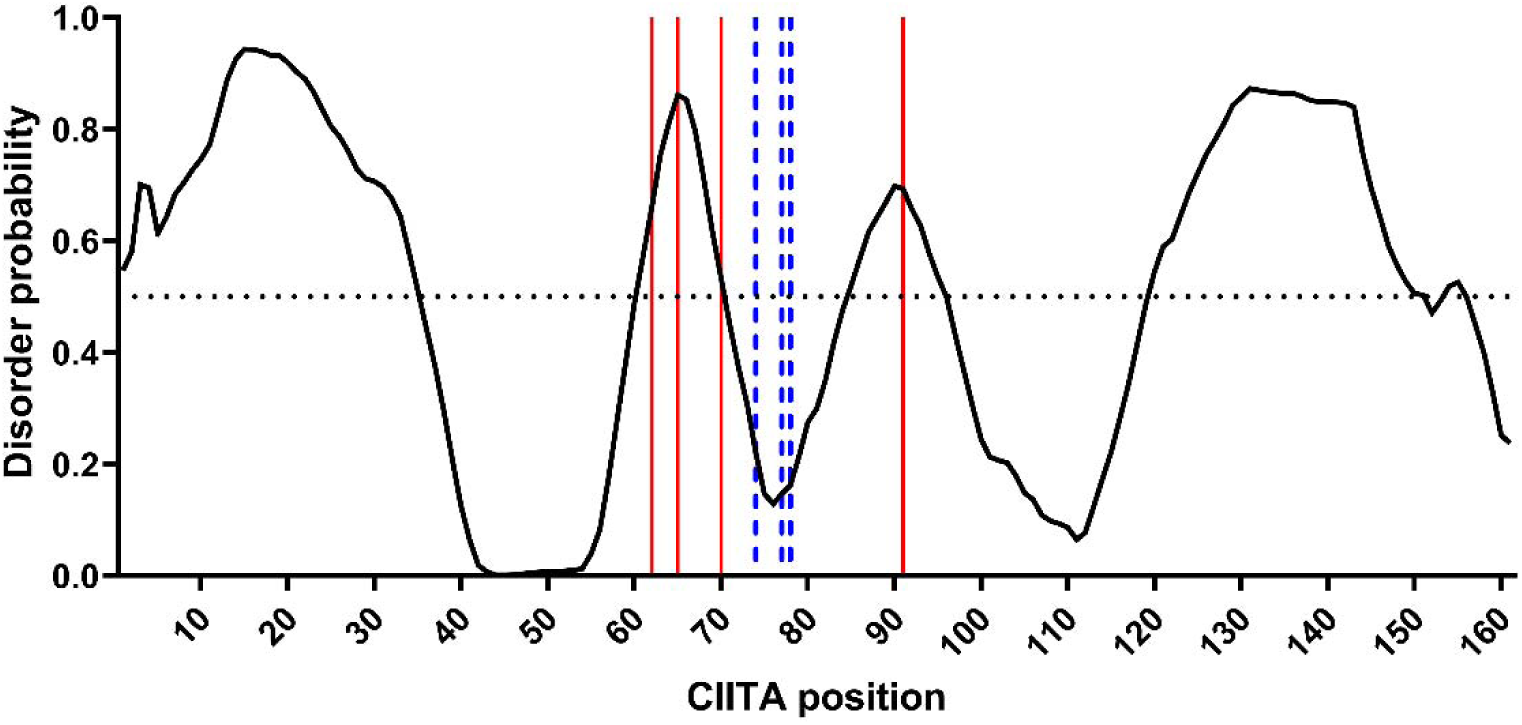
Intrinsic disorder propensities for the N-terminal transcriptional activation domain of CIITA (positions 1-161) by PONDR. Predictions for the full-length protein by PONDR and alternative algorithms are in Supplementary figure 2. Propensity values >0.5 indicate positions with high probability of disorder (e.g., positions 62, 65, 70, 91; solid vertical red lines); values <0.5 indicate positions with low probability of disorder (e.g., positions 74, 77, 78; dashed vertical blue lines). Propensity values for these seven positions are listed in Supplementary table 1.

Another general feature of IDRs is that they appear to lack secondary structures in their unbound states,^45^ although some may fold upon binding ligand.^46–48^ To experimentally assess the secondary structure in the activation domain of CIITA, we expressed and purified fragment 1-210 of CIITA and performed circular dichroism (CD) studies. CD spectra are consistent with this region being primarily random coil (Figure 2), even in the presence of the structure-promoting agent trimethylamine N-oxide (TMAO).

**Figure 2.**
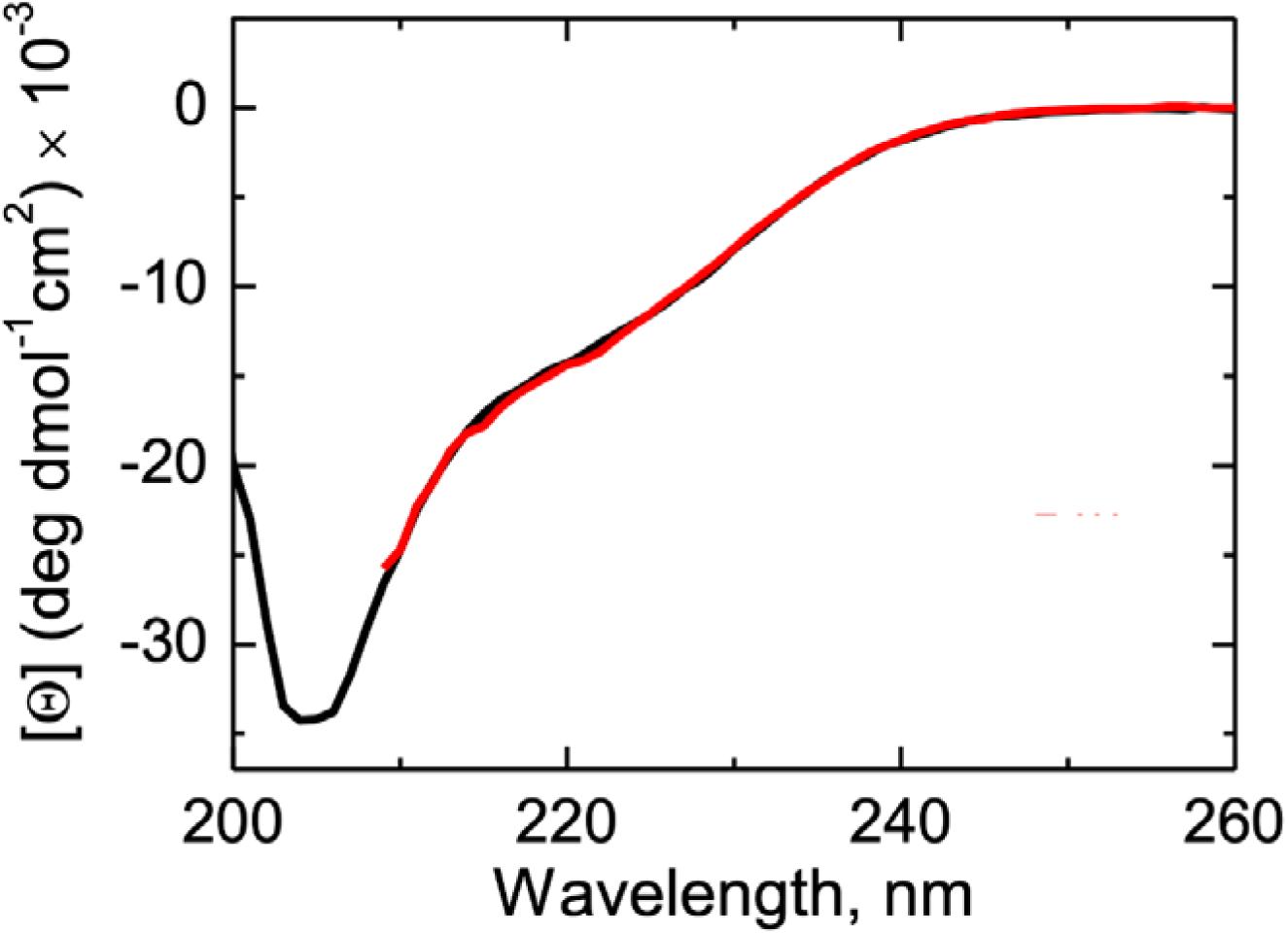
Circular dichroism (CD) of purified CIITA 1-210. The CD spectra of the purified CIITA 1-210 fragment was measured in the absence (black) and presence of 1 M TMAO (red) at room temperature. Samples contained 4 μM CIITA in 50 mM sodium phosphate buffer, pH 8.0. Results shown are the average of 100-150 replicate scans with background subtraction.

To investigate the possibility of persistent residual structure at the atomic level, 5 μs of classical molecular dynamics (MD) simulations of the fragment containing positions 56-94 of CIITA were performed with three different force fields specifically developed and tested for IDRs (Figure 3 and Supplementary figure 3). The radius of gyration and end-to-end distances observed for all three force fields were indicative of peptides with little to no residual long-range structure, as expected for an IDR. This result was consistent for multiple independent simulations using different starting structures for each force field. The only exception was a single simulation using the CHARMM36m force field, which predicted a more collapsed structure (Supplementary figure 3). However, this collapsed structure did not display any significant secondary structure elements when the trajectory was analysed with Define Secondary Structure of Proteins (DSSP)^49; 50^. An analysis of the helical content for each force field, averaged over all three simulations, is shown in Figure 3. The KBFF20 force field suggested some transient helical content for several regions of the peptide. Together, the results suggest the simulated CIITA fragment displays the characteristics of an IDR with possible transient helix formation.

**Figure 3.**
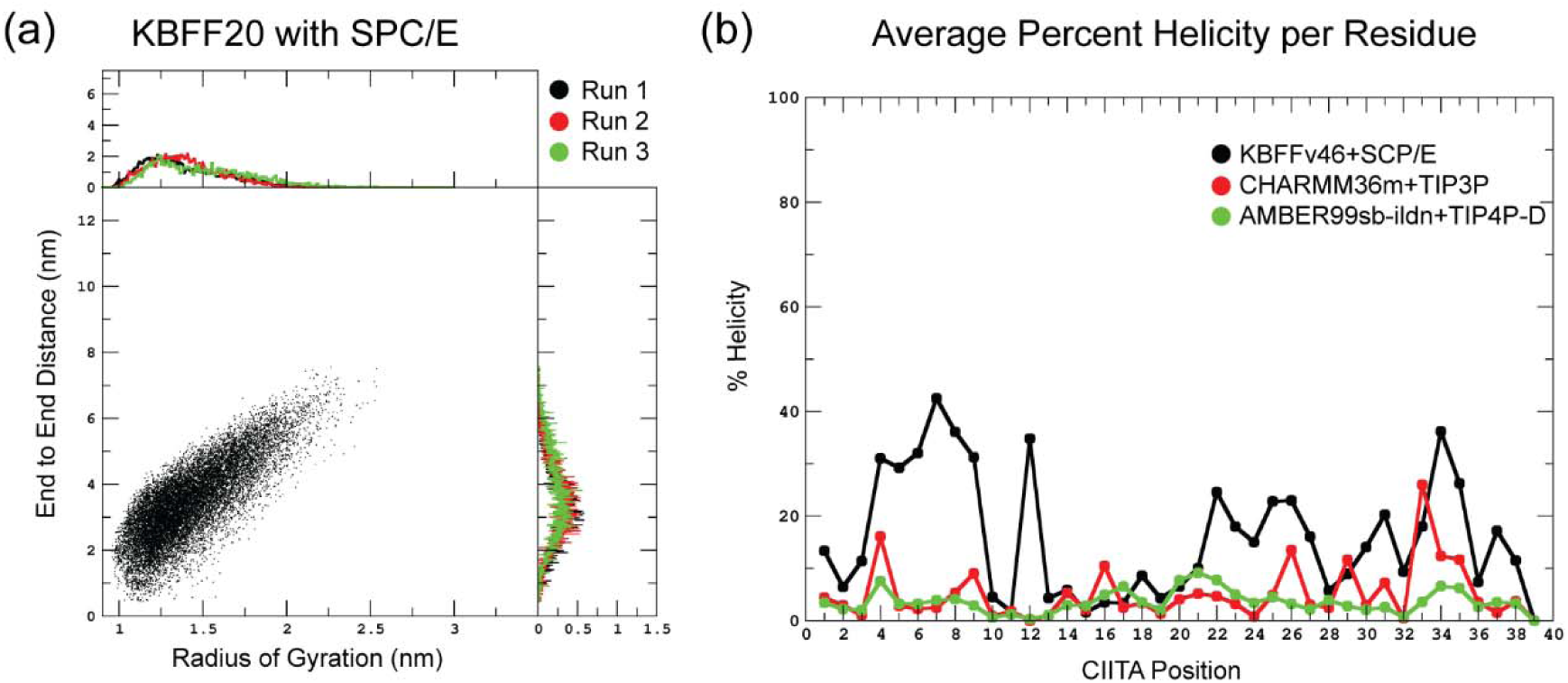
Simulated dynamics properties for the fragment of CIITA containing residues 56-94. (a) The radius of gyration and end-to-end probability distributions of CIITA 56-94 from three independent simulations (Runs 1-3) using the KBFF20 protein force field predict little-to-no persistent structure. Similar plots for simulations using the CHARMM and AMBER force fields are shown in Supplementary figure 3. (b) Helix residue probability distributions averaged over the three simulations for each of three different force fields; the numbering on the x axis corresponds to the peptide numbering ( *e.g*., position 1 on the plot is position 56 in the full-length protein).

Towards the end of our project, the predicted structure for full-length CIITA became available from AlphaFold2 (Supplementary figure 4).^51^ In agreement with sequence analyses, AlphaFold2 predicts that most of the activation domain lacks secondary or tertiary structure; the region tested in this study (between amino acids 61 and 92) was predicted to contain a small helix, albeit with low confidence.

### The intrinsically disordered activation domain of CIITA is functionally tuneable by single substitutions

Functionally, experimental evidence shows that CIITA binds to several protein complexes through its activation domain: BRG1^52^, CBP^19^, TAF9^53^, and SRC1^54^. These protein-protein interactions are critical for various aspects of transcriptional activation and could be sensitive to single amino acid changes in CIITA. Since the G58P and A94P substitutions partially impeded CIITA function,^55^ we reasoned that the regions near these two positions might be functionally sensitive to single amino acid changes. Thus, between positions 58 and 94, we chose seven CIITA positions from regions with high (I62, Y65, T70, T91) and low (N74, Q77, F78) probabilities for intrinsic disorder (Table 1). These regions contain binding sites for the four protein complexes listed above as well as an additional 10 short linear motifs (SLiMs)^56^ predicted to be involved in protein-protein interactions (Supplementary table 2).

To identify the roles of CIITA’s positions in tuning protein function, each position must be experimentally assessed with multiple amino acid substitutions. If >70% of substitutions maintain WT-like function for all measured parameters, the position is assigned to the “neutral” category.^57^ If its set of substitutions samples ≥50% of the accessible functional range, the position is assigned to the “rheostat” category;^2^ this behavior has been observed for numerous positions in globular proteins^1–3; 5; 6; 9; 38; 58; 59^. If >64% of substitutions abolish function, the position is assigned to the “toggle” category;^60^ this textbook substitution behavior is frequently observed at important, conserved positions in globular proteins. These three types of outcomes can be quantitatively identified using a modified histogram analysis that returns three scores (neutral, rheostat, and toggle).

Thus, we used an unbiased approach to generate at least 10-14 substitutions (including WT) at each targeted position in full-length CIITA.^38^ Next, we measured the effects of substitutions on CIITA’s overall *in vivo* function in regulating an MHC class II promoter controlling expression of a luciferase reporter gene (Supplementary figure 5). The functional range observed for the complete set of substitutions spanned ∼80-fold (Supplementary figure 6). All seven positions were non-neutral, and three positions (position 62, 65, and 77) met the threshold for being strong rheostat positions (rheostat scores >0.5). Notably, our experimental design revealed information that could not be gleaned from deletion studies. First, the *in vivo* transcriptional activation for Y65H (5% of WT activity) was lower than that of the previously-assessed Δ51-71 deletion variant (27% of WT activity)^55^, which demonstrates the surprising finding that single substitution can be more inhibitory than deleting a 20 amino acid region. Second, variants with enhanced activation (*e.g.,* 400% of WT for I62V), which expands the functional tuneable range, could not be detected in deletion studies.

### Changes in *in vivo* function can arise from changes in cellular concentration and/or transcriptional activation

The effects on CIITA *in vivo* function could arise from changes in either cellular concentration, or changes in events required for transcriptional activation, or both. Thus, we also measured the effects of amino acid substitutions on cellular protein concentration of full-length CIITA in HEK293 cells and used these values to calculate normalized transcriptional activation for each variant (Supplementary figure 5).

Cellular protein concentrations can be affected by changes in numerous processes, including transcription of the CIITA cDNA, mRNA translation, proteolytic susceptibility, and protein stability. Although any one (or more) of these processes could be altered by amino acid substitution, many of the 81 substituted CIITA variants had WT-like cellular concentrations (58 variants from one-way ANOVA; 48 variants from FDR analyses; Figure 4, Supplementary table 3). However, when effects were considered at individual positions, only position 91 was classified as neutral (RheoScale neutral score >0.7; Table 1). All other positions had a few variants with diminished concentration and, as such, were non-neutral. No positions met the threshold of a strong rheostat position (RheoScale rheostat score >0.5) (Figure 4; Table 1).

**Figure 4.**
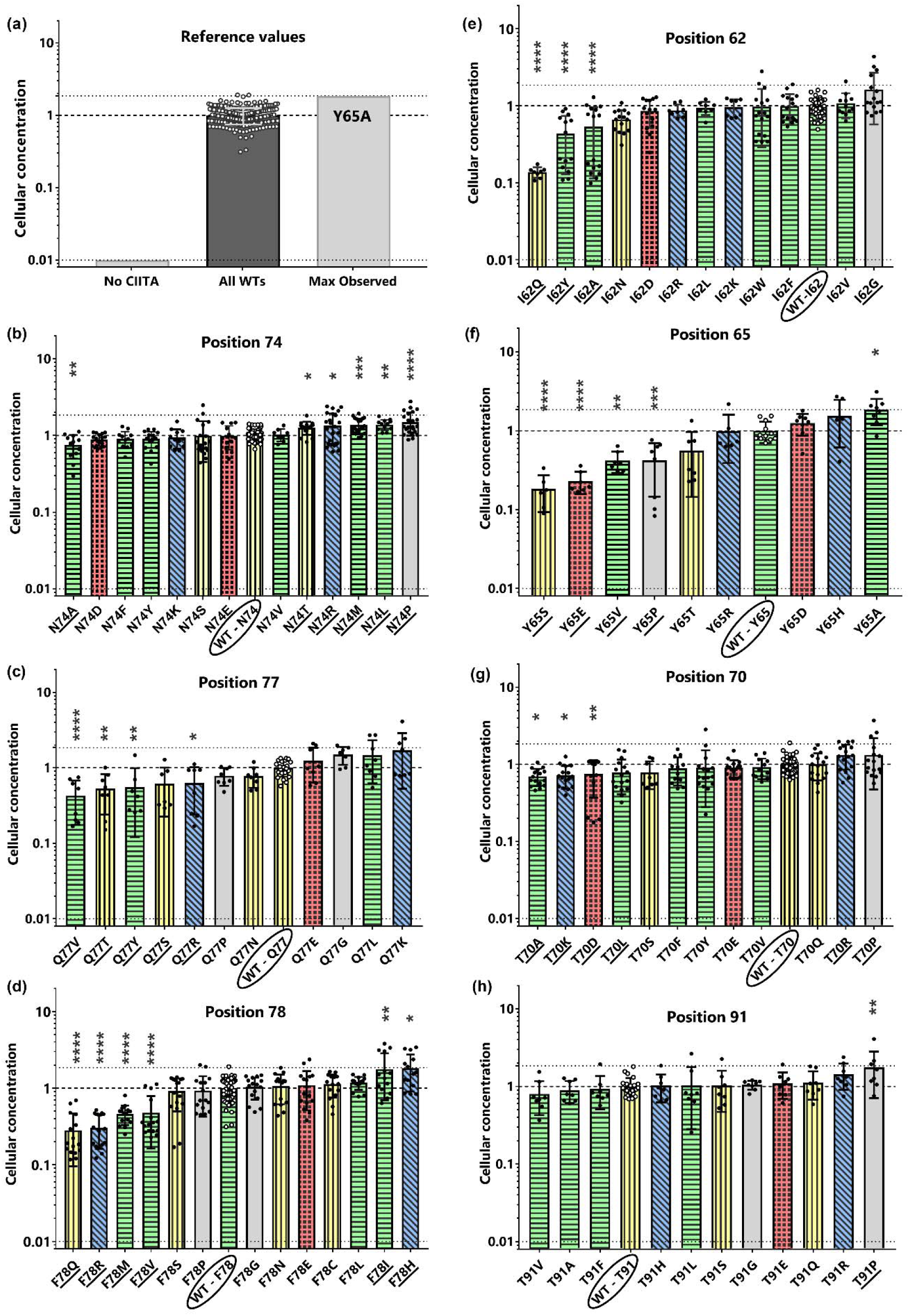
Cellular concentrations for CIITA variants. All data were normalized to a WT value that was set to 1. (a) Reference range for cellular concentrations: The background signal (“No CIITA”) was estimated as ∼1% of the WT CIITA concentration (dotted line at 0.01); the variant with the highest cellular concentration of CIITA was set as “Max Observed” (dotted line at 1.85). All values for the biological and technical replicates of WT are shown as dots; the average of these values was normalized to 1 and the standard error of means is shown with error bars. (b)-(h) The normalized cellular concentrations of CIITA variants are shown for positions with low (b) N74, (c) Q77 (d) F78 and high disorder probabilities (e) I62, (f) Y65, (g) T70, (h) T91. Individual measurements are shown as black dots, and the standard error of the means are indicated with error bars. The variant corresponding to WT is circled on the X-axis legend, and the normalized WT value of 1 is indicated with a dashed horizontal line; the WT samples shown on (b)-(h) were measured in parallel with each position’s variants. Bars are coloured according to amino acid type: acidic (D, E; red, checked), basic (R, H, K; blue, slanted lines), polar uncharged (S, T, N, C, Q; yellow, vertical lines), hydrophobic (M, F, W, Y, A, V, I, L; green, horizontal lines), and structure breakers (P, G; grey, solid). Substitution variants with protein levels statistically different from WT were identified with (i) one-way ANOVAs with Dunnett’s correction (****p <0.0001, ***p<0.001, **p<0.01, *p<0.1), and (ii) false discovery rate analyses (underlined variants on the X-axis legend).

For normalized transcriptional activation, a wide range of outcomes was also observed (Figure 5), with some variants diminishing and others enhancing this function (Supplementary table 3). The overall range spanned >30-fold. No positions were neutral; all had at least a few variants that altered transcriptional activation (Figure 5). Four of the seven amino acid positions (62, 65, 77, and 78) met the threshold for rheostat behaviour (rheostat score ≥0.5). Of these four positions, position 78 was not a rheostat for the overall *in vivo* functional assays because effects on transcriptional activation were compensated by effects on protein concentration. For these four positions, we hypothesize that substitutions alter protein-protein interactions, although they might also alter CIITA’s sub-cellular localization^61^.

**Figure 5.**
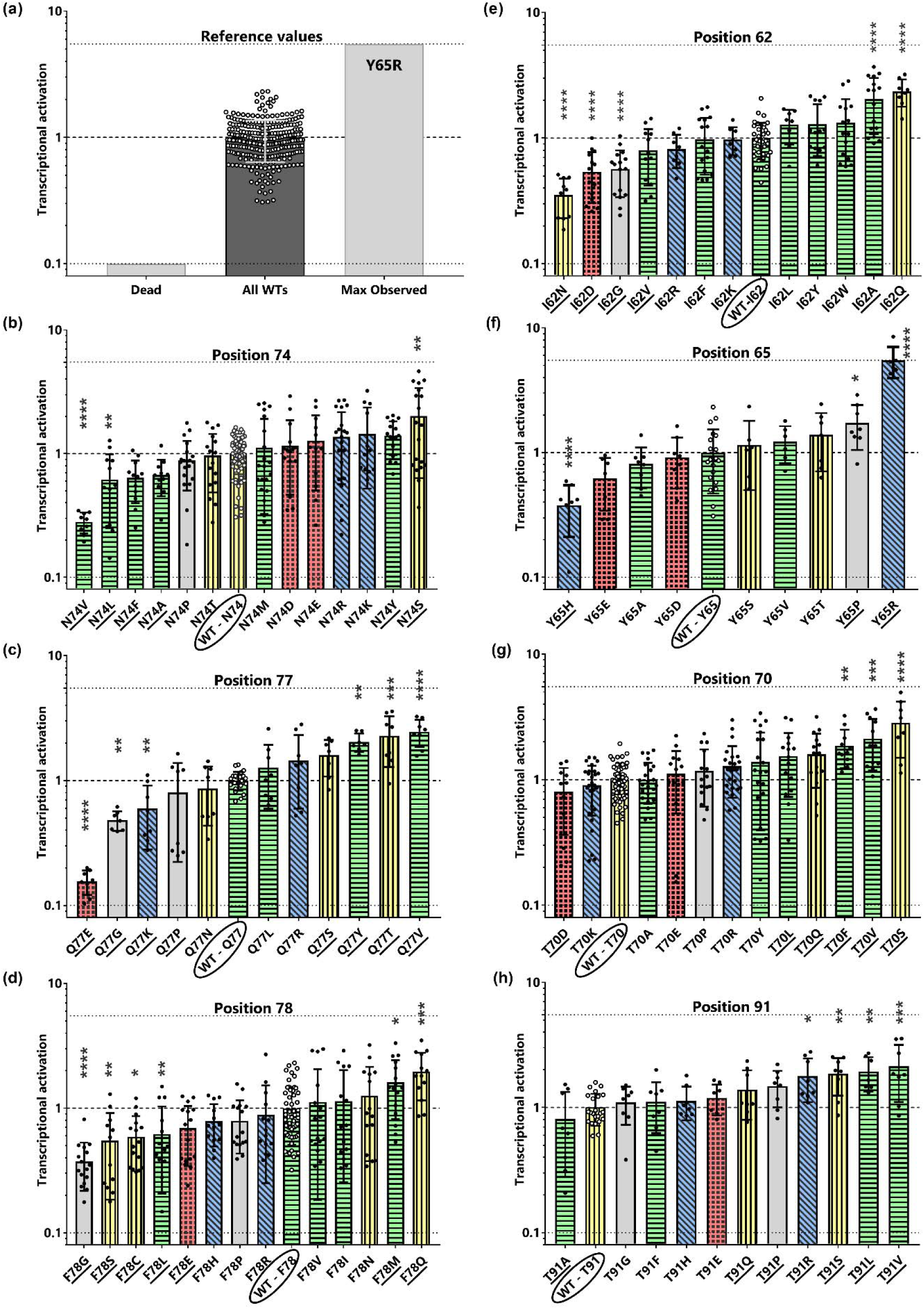
Normalized transcriptional activation for CIITA variants. All data were normalized to a WT value that was set to 1. (a) Reference range for transcriptional activation: At the lower end of the range, the transcriptional activation level for a hypothetical “Dead” CIITA was determined from a control sample that lacked CIITA (dotted line at 0.08; calculations in Supplementary table 7). The upper end of the range was defined by the variant with the greatest transcriptional activity, Y65R (“Max Observed”, dotted line at 5.5). All values for the biological and technical replicates of WT are shown with dots; the average of these values was normalized to 1 and the standard error of means is shown with error bars. (b)-(h) Normalized CIITA transcriptional activation levels are shown for each substitution variant at positions with low (b) N74, (c) Q77, (d) F78 and high disorder probabilities (e) I62, (f) Y65, (g) T70, (h) T91. Individual measurements are shown with black dots, and standard error of the means are indicated with error bars. The variant corresponding to WT is circled on the X-axis legend, and the normalized WT value of 1 is indicated with a dashed line; the WT samples shown on (b)-(h) were measured in parallel with each position’s variants. Bars are coloured according to amino acid type: acidic (D, E; red, checked), basic (R, H, K; blue, slanted lines), polar uncharged (S, T, N, C, Q; yellow, vertical lines), hydrophobic (M, F, W, Y, A, V, I, L; green, horizontal lines), and structure breakers (P, G; grey, solid). Variants with activity statistically different from WT were identified with (i) one-way ANOVAs with Dunnett’s correction (****p <0.0001, ***p<0.001, **p<0.01, *p<0.1), and (ii) false discovery rate analyses (underlined variants in the X-axis legend).

### The locations of these CIITA rheostat positions cannot be inferred from phylogenetic patterns of change

In two globular proteins, the set of positions that showed subfamily-specific evolutionary conservation were enriched with rheostat positions,^9^ and extremely nonconserved positions were neutral.^57^ Although disordered proteins were initially assumed to be highly non-conserved, increases in sequencing information have shown that disordered sequences have both conserved and non-conserved positions.^62^ To ascertain whether the rheostat/phylogeny correlation holds for CIITA, we used ConSurf analyses of multiple sequence alignments (MSAs).^63^ For each amino acid position (MSA column), ConSurf uses a tripartite function to compute a conservation score, which is then combined with information from the protein’s phylogenetic tree. This generates final ConSurf scores that differentiate positions with whole-family conservation from those with subfamily conservation and those with nonconservation.

For ConSurf analyses, input MSAs for CIITA were generated by two independent approaches (see Methods). Both MSAs yielded similar conservation trends (Supplementary table 4), which suggests that results were not biased by MSA construction. The CIITA N-terminus contains alternating regions of high and low conservation, with no apparent correlation with disorder probabilities (Supplementary figure 7). For the seven tested positions, no correlation was observed between ConSurf scores and substitution outcomes. However, CIITA is only present in vertebrates; the other studies correlating substitution outcomes with evolutionary scores were for pyruvate kinase, which are present in all domains of life^7^ and LacI/GalR homologs, which are present in almost all bacteria^58; 64^. Both of these families are much older than CIITA, providing more opportunities to explore sequence space. As such, MSA-based predictions about CIITA positions (and perhaps other taxonomically-restricted proteins) may not be meaningful.

### Tuneability from single substitutions in the activation domain of CIITA are not explained by amino acid side chain chemistries

For rheostat positions in globular proteins, substitution outcomes were not well explained by similarities in physicochemical properties of amino acid side chains.^2; 5; 6; 58^ Thus, we next assessed whether chemically-similar side chains had similar outcomes on CIITA’s transcriptional activation. When the rank order of substitution outcomes was colour-coded according to amino acid side chain chemistry, we again did not observe any obvious relationship (Figures 4 and 5, Supplementary figure 6, bar colours). For example, the transcriptional activation of I62N was 35% of WT, whereas the “similar” I62Q substitution increased transcriptional activation to 200%.

Next, we considered whether the observed changes in CIITA’s transcriptional activation were consistent with the acidic exposure model. Activation domains typically contain hydrophobic motifs flanked by acidic residues, and studies using engineered transcriptional activator proteins showed that the balance of these two features can be altered to vary transcriptional activation.^13–16^ In the current study, although we did not mutate acidic residues in the WT sequence, our unbiased substitution strategy did introduce at least one acidic residue into most of the chosen positions. All of the acidic substitutions either reduced or maintained transcriptional activation, consistent with the acidic exposure model. However, substitutions with non-acidic amino acids often showed larger effects on CIITA function than did acidic side chains. Furthermore, most of the aromatic substitutions maintained WT function; only T70F showed the enhanced function predicted by the acidic exposure model.

### Changes in transcriptional activation may be explained by altered structural features

Since side chain chemistries do not explain changes in functional outcomes, we next assessed other structural features. To this end, we correlated the experimentally-measured parameters with: (i) side chain size, as calculated from G-X-G tripeptides;^65^ (ii) amino acid helical propensities, which were expressed as free energies (*i.e.,* lower values indicate higher helical propensity);^66^ and (iii) changes in disorder probabilities arising from substitution (Supplementary figure 8; control computations for this approach are described in Supplementary Figure 9). When the measured cellular concentrations of each CIITA variant were compared to the three structural features, no meaningful correlations were observed (Supplementary figures 10-12). In contrast, when transcriptional activation was assessed (Supplementary figures 13-15), three positions showed intriguing correlations (Figure 6).

**Figure 6.**
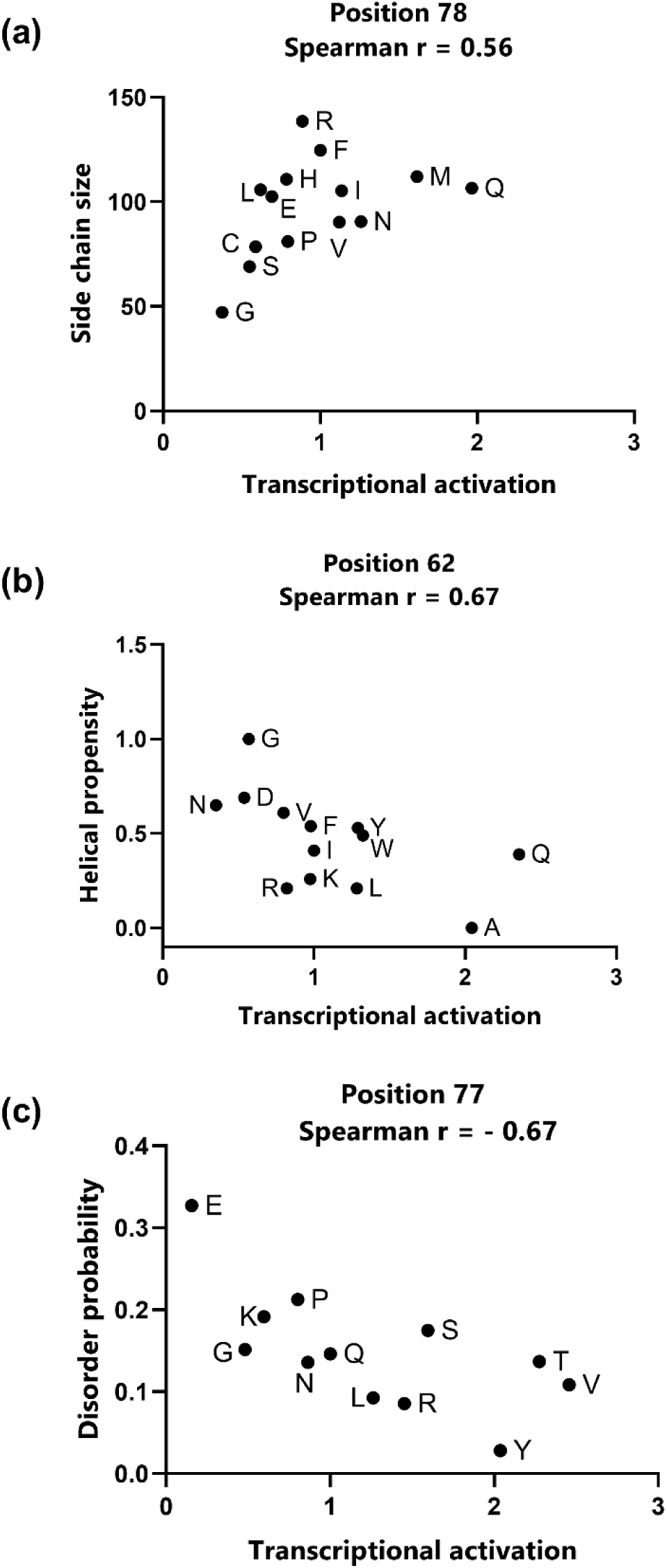
Correlation of transcriptional activation with structural features. Of the 28 correlations of transcriptional activation with structural features, three showed Spearman coefficients >0.5. (a) At position 78, transcriptional activation positively correlated with the amino acid side chain size obtained from the relative, solvent-accessible amino acid side chain surface area (Å ^2^) in a G-X-G tripeptide, where X is the amino acid. (b) At position 62, transcriptional activation negatively correlated with amino acid helical propensities obtained from host-guest peptide studies (reported in units of kcal/mol); lower values correspond to higher helical propensities. (c) At position 77, transcriptional activation positively correlated with disorder probabilities predicted by PONDR. Correlations for all other positions/structural features are shown in Supplementary figures 10-15.

Rheostat position 78 exhibited a relationship between transcriptional activation and amino acid size (Spearman r = 0.56), with an increase in size corresponding to increased transcriptional activation. Rheostat position 62 exhibited a relationship between transcriptional activation and helical propensities (Spearman r = -0.68), which suggests that an increase in helix formation favours transcriptional activation (Figure 6). Rheostat position 77 showed a negative correlation between transcriptional activation and PONDR’s disorder probability (Spearman r = -0.67), indicating that lower disorder probabilities led to higher transcriptional activation (Figure 6, Supplementary figure 15b); the same trend occurred when disorder probabilities were taken from an alternative disorder calculator, metapredictV2 (Supplementary figure 15c). Furthermore, the correlation between helical propensity and transcriptional activation was high for position 77 (Spearman r = 0.60) when the helix breakers G and P were excluded (Supplementary figure 14b).

The correlations for positions 62 and 77 are especially intriguing in light of the MD simulations, which predicted that transient helices occur in this region of the activation domain (average per residue helicity: KBFF20 = 16%, Charmm = 5%, and Amber = 4%; Figure 3). Indeed, in the KBFF20 simulations of WT CIITA, the average helicity at position 62 was as high as 43% and position 77 is in a region simulated to have ∼20% average helicity. Thus, substitutions at these positions might tune transcriptional activation by altering a coil-to-helix equilibrium important for transcriptional activation.

## DISCUSSION

Our results show that single substitutions in the intrinsically disordered activation domain of CIITA can alter function: This limited set of substitutions, created by semiSM of just seven of the 1130 CIITA positions, sampled an 80-fold range in overall *in vivo* function and a >30-fold range in normalized transcriptional activation (Figure 7). Because substitutions sampled most of the range bounded by WT and the “no activation” control, these changes also met our definition of “tuning”. In addition, comparable numbers of diminishing and enhancing substitutions were identified (Supplementary Table 3). As such, the CIITA activation domain was more tuneable than the activation domains of p53^20^ and PPAR□^21^.

**Figure 7.**
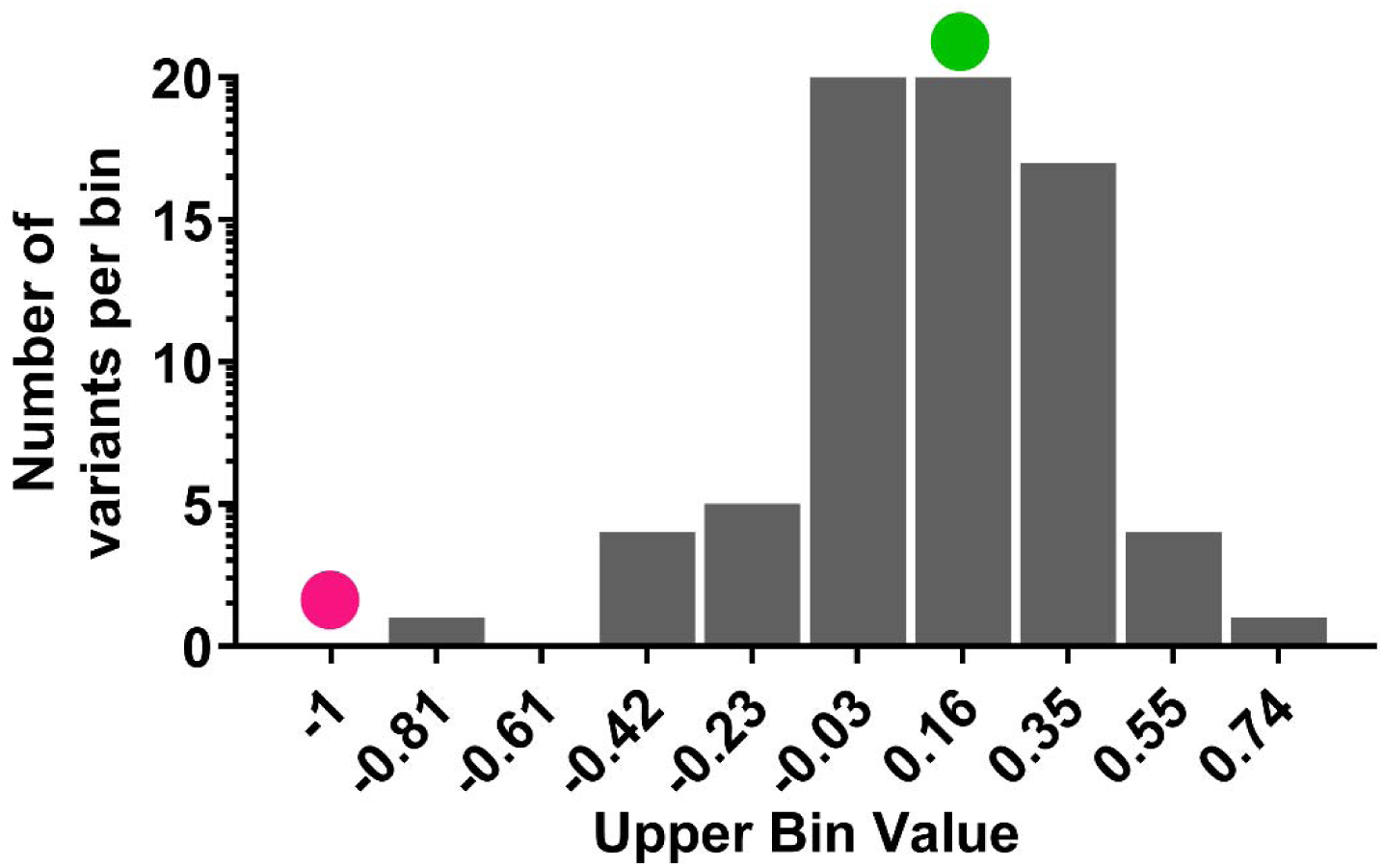
Overall tunability of CIITA’s transcriptional activation. A histogram of the values observed for the set of 82 variants at seven CIITA positions shows that transcriptional activation is fully tuneable by single substitutions. Numbers on the X-axis correspond to log(upper limits) for transcriptional activation. The overall range of observed change is >30-fold. The green dot indicates the bin containing WT. The magenta dot at the far left indicates the level of a hypothetical “dead” variant (see Supplementary table 7 for these calculations).

Furthermore, substitution outcomes did not follow the patterns predicted by the acidic exposure model. According to this model, increasing the fraction of acidic residues should enhance activation by skewing the equilibrium towards the exposed conformation. In contrast, increasing the net charge/hydrophobicity should diminish activation by skewing the equilibrium towards the collapsed conformation. A tempting extrapolation of this model is to assume that substituting amino acid side chains with other chemistries will have no effect on transcriptional activation and, thus, rheostat positions would not exist. However, this was not observed in CIITA: (i) Substituting a variety of side chain chemistries altered transcriptional activation over the tuneable range, (ii) several substitutions had greater impact than changes that altered the acidic/hydrophobic balance, and (iii) rheostat positions did exist.

The simplest explanation for the discrepancy between the results observed for CIITA and those expected from the acidic exposure model is that different types of activation domains exist, each associated with different sensitivities to substitution. Full-length CIITA and full-length p53/PPARɣ clearly exhibit different sensitivities to single substitutions. This demonstrates that substitution outcomes should *not* be generalized across all activation domains and allows for the possibility that the acidic/hydrophobic balance controls function in some proteins, whereas rheostat positions are present in others. As such, CIITA could either be an exception among proteins containing IDR activation domains or could exemplify a new paradigm for substitution sensitivity.

Alternatively, none of the existing studies rule out the possibility that the two means of functional tuning co-exist. First, the presence (or absence) of rheostat positions requires semiSM/SSM experimental design to interrogate the roles of individual positions. In contrast, the “rational mutagenesis” design used sets of substitutions in 8-amino acid windows to interrogate how changing the balance among acidic and hydrophobic residues altered function; this approach would not detect rheostat positions. Second, the acidic exposure studies were carried out with fragments of activation domains linked to a non-native DNA binding domain; the presence of rheostat positions might require the context of full-length protein, as was tested here for CIITA. Third, the current work with CIITA did not exhaustively test the whole activation domain, and – although various substitutions changed the charge or hydrophobicity at individual positions – the single substitutions only caused minor perturbations to the overall charge balance. Thus, future studies will be required to determine whether rheostat positions and acidic exposure behaviour co-exist or are mutually exclusive.

We must also determine the molecular mechanisms by which substitutions at IDR rheostat positions exert their effects on transcriptional activation. Since the existence of rheostat positions is not fold-dependent, perhaps the biophysical changes associated with their substitutions are similar in both IDRs and globular proteins. In the globular protein LacI, the modulating effects of substitutions were well-described by perturbations to an all-atom dynamics model comprising an asymmetric, weighted network of balls (nodes/atoms) and springs (edges/bonds);^8^ this computation has also been successfully correlated with single substitution outcomes in other proteins (*e.g.,* ^4^). Notably, such a model does *not* require the presence of ordered secondary structure. Instead, any atom added or deleted from the protein has the potential to perturb the asymmetric weighted network. CIITA function requires binding of its activation domain to partner proteins and thus CIITA participates in a structural network. As such, single amino acid substitutions could have the same potential to alter functionally-relevant dynamics.

In conclusion, substitution sensitivities cannot be generalized among IDR activation domains. Furthermore, aside from the p53 and PPARγ studies, IDRs have been largely passed over in SSM studies; of the 161 proteins with IDRs in MaveDB^67^, only their globular domains have been mutated as of this writing. Thus, future SSM/semiSM studies of full-length transcription factors are required to illuminate mechanisms of functional tuning in IDRs.

## MATERIAL AND METHODS

### Sequence analyses of full-length CIITA

The human CIITA sequence exists as multiple isoforms that arise from alternative promoter usage and mRNA splicing (44). The full-length human CIITA sequence for isoform 1 (Uniprot P22076.1), comprising 1130 amino acids, was used in all analyses.

#### Intrinsic disorder prediction

The CIITA primary amino acid sequence was analysed with the Predictor of Natural Disorder Region (PONDR) software using its VL-XT subroutine to obtain disorder probabilities ^68^ (Supplementary figure 2a, Supplementary table 1). We also computationally edited the WT CIITA sequence to create full-length sequences corresponding to all possible single substitutions at the seven chosen positions (133 sequences) and all possible substitutions at a different position with high disorder probability and repeated the PONDR analyses (Supplementary figures 8 and 9). As alternative computational approaches, we used IUPred3^44^, fIDPnn^43^, and metapredictV2^42^ using their default parameters (Supplementary figure 2b; Supplementary table 1; Supplementary figure 9) as suggested by RIDAO^69^.

#### Conservation scores

To compute evolutionary information for CIITA positions, we used the human CIITA sequence in an automated ConSurf computation using default parameters: This webserver first used an HMMER-based^70^ search to build a multiple sequence alignment (MSA; named “CIITA_ConSurf_default_MSA”, Supplementary data) using a MAFFT-LINS-i alignment ^71^, which was then subjected to phylogenetic/conservation analysis. This MSA was limited to 150 sequences, which might lead to biased ConSurf scores due to insufficient sampling; larger MSAs could not be constructed using the default ConSurf parameters because the CIITA sequence is so long that the process timed out.

Thus, we also created a larger CIITA MSA by manual curation: The CIITA primary amino acid sequence was used as a BLAST query against the nr-database in the NCBI server ^72^; results were filtered using 80 to 100% query coverage to yield 1102 unique sequences. These sequences were aligned using clustalOmega ^73^. This initial MSA (i) was manually curated to remove sequences with insertions >100 amino acids or >100 sequential gaps, leaving 1017 sequences, and (ii) a python script^74^ (https://github.com/ncbi/FixJ_KdpE/tree/main/clusters/cull_MSA.py) was used to exclude other long and short sequences: To filter out sequences with long inserts (which can disrupt alignments of other sequences), sequences were searched to identify windows of 8-positions for which <5% of the other sequences were occupied with an amino acid. To filter out sequences that were too short (which could bias the gap penalty calculations for the region and slow down ConSurf calculations), sequences were searched to identify windows of 10-positions for which >90% of the other sequences contained amino acids. The remaining 603 sequences were re-aligned in clustalOmega to produce the MSA named “CIITA_manual_MSA” (Supplementary data).

CIITA_manual_MSA was input into ConSurf^63^ to identify position-wise conservation scores. However, ConSurf was unable to analyze this MSA, perhaps again because it was too large. Thus, from the CIITA_manual_MSA, we randomly extracted five subsets for each of 250, 300, 350, 400, 450 sequences using python scripts (https://github.com/ShwethaSreenivasan/CIITA_Swint-Kruse_Lab_2023). Except for one of the 450-sequence MSA subsamples, all ConSurf analyses were successful; position raw scores were in good agreement with each other for subsets ≥300 sequences (Spearman, r = 0.96-0.99, Supplementary figure 16; ConSurf’s transformation of raw scores into 9 categories is further described in Supplementary table 4). Thus, we used the 19 largest subsets to calculate an average normalized ConSurf score for each position (Supplementary table 3).

ConSurf results for the manual and default MSAs were similar (Spearman, r = 0.85, Supplementary figure 16i). ConSurf scores of the experimentally-assessed positions ranged from 2 (low) to 9 (high) (Table 1, Supplementary table 4). ConSurf scores from 3-8 were previously associated with a high prevalence of rheostat^9^ positions in soluble globular proteins. Also in soluble globular proteins, positions which showed the least pattern of change within an MSA (*i.e.,* changes did not follow any of several phylogenetic or co-evolutionary patterns) were associated with a prevalence of neutral positions;^57^ such positions usually have ConSurf scores of 1-2.^9^ However, for CIITA, comparison of ConSurf and PONDR scores showed no correlation (Spearman, r = 0.16-0.17, Supplementary figure 7).

#### Predictions for short linear motifs (SLiMs)

SLiMs are 3-10 amino acid long regions found especially in intrinsically disordered sequences that contain putative protein-protein interaction sites. Using the full-length CIITA, analyses with Eukaryotic Linear Motif^56^ predicted 43 SLiMs in the activation domain (Supplementary table 2).

### Spectroscopic characterization of CIITA protein fragment 1-210

Based on the PONDR analysis (Figure 1), a fragment of the CIITA protein containing amino acids 1-210 with an N-terminal HIS6-tag was expressed utilizing a pET28b expression plasmid. This region spans all seven positions chosen for experimental substitution studies. Briefly, *E. coli* Rosetta cells (Millipore Sigma, catalogue #70954) were transformed with pET28b-CIITA 1-210 and a single colony was used to inoculate 5 mL LB medium with 50

μg/mL kanamycin and grown overnight at 37°C. From that overnight culture, 100 μL were inoculated in 1 L of LB medium supplied with kanamycin (50 μg/mL) and grown to OD of 0.6. The temperature was reduced to 20°C and 0.25 mM IPTG was added. After 22 h growth, cells were harvested by centrifugation and frozen at -80 °C until purification.

For purification of the CIITA protein fragment, cells were resuspended in binding buffer (50 mM Tris-HCl pH 8.0, 300 mM NaCl, 5 mM imidazole, and 1 mM PMSF) and lysed by sonication. Cleared supernatant was mixed with Ni-NTA agarose (GE Healthcare, MO), bound overnight with mixing, and washed with binding buffer. Protein was eluted with 0.5 M imidazole in the same buffer. Additional purification was performed using AKTA FPLC system on the Superose 12 column (GE Healthcare, MO) equilibrated with 50 mM sodium phosphate buffer (pH 8.0).

CIITA fragment 1-210 concentration was determined by absorbance measurements at 280 nm using molecular extinction coefficient of 14000 mol^-1^, which was estimated from the one tryptophan and seven tyrosine residues in this sequence. Circular dichroic (CD) spectra of the CIITA 1-210 fragment were obtained using a Jasco-720 spectropolarimeter (Japan Spectroscopic, Tokyo, Japan). Samples containing 4 μM of protein in 50 mM sodium phosphate buffer pH 8.0 were placed in 1-mm optical path cuvette at room temperature in the absence or presence of 1 M trimethylamine N-oxide hydrate (TMAO) (Hampton Research, CA). Ellipticity data were collected for 200-260 nm with a 1 nm step. Data from 100-150 scans were averaged and the background was subtracted to generate the final spectra.

### Molecular dynamics simulations of CIITA fragment 56-95

Based on the PONDR analysis (Figure 1), a truncated sequence of CIITA encompassing residues L56 to A95 (39 residues total) was selected for molecular dynamics simulations (MD). This region spans all seven positions chosen for experimental substitution studies. First, the coordinates for a blocked (N-acetylated and C-N methylated) sequence in a fully extended conformation were generated using the Avogadro software ^75^. Then, three random initial configurations were selected from preliminary 100 ns implicit solvent simulations using a simple dielectric constant of 80. Finally, explicit solvent simulations were performed starting from the three different initial conformations.

All simulations were performed with the GROMACS package version 2016.4 ^76^. Simulations of each configuration were performed with three different force fields: KBFF20 ^77^ with SPC/E water model ^78^; CHARM36m ^79^ with TIP3P water model ^80^; and AMBER99sb-ildn ^81^ with TIP4P-D water model ^82^. Each configuration of the CIITA fragment was solvated in a rhombic dodecahedral box with an image distance of 7.5 nm using periodic boundary conditions. The systems contained approximately 9400 water molecules, after neutralization with 12 Na^+^ ions and the addition of 25 more Na^+^ and 25 Cl^−^ ions to achieve a physiological (0.15 M) salt concentration.

The equations of motion were numerically integrated using the Verlet leapfrog algorithm ^83^ with a time step of 2 fs. The short-range interactions were calculated using a nonbonded pair list with a single cut-off of 1 nm with a Verlet list update. PME was applied for the long-range electrostatic interactions ^84^. Nose-Hoover temperature coupling was used to maintain a target temperature of 300 K ^85^, and Parrinello−Rahman pressure coupling was used to maintain a target pressure of 1 bar ^86^. All peptide bond lengths were constrained using the LINCS algorithm ^87^, while all solvent bonds were constrained using SETTLE ^88^. After minimization using the steepest descents method, a 1 ns equilibration was performed using Berendsen temperature and pressure coupling at 300 K and 1 bar ^89^. Production simulations of 5μs were performed for each initial configuration and each force field.

### Generation of variants in full-length CIITA

The plasmid pUNO1-hCIITA (InVivogen, Inc., San Diego, CA; catalog code pUno1-ciita), which encodes the CIITA cDNA, was used to generate CIITA variants. For each targeted amino acid position, 9-13 unique mutations were generated by random, site-directed mutagenesis using the QuikChange II XL kit (Agilent Technologies, Inc., Santa Clara, CA). For each targeted codon, four sets of primers were synthesized (Integrated DNA Technologies, Inc. (Coralville, IA) with each set substituting one of the random codons ANN, CNN, GNN, or TNN for the target codon. Plasmids purified from individual clones were sequenced to identify mutations in the CIITA cDNA (ACGT, Wheeling, IL). All plasmid isolations used the ZymoPURE Plasmid Miniprep Kit (Zymo Research Corp., Irvine, CA). Unless otherwise stated, the manufacturer’s protocols were followed for all procedures.

### Transfection and Luciferase Assays

HEK293 cells (American Type Culture Collection, Manassas, VA; catalog number CRL-1573) do not express the MHC class II genes, making this cell line suitable for reporter assays of CIITA transcriptional activation. To that end, HEK293 cells were cultured in complete growth medium consisting of Dulbecco’s Modified Eagle Medium with 4.5 g/L glucose, supplemented with 10% fetal bovine serum, 100 µg/mL penicillin, and 100 U/mL streptomycin. Cells were maintained at 37 °C, under a humidified atmosphere containing 5% CO_2_. Growth was monitored daily, and the cells were split once they reached 80-90% confluence. Medium was replaced every two to three days, as required. Cells were used for a maximum of 15 passages. Subcultures for transfection were prepared in 24-well plates with 1.5×10^5^ cells and 1 mL complete medium per well. Subcultures were incubated overnight prior to transfection.

All plasmid transfections were performed in quadruplicate. Each transfected well received 100 µL Opti-MEM (Thermo Fisher Scientific, Waltham, MA), 100 ng of reporter plasmid pDRA-Gluc (the *Gaussia* luciferase gene under the control of the HLA-DRA promoter, responsive to CIITA)^90^, 100 ng pSV40-Cluc (expressing *Cyrpridina* luciferase, which served as a transfection efficiency control), 800 ng pUNO1-hCIITA (WT or variant), and 2 µL TurboFect transfection reagent (Thermo Fisher Scientific). Culture media was changed four hours post transfection. Cells were incubated overnight before performing the luciferase assay and harvesting cells for western blot analysis.

Luciferase assays were carried out in 24-well culture plates. For each plate, one transfection mixture was prepared with WT pUNO1-hCIITA as a positive control for activation of expression from the pDRA-GLuc plasmid and one transfection mixture was prepared with pCMV-LacZ as a negative control. Luciferase assays were performed on aliquots of cleared media harvested from transfected cells using the Pierce *Gaussia* and *Cypridina* luciferase flash assay kits (catalog numbers 16159 and 16169 respectively; Thermo Fisher Scientific) and a GloMax 20/20 luminometer (Promega) per manufacturer’s instructions.

### Cellular concentrations of CIITA variant proteins

To assess the relative *in vivo* concentrations of CIITA variant proteins, cells were harvested and lysed in western loading buffer as described. Proteins were separated on precast 4-15% gradient polyacrylamide gels (Bio-Rad, Hercules, CA) and transferred to PVDF membranes (Bio-Rad, Hercules, CA). Membranes were blocked in 1X TBST and 1X fish gelatin (Biotium, Fremont, CA) and blotted with E-12 mouse monoclonal anti-human CIITA (catalog number sc-376174, Santa Cruz Biotechnology, Inc., Santa Cruz, CA) at a dilution of 1:100 and 9F3 rabbit monoclonal anti-human β-tubulin from (catalog number 2128, Cell Signaling Technology, Inc., Danvers, MA) at a dilution of 1:5000, incubating overnight at 4°C. After washing, secondary blotting was carried out in the same buffer with donkey anti-mouse IgG H+L Alexa Fluor 488 at a dilution of 1:2000 and donkey anti-rabbit IgG H+L Alexa Fluor 647 at a dilution of 1:5000 (Catalog #150079, Abcam, PLC, Cambridge, UK). Membranes were imaged using a GE Typhoon FLA9500 biomolecular imager in fluorescence mode. Two images were obtained for each membrane, using the 473 nm and the 635 nm excitation wavelengths. Quantification relative concentrations of protein in the CIITA and β-tubulin bands was accomplished using ImageJ version 1.52a (Rasband, W.S., U. S. National Institutes of Health, Bethesda, MD).

### Analyses of experimental data

Four experimental measurements were made for each CIITA WT and variant sample (Supplementary Figure 5): (i) *Transcriptional activation by CIITA* was measured using the *Gaussia* luciferase activity from the reporter plasmid (pDRA-GLuc) as a proxy. (ii) Variations in *transfection efficiency* were controlled by measuring the *Cypridina* luciferase signal intensity from the transfection control plasmid (pSV40-CLuc). (iii) *Cellular CIITA protein concentration* was determined from the intensity of the WT or variant CIITA western blot bands. (iv) Experimental variation in *cell density* in the western blot was controlled by the intensity of the β-tubulin band. For each variant, two or more technical replicates of four or more biological replicates were assayed. All measurements were then normalized to the value for a WT CIITA measured in parallel, to control for day-to-day technical variation. The four normalized, independent measurements were used to determine the different functional outcomes available in https://github.com/ShwethaSreenivasan/CIITA_Swint-Kruse_Lab_2023.

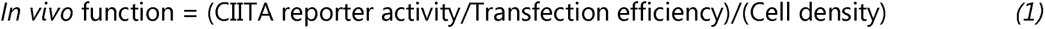

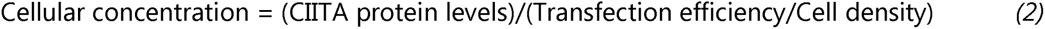

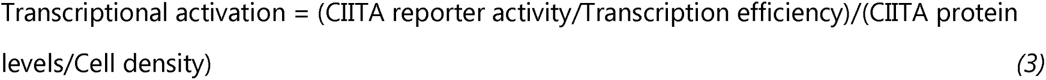

To determine which variants significantly differed from WT, statistical analyses were carried out using GraphPad Prism version 7.05.237. D’Agostino-Pearson tests showed that the replicates for several variants followed a log normal distribution, rather than a normal distribution. Hence, all values were transformed to log10 prior to statistical tests. One-way ANOVA was performed with Dunnett’s correction for multiple comparison of each substitution at a position to the value collected for WT of that position’s data set; statistically significant variants were identified (P < 0.0001, significant). Similar analyses were performed using correction for multiple comparisons by controlling for False Discovery Rate (FDR) analysis using the Two-stage step-up method of Benjamini, Krieger, and Yekutieli to identify “discoveries” significantly different from the WT. All parameters were set to Prism’s default values. Complete reports of the statistical analyses are included in Supplementary tables 5a-u and 6a-u.

### Assigning overall substitution outcomes to each CIITA position

For each position, a set of 10-14 variants (including WT) was used determine overall substitution outcomes. To that end, we used the RheoScale calculator, which uses a modified histogram analysis to quantify how a set of variants samples the empirically observed range of possible functional outcomes.^38^ The full range is determined by the full dataset – here, 81 variants and WT CIITA – in addition to the negative control that determines the value for “no detectable activity” (or “no detectable protein”). Using this range, the Rheoscale calculator reports three scores for each position: (1) the fraction of variants equivalent to WT (neutral score); (2) the fraction of variants that showed no activity (toggle score), and (3) the fraction of the accessible functional range that is sampled by the set of variants (rheostat score). For the latter, the observed range is divided into histogram bins; bins occupied by at least one variant are counted as filled; to account for potential error in the WT measurement, bins farther from WT are given higher weights than the WT-containing and adjacent bins. The number of bins used to analyse each of the measured outcomes were set by the default recommendations of the RheoScale calculator: Cellular concentration - 11 bins; transcriptional activation - 9 bins; combined effects - 6 bins.

## DATA AVAILABILITY

The number of replicates, averages and standard deviations for the values shown in Figures 4 and 5 are available in https://github.com/ShwethaSreenivasan/CIITA_Swint-Kruse_Lab_2023.

## SUPPLEMENTARY DATA

Supplementary Data are available at Protein Science.

## ACKNOWLEDGEMENTS

We thank Kristen Schwingen, Ibtihal Alghusen, and Halya Fedosyuk (KUMC) for assistance with initial experiments and Braelyn Page (KUMC) and Dr. Sarah Bondos (Texas A&M Health Science Center) for discussions on the manuscript. We also thank Jordan Baker (KUMC, Department of Biostatistics and Data Science) for providing suggestions for data analyses.

