## Supplementary for "The intrinsically disordered transcriptional activation domain of CIITA is functionally tuneable by single substitutions: An exception or a new paradigm?"

Average values and standard errors of the mean for each variant of CIITA in Figures 3-6 of the main text are posted as an Excel™ file on <https://github.com/ShwethaSreenivasan/CIITA_Swint-Kruse_Lab_2023>.

**Supplementary figure 1. Functional tuning of the Hif1α-derived activation domain.** Data for 15 singly-substituted variants of an Hif1α activation domain fragment (positions 781-826) were taken from Staller *et al.*^1^ and used to assess the overall turning associated with single substitutions. The numbers on the X-axis correspond to log(upper limits) for values of transcriptional activation. The green dot indicates the bin containing the parent Hif1α fragment. The magenta dot indicates basal transcriptional activation in the absence of Hif1α.

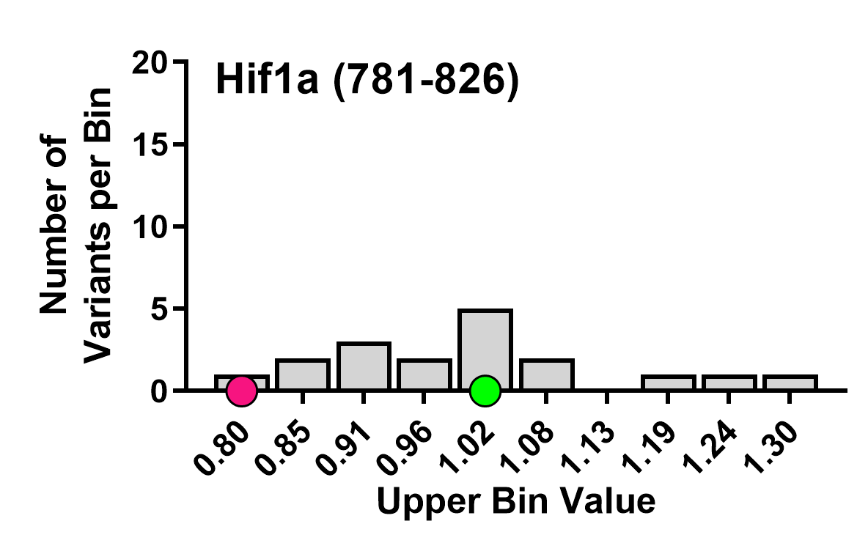

**Supplementary figure 2. Disorder probabilities for CIITA. (a)** Disorder probabilities predicted by PONDR (VL-XT subroutine) for full length-CIITA (positions 1-1130). Of the seven experimentally tested positions, those in regions with high disorder probabilities (>0.5) are indicated by vertical red, solid lines (62, 65, 70, 91); positions in regions with low disorder probabilities (<0.5) are indicated with vertical blue dashed lines. Figure 1 in the main text shows an expanded version of this plot for the N-terminal activation domain. **(b)** Disorder probabilities predicted by four different predictors^2-5^ for full length-CIITA (positions 1-1130). The seven tested positions are again indicated by solid red and dashed blue vertical lines; values for these positions are in Supplementary table 1.

**(a)**

**(b)**

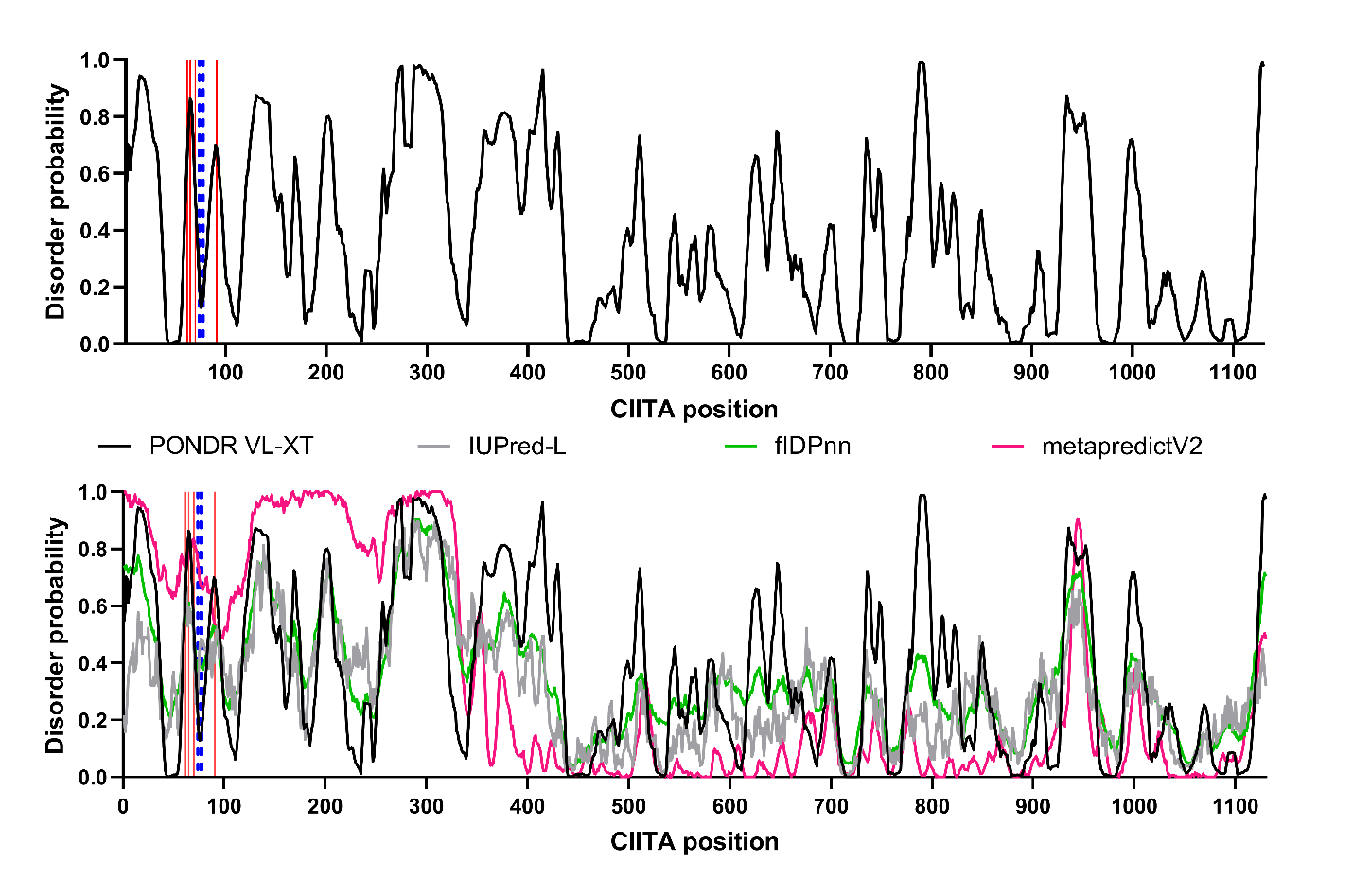

**Supplementary table 1.** Disorder probabilities for the seven chosen positions using alternative algorithms ^2-5^.

| **CIITA position** | **PONDR VL-XT** | **IUPred-L** | **fIDPnn** | **metapredictV2** |
| --- | --- | --- | --- | --- |
| 62 | 0.66 | 0.68 | 0.59 | 0.76 |
| 65 | 0.86 | 0.60 | 0.61 | 0.77 |
| 70 | 0.53 | 0.46 | 0.50 | 0.83 |
| 74 | 0.22 | 0.30 | 0.37 | 0.75 |
| 77 | 0.15 | 0.48 | 0.39 | 0.67 |
| 78 | 0.16 | 0.48 | 0.38 | 0.67 |
| 91 | 0.69 | 0.43 | 0.52 | 0.54 |

**Supplementary figure 3.** **MD simulations show that CIITA fragment 56-95 is not compact.** The plots below show the end-to-end distances and radius of gyration distributions for simulations starting from three different initial conformations (Runs 1-3) using each of three different protein force fields.

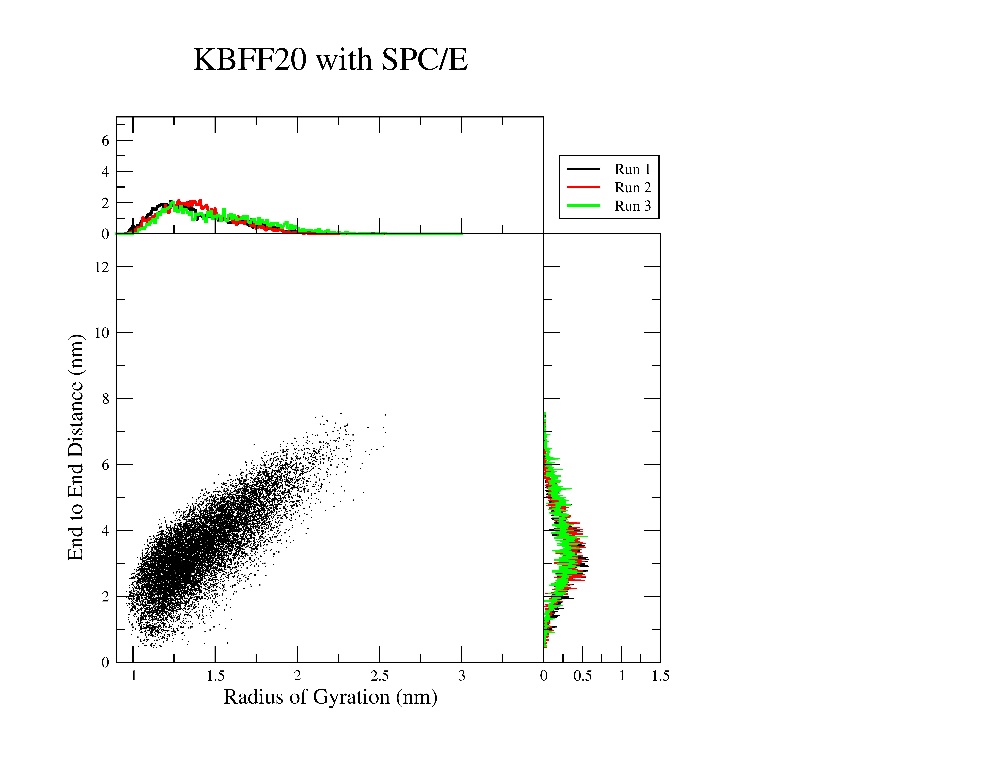

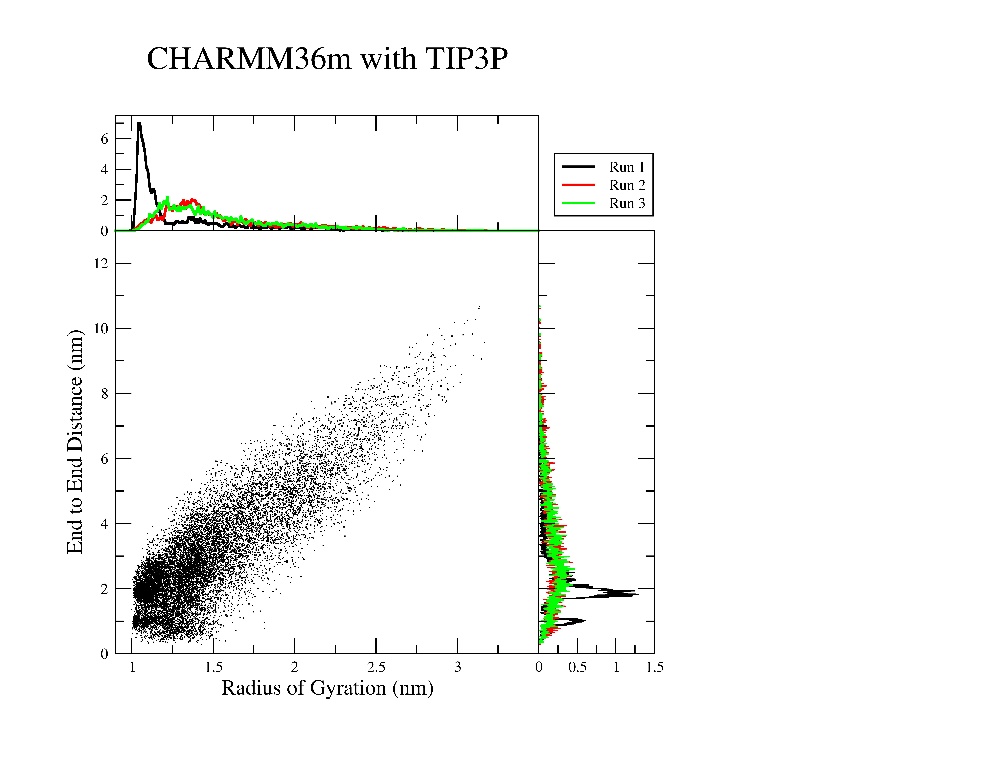

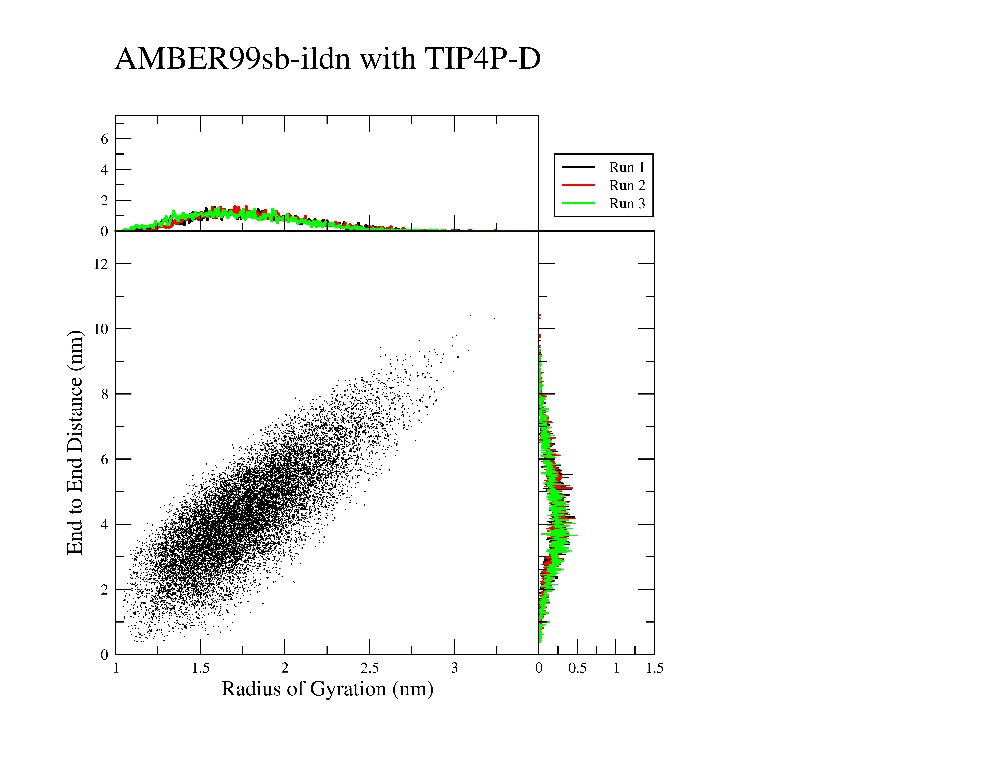

**Supplementary figure 4.** **AlphaFold2 prediction of the CIITA structure.** The structure for CIITA predicted by AlphaFold2^7^ for Uniprot P33076 was represented using UCSF Chimera v1.15^8^. The N-terminal TAD is in gold and yellow (residues 56-92). Positions 56-91 are shown in the inset with four positions with high disorder probabilities in red (I62, Y65, T70, T91) and three with low disorder probabilities in blue (N74, Q77, F78). Note that position 91 was the most neutral of the seven in our substitution studies.

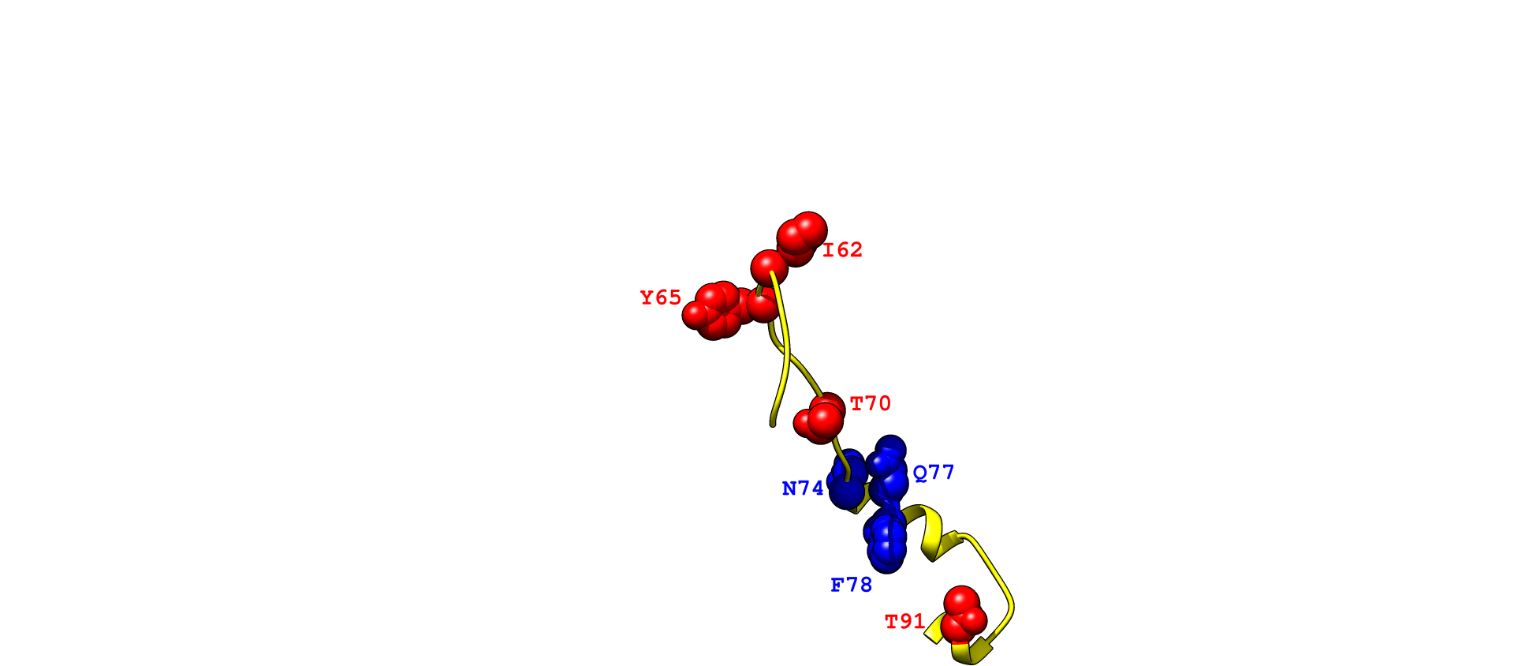

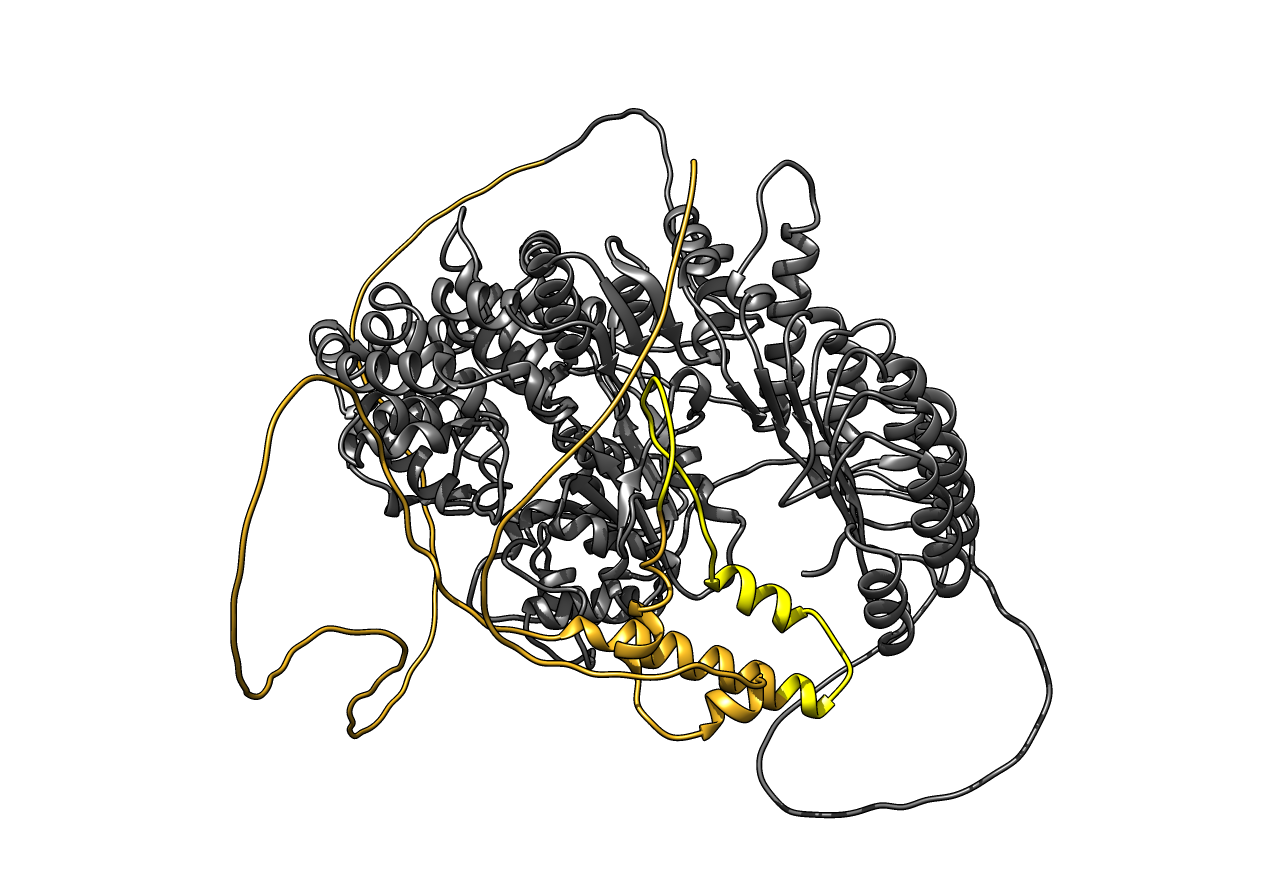

**Supplementary table 2. List of predicted short linear motifs (SLiMs) from Eukaryotic Linear Motifs (ELM) for the activation domain of CIITA (Uniprot P33076).** The first column shows the CIITA positions that comprise predicted SLiMs. The fourteen SLiMs that overlap the region tested in this study (positions 61-92) are highlighted in grey.

| **Predicted CIITA SLiMs** | **ELM description** |
| --- | --- |
| 16-20 | The USP7 MATH domain binding motif variant based on the MDM2 and p53 interactions. |
| 17-24 | GSK3 phosphorylation recognition site |
| 63-70 |  |
| 17-23 | (ST)Q motif which is phosphorylated by PIKK family members |
| 139-145 |  |
| 22-28 | Phosphothreonine motif binding a subset of FHA domains that show a preference for a large aliphatic amino acid at the pT+3 position. |
| 68-74 |  |
| 27-35 | TRAF6 binding site. Members of the tumour necrosis factor receptor (TNFR) superfamily initiate intracellular signalling by recruiting the C-domains of the TNFR-associated factors (TRAFs) using motifs in their cytoplasmic tails. |
| 31-42 | Canonical LIR motif that binds to Atg8/LC3 protein family members to mediate processes involved in autophagy. |
| 76-85 |  |
| 93-101 |  |
| 31-37 | Nematode-specific variant of the canonical LIR motif that binds to Atg8 protein family members to mediate processes involved in autophagy. |
| 76-81 |  |
| 93-98 |  |
| 117-122 |  |
| 34-37 | Fungi-specific variant of the WDR5-binding motif that binds to a cleft between blades 5 and 6 of the WD40 repeat domain of WDR5, opposite of the Win motif-binding site, to mediate assembly of histone modification complexes. |
| 59-64 |  |
| 60-64 |  |
| 95-101 |  |
| 131-138 |  |
| 133-138 |  |
| 149-155 |  |
| 150-155 |  |
| 41-45 | These IBMs are found at the N-terminal regions of caspase subunits where they mediate the inhibition of activated caspases by binding to conserved surface grooves on type III BIR domains of Inhibitor of Apoptosis Proteins (IAPs). |
| 45-57 | Reverse NES binding the CRM1 groove in the minus direction. The spacing of the initial two hydrophobic residues ΦxΦ dictates the reverse orientation |
| 51-54 | Tyrosine-based sorting signal responsible for the interaction with mu subunit of AP (Adaptor Protein) complex |
| 57-60 | Major TRAF2-binding consensus motif. Members of the tumor necrosis factor receptor (TNFR) superfamily initiate intracellular signaling by recruiting the C-domain of the TNFR-associated factors (TRAFs) through their cytoplasmic tails. |
| 59-65 | Acidic dileucine motifs with a monoleucine preference and extra glutamate sorting in Endosomal-Basolateral trafficking. |
| 61-64 |  |
| 62-67 | Amphipathic motif that is involved in APC/C inhibition by binding of CDH1/CDC20. In metazoan cyclin A, the motif also acts as a degron, enabling the cyclin's degradation in prometaphase. |
| 63-68 | Apicomplexa specific variant of the canonical LIR motif that binds to Atg8 protein family members to mediate processes involved in autophagy. |
| 63-69 | Ser/Thr residue phosphorylated by Plk2 and Plk3 |
| 65-69 | NCK Src Homology 2 (SH2) domain binding motif |
| 95-99 | CRK family SH2 domain binding motif. |
| 77-82 | Dileucine motifs lacking Glu+1 with Pro-Arg preference at +4 sorting in Endosomal-Basolateral-Lysosomal trafficking. |
| 80-84 | Plasmodium Export Element, PEXEL, is a trafficking signal for protein cleavage by PMV protease and export from Plasmodium parasites to infected host cells. |
| 95-100 | Docking site required for the regulatory subunit B56 of PP2A for protein dephosphorylation. |
| 95-98 | GRB2-like Src Homology 2 (SH2) domains binding motif |
| 104-107 | YXXQ motif found in the cytoplasmic region of cytokine receptors that bind STAT3 SH2 domain. |
| 106-112 | Casein kinase 2 (CK2) phosphorylation site |
| 116-120 | Subtilisin/kexin isozyme-1 (SKI1) cleavage site ([RK]-X-[hydrophobic]-[LTKF]-\|-X). |
| 144-146 | Yeast kexin 2 cleavage site (K-R-\|-X or R-R-\|-X). |
| 157-163 | This is the motif recognized by those SH3 domains with a non-canonical class I recognition specificity. |

**Supplementary figure 5. Experimental design for testing outcomes of singly substituted variants of CIITA.** The oval represents HEK293 cells that were co-transfected with three plasmids: (i) The “CIITA” plasmid expressed the WT or the CIITA variant under the control of constitutively active promoter hEF1-HTLV, (ii) the “Reporter” plasmid expressed the *Gaussia* luciferase (G-Luc) gene under the control of the HLA-DRA promoter that is responsive to CIITA), (iii) the “Transfection control” plasmid expressed *Cyrpridina* luciferase (C-Luc) under the control of the constitutively active promoter SV40. Four independent measurements were made for the WT and each CIITA variant: (1) The signal from the G-Luc reporter under control by CIITA variants. (2) The signal from the C-Luc control for transfection efficiency. (3) Western blots that measure the cellular concentration of CIITA protein. (4) Western blots of endogenous β-tubulin, to control for cell density in the sample. These values were used in equations below (equations 1-3 in the main text) to compute *in vivo* function, cellular concentration, and transcriptional activation for each CIITA variant.

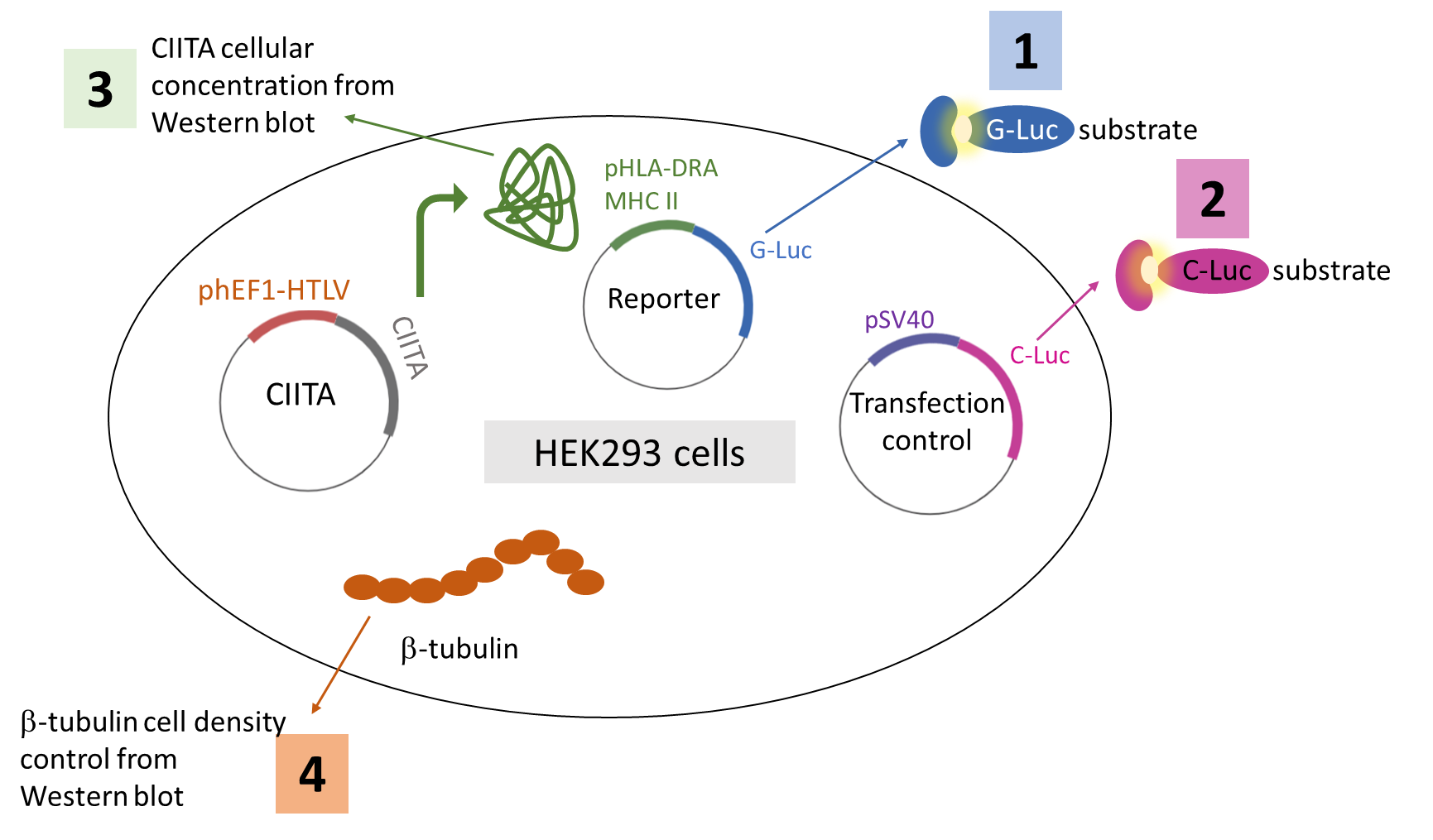

Eqn 1. *In vivo* function = (CIITA reporter activity/Transfection efficiency)/(Cell density)

Eqn 2. Cellular concentration = (CIITA protein levels)/(Transfection efficiency/Cell density)

Eqn 3. Transcriptional activation = (CIITA reporter activity/Transcription efficiency)/(CIITA protein levels/Cell density)

**Supplementary figure 6 (next page). Effects of single substitutions on *in vivo* function of variants of CIITA**. All data were normalized to a WT value that was set to 1. **(a)** Reference range for *in vivo* function: At the lower end of the range, the hypothetical “Dead” CIITA was determined from a control sample that lacked CIITA (dotted line at 0.15; calculations in Supplementary table 7). The upper end of the range was defined by the variant with the greatest transcriptional activity, Y65H (“Max Observed”, dotted line at 4). All values for the biological and technical replicates of WT are shown with dots; the averages of these values were normalized to 1 and the standard errors of the means are shown with error bars. **(b)-(h)** Normalized values of *in vivo* function are shown for each substitution at positions with low **(b)** N74, **(c)** Q77, **(d)** F78 and high disorder probabilities **(e)** I62, **(f)** Y65, **(g)** T70, **(h)** T91. Individual measurements are shown with black dots, and standard errors of the means are indicated with error bars. The variant corresponding to WT is circled on the X-axis legend, and the normalized WT value of 1 is indicated with a dashed line; the WT samples shown on **(b)-(h)** were measured in parallel with each position’s variants. Bars are coloured according to amino acid type: acidic (D, E; red, checked), basic (R, H, K; blue, slanted lines), polar uncharged (S, T, N, C, Q; yellow, vertical lines), hydrophobic (M, F, W, Y, A, V, I, L; green, horizontal lines), and structure breakers (P, G; grey, solid). Variants with activity statistically different from WT were identified with (i) one-way ANOVAs with Dunnett’s correction (****p <0.0001, ***p<0.001, **p<0.01, *p<0.1), and (ii) false discovery rate analyses (underlined variants in the X-axis legend).

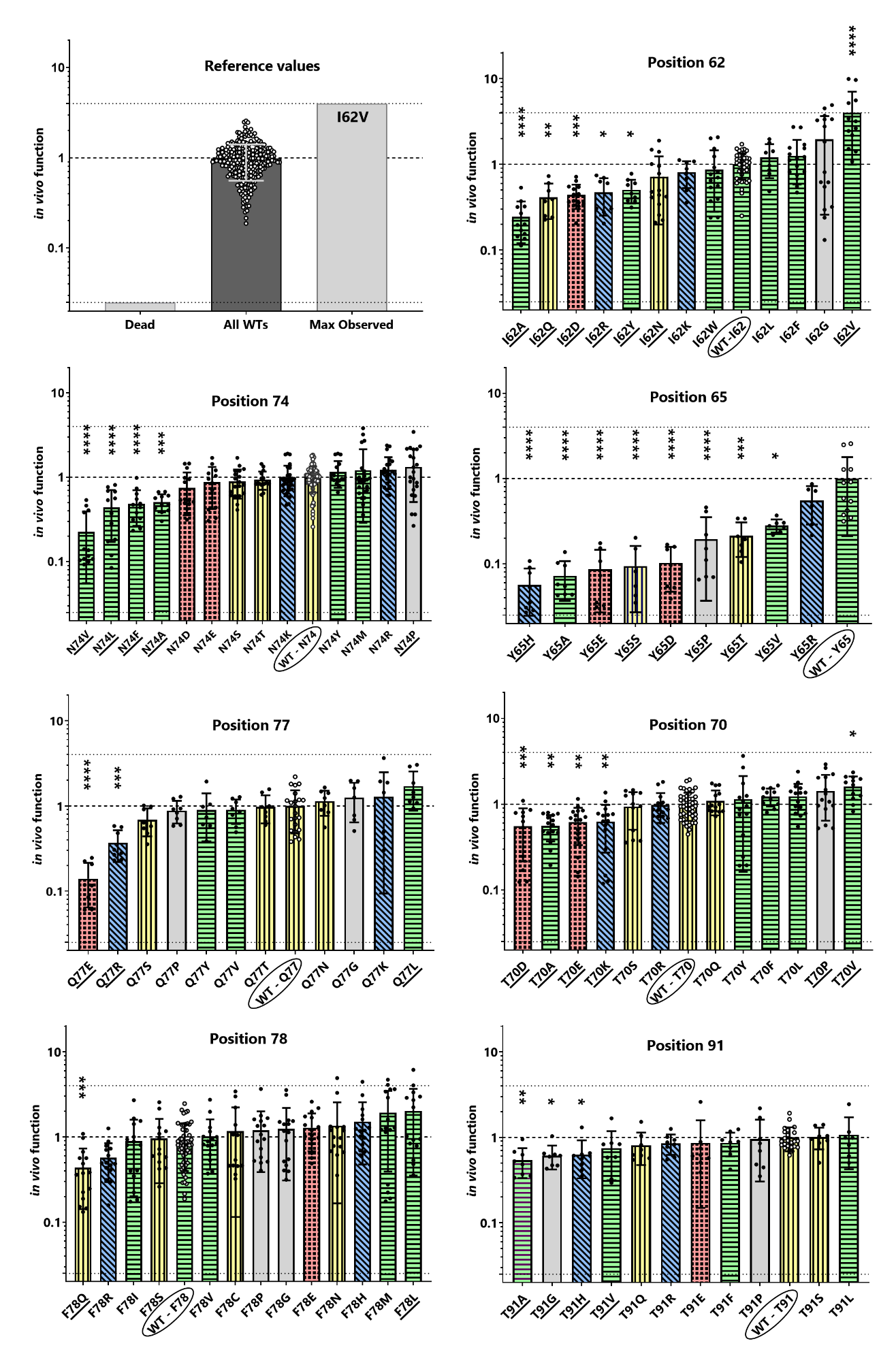

**(h)**

**(g)**

**(f)**

**(e)**

**(d)**

**(c)**

**(b)**

**(a)**

**Supplementary table 3. Numbers of CIITA variants with statistically-significant enhanced or diminished outcomes compared to WT.**

|  | No. of variants (%) with diminished outcomes | No. of variants (%) with enhanced outcomes | No. of variants (%) with WT-like outcomes |
| --- | --- | --- | --- |
| ***In vivo* function:** |  |  |  |
| One way ANOVA | 28 (34 %) | 2 (2.5 %) | 52 (63.5 %) |
| FDR | 30 (36.5 %) | 6 (7.3 %) | 46 (56.2 %) |
| **Cellular concentration:** |  |  |  |
| One way ANOVA | 15 (18.3 %) | 9 (11 %) | 58 (70.7 %) |
| FDR | 22 (26.8 %) | 12 (14.6 %) | 48 (58.6 %) |
| **Transcriptional activation:** |  |  |  |
| One way ANOVA | 13 (15.8 %) | 16 (19.5 %) | 53 (64.7 %) |
| FDR | 20 (24.4 %) | 23 (28 %) | 39 (47.6 %) |

**Supplementary figure 7. Comparison of intrinsic disorder with conservation.** Correlation plots for PONDR disorder probabilities versus ConSurf raw scores obtained for **(a)-(b)** the N-terminal activation domain and **(c)-(d)** full length CIITA. **(a)** and **(c)** show the average scores from 19 subsets of “CIITA_manual_MSA”, and **(b)** and **(d)** show “CIITA_default_MSA”.

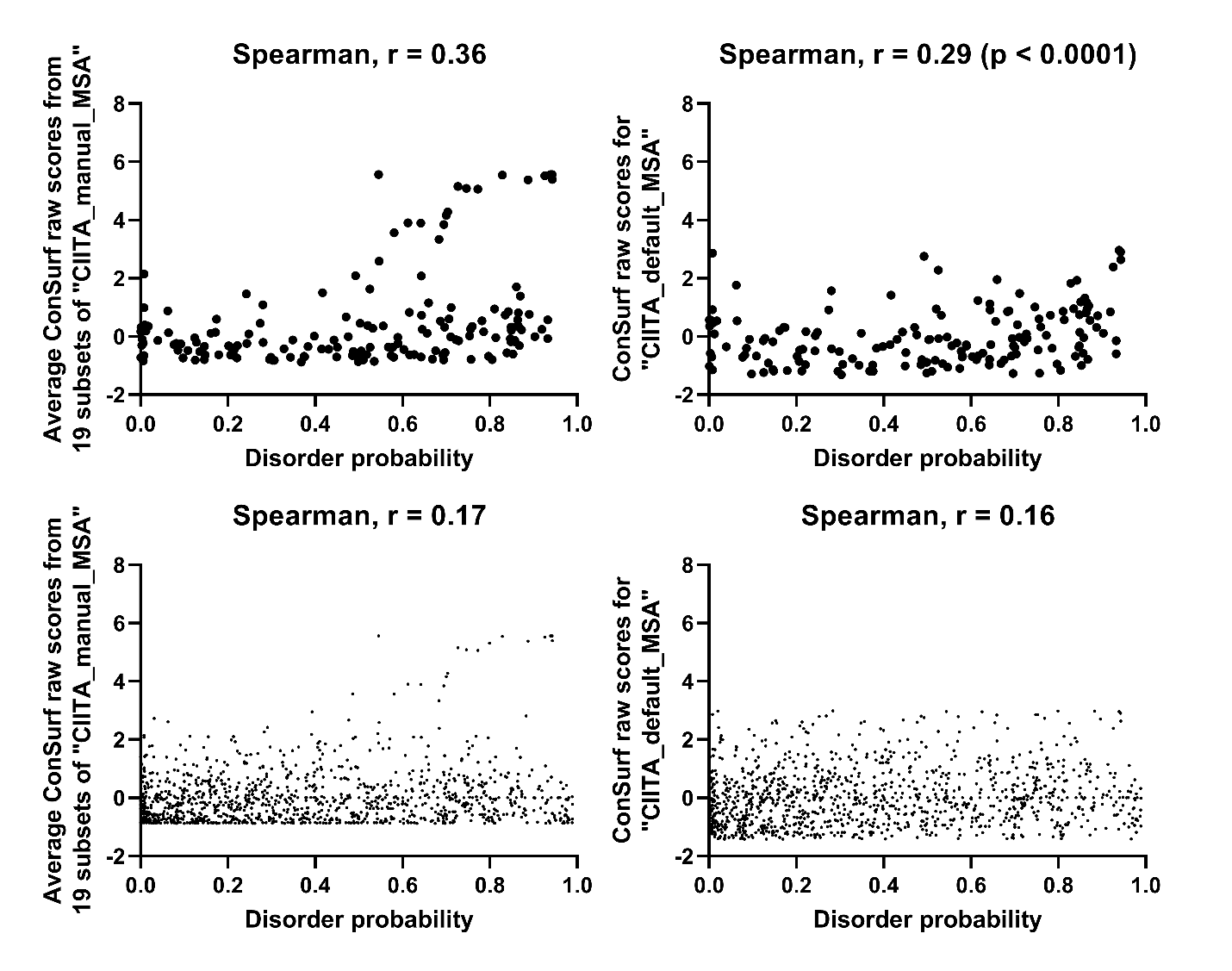

**(d)**

**(c)**

**(b)**

**(a)**

**Supplementary table 4.** **ConSurf scores for the full-length CIITA sequences from two alternative approaches.** In ConSurf analyses, the range of raw scores (Supplementary Figure 2) are partitioned into 9 bins. ConSurf bin scores of 1 indicate the least conserved positions and bin scores of 9 indicate the most conserved positions. In the table below, column **(a)** contains the ConSurf grades for CIITA_default_MSA (aligned by MAFFT-L-Ins-I); column **(b)** contains the average scores of 19 MSAs subsets with 300-450 sequences extracted from CIITA_manual_MSA (aligned by clustalOmega). To aid visual inspection, Consurf scores between 1 and 9 were colour-coded using a three-colour gradient (red-yellow-green). The seven positions studied in the experiments are highlighted with red (62, 65, 70, 91; high disorder probability) and blue (74, 77, 78; low disorder probability) fonts with grey backgrounds.

| 1 | 2 | 3 | 4 | 5 | 6 | 7 | 8 | 9 |
| --- | --- | --- | --- | --- | --- | --- | --- | --- |

| **Pos#** | **a** | **b** | **Pos#** | **a** | **b** | **Pos#** | **a** | **b** | **Pos#** | **a** | **b** | **Pos#** | **a** | **b** | **Pos#** | **a** | **b** |
| --- | --- | --- | --- | --- | --- | --- | --- | --- | --- | --- | --- | --- | --- | --- | --- | --- | --- |
| 1 | 8 | 3 | 201 | 1 | 1 | 401 | 8 | 9 | 601 | 2 | 5 | 801 | 8 | 9 | 1001 | 9 | 9 |
| 2 | 7 | 2 | 202 | 3 | 4 | 402 | 3 | 4 | 602 | 5 | 8 | 802 | 4 | 4 | 1002 | 5 | 5 |
| 3 | 6 | 2 | 203 | 4 | 5 | 403 | 4 | 6 | 603 | 6 | 5 | 803 | 7 | 6 | 1003 | 8 | 9 |
| 4 | 5 | 2 | 204 | 3 | 5 | 404 | 3 | 4 | 604 | 8 | 9 | 804 | 4 | 5 | 1004 | 7 | 9 |
| 5 | 8 | 2 | 205 | 4 | 5 | 405 | 1 | 3 | 605 | 5 | 5 | 805 | 6 | 7 | 1005 | 4 | 5 |
| 6 | 8 | 2 | 206 | 1 | 3 | 406 | 3 | 4 | 606 | 8 | 6 | 806 | 8 | 9 | 1006 | 5 | 6 |
| 7 | 4 | 2 | 207 | 3 | 4 | 407 | 2 | 4 | 607 | 8 | 8 | 807 | 9 | 9 | 1007 | 9 | 9 |
| 8 | 7 | 2 | 208 | 1 | 3 | 408 | 5 | 5 | 608 | 6 | 5 | 808 | 8 | 9 | 1008 | 7 | 6 |
| 9 | 5 | 1 | 209 | 3 | 4 | 409 | 4 | 4 | 609 | 7 | 9 | 809 | 8 | 9 | 1009 | 6 | 8 |
| 10 | 3 | 1 | 210 | 3 | 4 | 410 | 1 | 4 | 610 | 7 | 8 | 810 | 6 | 9 | 1010 | 7 | 7 |
| 11 | 5 | 1 | 211 | 3 | 4 | 411 | 5 | 5 | 611 | 9 | 9 | 811 | 8 | 9 | 1011 | 7 | 6 |
| 12 | 2 | 1 | 212 | 2 | 4 | 412 | 6 | 8 | 612 | 4 | 5 | 812 | 8 | 9 | 1012 | 6 | 8 |
| 13 | 5 | 1 | 213 | 5 | 6 | 413 | 8 | 9 | 613 | 5 | 5 | 813 | 6 | 7 | 1013 | 4 | 5 |
| 14 | 1 | 1 | 214 | 2 | 5 | 414 | 5 | 4 | 614 | 9 | 9 | 814 | 7 | 9 | 1014 | 9 | 9 |
| 15 | 1 | 1 | 215 | 5 | 5 | 415 | 7 | 9 | 615 | 5 | 4 | 815 | 9 | 9 | 1015 | 5 | 8 |
| 16 | 1 | 1 | 216 | 4 | 5 | 416 | 6 | 7 | 616 | 5 | 6 | 816 | 8 | 8 | 1016 | 7 | 7 |
| 17 | 1 | 1 | 217 | 1 | 4 | 417 | 7 | 8 | 617 | 5 | 5 | 817 | 3 | 4 | 1017 | 9 | 9 |
| 18 | 7 | 5 | 218 | 5 | 5 | 418 | 7 | 8 | 618 | 8 | 9 | 818 | 4 | 6 | 1018 | 9 | 9 |
| 19 | 5 | 6 | 219 | 5 | 6 | 419 | 8 | 9 | 619 | 5 | 6 | 819 | 3 | 5 | 1019 | 7 | 9 |
| 20 | 4 | 5 | 220 | 5 | 5 | 420 | 9 | 9 | 620 | 6 | 9 | 820 | 3 | 4 | 1020 | 9 | 9 |
| 21 | 5 | 5 | 221 | 4 | 5 | 421 | 7 | 6 | 621 | 6 | 5 | 821 | 7 | 6 | 1021 | 9 | 9 |
| 22 | 4 | 4 | 222 | 5 | 6 | 422 | 9 | 8 | 622 | 6 | 6 | 822 | 3 | 6 | 1022 | 9 | 9 |
| 23 | 4 | 5 | 223 | 5 | 5 | 423 | 9 | 9 | 623 | 5 | 5 | 823 | 4 | 5 | 1023 | 9 | 9 |
| 24 | 4 | 4 | 224 | 4 | 5 | 424 | 5 | 5 | 624 | 3 | 6 | 824 | 7 | 8 | 1024 | 9 | 9 |
| 25 | 9 | 9 | 225 | 6 | 6 | 425 | 9 | 9 | 625 | 4 | 6 | 825 | 8 | 9 | 1025 | 9 | 9 |
| 26 | 6 | 5 | 226 | 2 | 4 | 426 | 9 | 9 | 626 | 3 | 4 | 826 | 5 | 5 | 1026 | 5 | 6 |
| 27 | 5 | 5 | 227 | 4 | 5 | 427 | 9 | 9 | 627 | 1 | 4 | 827 | 4 | 4 | 1027 | 9 | 9 |
| 28 | 5 | 6 | 228 | 8 | 8 | 428 | 5 | 4 | 628 | 4 | 4 | 828 | 5 | 6 | 1028 | 9 | 9 |
| 29 | 3 | 4 | 229 | 3 | 4 | 429 | 5 | 5 | 629 | 1 | 3 | 829 | 8 | 9 | 1029 | 9 | 9 |
| 30 | 4 | 5 | 230 | 5 | 5 | 430 | 7 | 7 | 630 | 1 | 3 | 830 | 6 | 9 | 1030 | 4 | 6 |
| 31 | 9 | 5 | 231 | 5 | 5 | 431 | 5 | 4 | 631 | 7 | 6 | 831 | 3 | 4 | 1031 | 9 | 9 |
| 32 | 8 | 8 | 232 | 5 | 5 | 432 | 6 | 5 | 632 | 2 | 4 | 832 | 1 | 4 | 1032 | 8 | 9 |
| 33 | 4 | 4 | 233 | 3 | 4 | 433 | 8 | 8 | 633 | 8 | 9 | 833 | 9 | 9 | 1033 | 5 | 6 |
| 34 | 7 | 9 | 234 | 3 | 5 | 434 | 7 | 6 | 634 | 9 | 9 | 834 | 8 | 9 | 1034 | 4 | 5 |
| 35 | 9 | 9 | 235 | 5 | 5 | 435 | 6 | 5 | 635 | 7 | 7 | 835 | 9 | 9 | 1035 | 9 | 9 |
| 36 | 5 | 6 | 236 | 5 | 7 | 436 | 8 | 6 | 636 | 9 | 9 | 836 | 4 | 7 | 1036 | 9 | 9 |
| 37 | 9 | 9 | 237 | 5 | 5 | 437 | 9 | 9 | 637 | 9 | 9 | 837 | 6 | 9 | 1037 | 3 | 5 |
| 38 | 9 | 9 | 238 | 7 | 9 | 438 | 8 | 9 | 638 | 9 | 9 | 838 | 5 | 9 | 1038 | 8 | 9 |
| 39 | 7 | 7 | 239 | 5 | 6 | 439 | 2 | 3 | 639 | 4 | 7 | 839 | 5 | 9 | 1039 | 9 | 9 |
| 40 | 9 | 9 | 240 | 4 | 5 | 440 | 8 | 9 | 640 | 6 | 9 | 840 | 9 | 9 | 1040 | 7 | 9 |
| 41 | 2 | 4 | 241 | 4 | 5 | 441 | 4 | 6 | 641 | 4 | 6 | 841 | 5 | 5 | 1041 | 5 | 6 |
| 42 | 4 | 5 | 242 | 4 | 5 | 442 | 6 | 9 | 642 | 6 | 9 | 842 | 9 | 8 | 1042 | 8 | 9 |
| 43 | 9 | 9 | 243 | 6 | 8 | 443 | 6 | 8 | 643 | 4 | 4 | 843 | 6 | 6 | 1043 | 8 | 9 |
| 44 | 8 | 9 | 244 | 5 | 7 | 444 | 8 | 9 | 644 | 5 | 9 | 844 | 9 | 9 | 1044 | 4 | 5 |
| 45 | 4 | 5 | 245 | 5 | 6 | 445 | 8 | 7 | 645 | 5 | 7 | 845 | 6 | 5 | 1045 | 8 | 8 |
| 46 | 4 | 5 | 246 | 2 | 4 | 446 | 7 | 9 | 646 | 5 | 5 | 846 | 6 | 5 | 1046 | 9 | 9 |
| 47 | 7 | 7 | 247 | 5 | 5 | 447 | 8 | 9 | 647 | 5 | 5 | 847 | 7 | 5 | 1047 | 5 | 5 |
| 48 | 4 | 6 | 248 | 4 | 5 | 448 | 8 | 7 | 648 | 5 | 7 | 848 | 8 | 9 | 1048 | 7 | 8 |
| 49 | 7 | 6 | 249 | 4 | 5 | 449 | 9 | 9 | 649 | 4 | 5 | 849 | 3 | 4 | 1049 | 8 | 9 |
| 50 | 1 | 3 | 250 | 5 | 5 | 450 | 2 | 4 | 650 | 1 | 4 | 850 | 4 | 4 | 1050 | 9 | 9 |
| 51 | 4 | 4 | 251 | 8 | 8 | 451 | 4 | 4 | 651 | 2 | 5 | 851 | 4 | 9 | 1051 | 9 | 9 |
| 52 | 9 | 8 | 252 | 5 | 6 | 452 | 6 | 7 | 652 | 3 | 5 | 852 | 8 | 9 | 1052 | 9 | 9 |
| 53 | 4 | 5 | 253 | 5 | 6 | 453 | 5 | 7 | 653 | 5 | 9 | 853 | 4 | 4 | 1053 | 8 | 9 |
| 54 | 5 | 5 | 254 | 3 | 4 | 454 | 5 | 5 | 654 | 4 | 6 | 854 | 5 | 6 | 1054 | 9 | 9 |
| 55 | 6 | 6 | 255 | 6 | 8 | 455 | 3 | 4 | 655 | 4 | 6 | 855 | 8 | 9 | 1055 | 2 | 5 |
| 56 | 7 | 6 | 256 | 4 | 6 | 456 | 5 | 7 | 656 | 5 | 8 | 856 | 3 | 5 | 1056 | 9 | 9 |
| 57 | 5 | 5 | 257 | 4 | 6 | 457 | 6 | 7 | 657 | 7 | 9 | 857 | 4 | 4 | 1057 | 5 | 5 |
| 58 | 3 | 4 | 258 | 4 | 5 | 458 | 5 | 4 | 658 | 2 | 4 | 858 | 5 | 7 | 1058 | 9 | 9 |
| 59 | 6 | 7 | 259 | 4 | 5 | 459 | 4 | 5 | 659 | 2 | 4 | 859 | 8 | 9 | 1059 | 2 | 3 |
| 60 | 8 | 8 | 260 | 3 | 4 | 460 | 4 | 9 | 660 | 9 | 8 | 860 | 7 | 7 | 1060 | 9 | 9 |
| 61 | 8 | 8 | 261 | 4 | 5 | 461 | 4 | 4 | 661 | 8 | 9 | 861 | 8 | 9 | 1061 | 9 | 9 |
| **62** | 2 | 4 | 262 | 5 | 5 | 462 | 2 | 4 | 662 | 2 | 5 | 862 | 9 | 9 | 1062 | 5 | 5 |
| 63 | 6 | 5 | 263 | 5 | 7 | 463 | 5 | 5 | 663 | 4 | 9 | 863 | 9 | 9 | 1063 | 5 | 8 |
| 64 | 4 | 6 | 264 | 6 | 5 | 464 | 6 | 5 | 664 | 9 | 9 | 864 | 7 | 9 | 1064 | 8 | 9 |
| **65** | 3 | 4 | 265 | 5 | 6 | 465 | 8 | 9 | 665 | 6 | 9 | 865 | 5 | 5 | 1065 | 9 | 9 |
| 66 | 8 | 8 | 266 | 1 | 4 | 466 | 4 | 5 | 666 | 4 | 8 | 866 | 8 | 9 | 1066 | 4 | 5 |
| 67 | 8 | 8 | 267 | 3 | 4 | 467 | 7 | 9 | 667 | 4 | 5 | 867 | 5 | 8 | 1067 | 7 | 6 |
| 68 | 5 | 5 | 268 | 5 | 5 | 468 | 9 | 9 | 668 | 7 | 9 | 868 | 7 | 7 | 1068 | 9 | 9 |
| 69 | 7 | 8 | 269 | 4 | 5 | 469 | 9 | 9 | 669 | 8 | 8 | 869 | 7 | 6 | 1069 | 9 | 9 |
| **70** | 4 | 5 | 270 | 5 | 5 | 470 | 5 | 5 | 670 | 3 | 4 | 870 | 4 | 7 | 1070 | 3 | 5 |
| 71 | 8 | 9 | 271 | 4 | 4 | 471 | 5 | 6 | 671 | 6 | 9 | 871 | 5 | 5 | 1071 | 8 | 9 |
| 72 | 8 | 8 | 272 | 3 | 5 | 472 | 4 | 5 | 672 | 5 | 4 | 872 | 9 | 7 | 1072 | 3 | 5 |
| 73 | 8 | 9 | 273 | 4 | 5 | 473 | 6 | 5 | 673 | 4 | 5 | 873 | 8 | 9 | 1073 | 8 | 8 |
| **74** | 8 | 9 | 274 | 5 | 5 | 474 | 4 | 5 | 674 | 4 | 5 | 874 | 5 | 5 | 1074 | 9 | 9 |
| 75 | 6 | 7 | 275 | 4 | 5 | 475 | 4 | 5 | 675 | 5 | 6 | 875 | 4 | 5 | 1075 | 7 | 9 |
| 76 | 5 | 6 | 276 | 6 | 7 | 476 | 5 | 5 | 676 | 3 | 4 | 876 | 8 | 9 | 1076 | 8 | 8 |
| **77** | 9 | 9 | 277 | 5 | 7 | 477 | 4 | 4 | 677 | 4 | 5 | 877 | 8 | 9 | 1077 | 7 | 6 |
| **78** | 5 | 5 | 278 | 8 | 8 | 478 | 4 | 5 | 678 | 1 | 3 | 878 | 5 | 8 | 1078 | 7 | 9 |
| 79 | 6 | 7 | 279 | 5 | 5 | 479 | 3 | 4 | 679 | 4 | 5 | 879 | 9 | 9 | 1079 | 8 | 9 |
| 80 | 4 | 5 | 280 | 8 | 8 | 480 | 5 | 7 | 680 | 2 | 4 | 880 | 4 | 5 | 1080 | 8 | 9 |
| 81 | 7 | 9 | 281 | 7 | 9 | 481 | 3 | 5 | 681 | 4 | 5 | 881 | 8 | 6 | 1081 | 6 | 6 |
| 82 | 5 | 6 | 282 | 4 | 5 | 482 | 6 | 7 | 682 | 8 | 9 | 882 | 8 | 9 | 1082 | 9 | 9 |
| 83 | 3 | 4 | 283 | 7 | 9 | 483 | 8 | 8 | 683 | 3 | 5 | 883 | 4 | 7 | 1083 | 8 | 9 |
| 84 | 5 | 5 | 284 | 5 | 7 | 484 | 5 | 4 | 684 | 4 | 5 | 884 | 4 | 3 | 1084 | 8 | 8 |
| 85 | 4 | 5 | 285 | 5 | 7 | 485 | 3 | 4 | 685 | 4 | 5 | 885 | 8 | 9 | 1085 | 9 | 9 |
| 86 | 6 | 7 | 286 | 8 | 9 | 486 | 6 | 6 | 686 | 4 | 5 | 886 | 9 | 9 | 1086 | 7 | 8 |
| 87 | 3 | 4 | 287 | 5 | 7 | 487 | 6 | 6 | 687 | 6 | 6 | 887 | 8 | 9 | 1087 | 4 | 5 |
| 88 | 7 | 9 | 288 | 8 | 9 | 488 | 5 | 4 | 688 | 5 | 8 | 888 | 6 | 5 | 1088 | 9 | 9 |
| 89 | 8 | 9 | 289 | 7 | 9 | 489 | 6 | 6 | 689 | 8 | 9 | 889 | 7 | 9 | 1089 | 9 | 9 |
| 90 | 6 | 8 | 290 | 6 | 8 | 490 | 6 | 5 | 690 | 6 | 6 | 890 | 6 | 9 | 1090 | 8 | 9 |
| **91** | 7 | 8 | 291 | 6 | 8 | 491 | 8 | 8 | 691 | 5 | 6 | 891 | 9 | 9 | 1091 | 6 | 8 |
| 92 | 4 | 5 | 292 | 9 | 9 | 492 | 3 | 5 | 692 | 1 | 3 | 892 | 7 | 8 | 1092 | 9 | 9 |
| 93 | 7 | 8 | 293 | 7 | 9 | 493 | 7 | 8 | 693 | 5 | 8 | 893 | 6 | 7 | 1093 | 7 | 6 |
| 94 | 7 | 7 | 294 | 7 | 7 | 494 | 7 | 5 | 694 | 5 | 6 | 894 | 3 | 4 | 1094 | 5 | 8 |
| 95 | 8 | 9 | 295 | 5 | 7 | 495 | 9 | 9 | 695 | 4 | 6 | 895 | 9 | 9 | 1095 | 9 | 8 |
| 96 | 8 | 8 | 296 | 8 | 9 | 496 | 8 | 7 | 696 | 3 | 4 | 896 | 8 | 9 | 1096 | 9 | 9 |
| 97 | 5 | 7 | 297 | 5 | 8 | 497 | 5 | 8 | 697 | 5 | 5 | 897 | 8 | 8 | 1097 | 6 | 6 |
| 98 | 9 | 9 | 298 | 8 | 9 | 498 | 8 | 9 | 698 | 3 | 3 | 898 | 7 | 9 | 1098 | 5 | 6 |
| 99 | 9 | 9 | 299 | 5 | 5 | 499 | 9 | 9 | 699 | 4 | 5 | 899 | 9 | 9 | 1099 | 7 | 8 |
| 100 | 5 | 6 | 300 | 5 | 9 | 500 | 5 | 5 | 700 | 4 | 4 | 900 | 5 | 5 | 1100 | 7 | 9 |
| 101 | 9 | 8 | 301 | 8 | 9 | 501 | 6 | 7 | 701 | 4 | 5 | 901 | 4 | 6 | 1101 | 4 | 5 |
| 102 | 8 | 8 | 302 | 7 | 9 | 502 | 7 | 9 | 702 | 3 | 4 | 902 | 4 | 4 | 1102 | 5 | 6 |
| 103 | 7 | 7 | 303 | 4 | 6 | 503 | 6 | 7 | 703 | 3 | 4 | 903 | 4 | 7 | 1103 | 6 | 5 |
| 104 | 9 | 8 | 304 | 7 | 9 | 504 | 6 | 8 | 704 | 1 | 4 | 904 | 7 | 7 | 1104 | 8 | 8 |
| 105 | 9 | 8 | 305 | 6 | 8 | 505 | 6 | 9 | 705 | 4 | 5 | 905 | 4 | 5 | 1105 | 7 | 8 |
| 106 | 8 | 8 | 306 | 6 | 8 | 506 | 7 | 5 | 706 | 4 | 7 | 906 | 3 | 4 | 1106 | 6 | 7 |
| 107 | 7 | 7 | 307 | 6 | 8 | 507 | 5 | 6 | 707 | 5 | 6 | 907 | 7 | 9 | 1107 | 8 | 8 |
| 108 | 9 | 9 | 308 | 5 | 7 | 508 | 6 | 6 | 708 | 8 | 9 | 908 | 9 | 9 | 1108 | 8 | 7 |
| 109 | 5 | 6 | 309 | 3 | 5 | 509 | 4 | 5 | 709 | 6 | 5 | 909 | 4 | 5 | 1109 | 9 | 8 |
| 110 | 6 | 7 | 310 | 1 | 4 | 510 | 3 | 4 | 710 | 7 | 8 | 910 | 4 | 6 | 1110 | 7 | 8 |
| 111 | 4 | 5 | 311 | 1 | 3 | 511 | 5 | 6 | 711 | 7 | 9 | 911 | 7 | 6 | 1111 | 8 | 9 |
| 112 | 7 | 6 | 312 | 3 | 5 | 512 | 6 | 7 | 712 | 4 | 6 | 912 | 5 | 8 | 1112 | 9 | 8 |
| 113 | 5 | 5 | 313 | 6 | 8 | 513 | 3 | 5 | 713 | 6 | 8 | 913 | 4 | 5 | 1113 | 9 | 9 |
| 114 | 5 | 5 | 314 | 2 | 4 | 514 | 2 | 4 | 714 | 9 | 9 | 914 | 9 | 9 | 1114 | 7 | 7 |
| 115 | 5 | 7 | 315 | 2 | 4 | 515 | 2 | 3 | 715 | 8 | 9 | 915 | 8 | 9 | 1115 | 8 | 7 |
| 116 | 6 | 6 | 316 | 1 | 4 | 516 | 4 | 4 | 716 | 8 | 9 | 916 | 6 | 9 | 1116 | 8 | 9 |
| 117 | 8 | 9 | 317 | 5 | 6 | 517 | 4 | 5 | 717 | 8 | 9 | 917 | 8 | 8 | 1117 | 8 | 7 |
| 118 | 6 | 7 | 318 | 3 | 4 | 518 | 3 | 3 | 718 | 5 | 6 | 918 | 5 | 8 | 1118 | 7 | 8 |
| 119 | 8 | 9 | 319 | 4 | 5 | 519 | 2 | 4 | 719 | 8 | 9 | 919 | 9 | 9 | 1119 | 8 | 8 |
| 120 | 5 | 1 | 320 | 4 | 5 | 520 | 1 | 4 | 720 | 3 | 4 | 920 | 8 | 9 | 1120 | 9 | 9 |
| 121 | 6 | 6 | 321 | 1 | 2 | 521 | 5 | 8 | 721 | 8 | 7 | 921 | 8 | 8 | 1121 | 6 | 8 |
| 122 | 7 | 7 | 322 | 3 | 4 | 522 | 3 | 5 | 722 | 8 | 9 | 922 | 8 | 8 | 1122 | 8 | 7 |
| 123 | 6 | 5 | 323 | 3 | 4 | 523 | 3 | 4 | 723 | 4 | 8 | 923 | 5 | 4 | 1123 | 6 | 6 |
| 124 | 4 | 5 | 324 | 3 | 4 | 524 | 7 | 8 | 724 | 6 | 6 | 924 | 8 | 9 | 1124 | 9 | 9 |
| 125 | 6 | 5 | 325 | 1 | 3 | 525 | 4 | 4 | 725 | 8 | 8 | 925 | 4 | 4 | 1125 | 7 | 7 |
| 126 | 4 | 5 | 326 | 4 | 5 | 526 | 6 | 8 | 726 | 4 | 4 | 926 | 6 | 9 | 1126 | 9 | 9 |
| 127 | 4 | 5 | 327 | 3 | 5 | 527 | 7 | 7 | 727 | 4 | 6 | 927 | 9 | 9 | 1127 | 9 | 9 |
| 128 | 4 | 4 | 328 | 5 | 6 | 528 | 8 | 9 | 728 | 4 | 6 | 928 | 7 | 8 | 1128 | 7 | 8 |
| 129 | 2 | 4 | 329 | 3 | 5 | 529 | 7 | 9 | 729 | 6 | 5 | 929 | 5 | 6 | 1129 | 8 | 9 |
| 130 | 7 | 6 | 330 | 4 | 5 | 530 | 8 | 8 | 730 | 7 | 8 | 930 | 6 | 8 | 1130 | 8 | 8 |
| 131 | 4 | 5 | 331 | 2 | 5 | 531 | 7 | 7 | 731 | 8 | 9 | 931 | 9 | 9 |  |  |  |
| 132 | 3 | 4 | 332 | 4 | 5 | 532 | 7 | 6 | 732 | 5 | 8 | 932 | 4 | 4 |  |  |  |
| 133 | 8 | 7 | 333 | 6 | 7 | 533 | 4 | 4 | 733 | 7 | 9 | 933 | 3 | 5 |  |  |  |
| 134 | 3 | 4 | 334 | 3 | 4 | 534 | 8 | 9 | 734 | 5 | 8 | 934 | 8 | 9 |  |  |  |
| 135 | 4 | 5 | 335 | 7 | 7 | 535 | 6 | 6 | 735 | 7 | 9 | 935 | 9 | 8 |  |  |  |
| 136 | 4 | 5 | 336 | 5 | 6 | 536 | 6 | 6 | 736 | 6 | 9 | 936 | 4 | 6 |  |  |  |
| 137 | 5 | 6 | 337 | 3 | 4 | 537 | 5 | 8 | 737 | 7 | 9 | 937 | 7 | 8 |  |  |  |
| 138 | 3 | 5 | 338 | 9 | 9 | 538 | 9 | 8 | 738 | 6 | 9 | 938 | 8 | 7 |  |  |  |
| 139 | 4 | 5 | 339 | 5 | 5 | 539 | 4 | 6 | 739 | 6 | 9 | 939 | 4 | 8 |  |  |  |
| 140 | 5 | 5 | 340 | 4 | 4 | 540 | 9 | 9 | 740 | 6 | 9 | 940 | 3 | 5 |  |  |  |
| 141 | 6 | 6 | 341 | 7 | 7 | 541 | 9 | 9 | 741 | 6 | 9 | 941 | 5 | 5 |  |  |  |
| 142 | 6 | 6 | 342 | 1 | 3 | 542 | 9 | 9 | 742 | 7 | 9 | 942 | 3 | 6 |  |  |  |
| 143 | 7 | 8 | 343 | 1 | 4 | 543 | 6 | 8 | 743 | 8 | 9 | 943 | 4 | 5 |  |  |  |
| 144 | 9 | 9 | 344 | 4 | 5 | 544 | 9 | 9 | 744 | 8 | 9 | 944 | 5 | 6 |  |  |  |
| 145 | 9 | 9 | 345 | 8 | 9 | 545 | 7 | 7 | 745 | 7 | 8 | 945 | 2 | 4 |  |  |  |
| 146 | 3 | 3 | 346 | 5 | 5 | 546 | 8 | 7 | 746 | 5 | 9 | 946 | 4 | 5 |  |  |  |
| 147 | 6 | 6 | 347 | 6 | 6 | 547 | 6 | 8 | 747 | 4 | 9 | 947 | 3 | 4 |  |  |  |
| 148 | 6 | 5 | 348 | 3 | 4 | 548 | 9 | 8 | 748 | 7 | 9 | 948 | 4 | 5 |  |  |  |
| 149 | 5 | 7 | 349 | 5 | 7 | 549 | 7 | 7 | 749 | 8 | 9 | 949 | 1 | 3 |  |  |  |
| 150 | 5 | 7 | 350 | 2 | 3 | 550 | 7 | 7 | 750 | 6 | 9 | 950 | 4 | 4 |  |  |  |
| 151 | 5 | 5 | 351 | 5 | 5 | 551 | 6 | 7 | 751 | 7 | 9 | 951 | 8 | 9 |  |  |  |
| 152 | 5 | 5 | 352 | 3 | 5 | 552 | 7 | 9 | 752 | 6 | 6 | 952 | 9 | 9 |  |  |  |
| 153 | 1 | 3 | 353 | 6 | 5 | 553 | 7 | 7 | 753 | 5 | 5 | 953 | 9 | 9 |  |  |  |
| 154 | 8 | 8 | 354 | 4 | 4 | 554 | 5 | 5 | 754 | 5 | 6 | 954 | 7 | 6 |  |  |  |
| 155 | 2 | 4 | 355 | 4 | 5 | 555 | 7 | 6 | 755 | 8 | 9 | 955 | 5 | 7 |  |  |  |
| 156 | 9 | 9 | 356 | 3 | 4 | 556 | 5 | 7 | 756 | 8 | 8 | 956 | 5 | 7 |  |  |  |
| 157 | 7 | 7 | 357 | 5 | 5 | 557 | 7 | 9 | 757 | 8 | 9 | 957 | 9 | 9 |  |  |  |
| 158 | 6 | 5 | 358 | 4 | 4 | 558 | 7 | 8 | 758 | 8 | 9 | 958 | 6 | 7 |  |  |  |
| 159 | 7 | 7 | 359 | 4 | 4 | 559 | 8 | 9 | 759 | 4 | 5 | 959 | 6 | 8 |  |  |  |
| 160 | 5 | 5 | 360 | 6 | 4 | 560 | 8 | 8 | 760 | 9 | 9 | 960 | 8 | 9 |  |  |  |
| 161 | 7 | 4 | 361 | 7 | 7 | 561 | 9 | 8 | 761 | 8 | 9 | 961 | 7 | 9 |  |  |  |
| 162 | 6 | 5 | 362 | 7 | 5 | 562 | 6 | 5 | 762 | 5 | 5 | 962 | 8 | 9 |  |  |  |
| 163 | 4 | 4 | 363 | 5 | 5 | 563 | 5 | 6 | 763 | 8 | 9 | 963 | 8 | 9 |  |  |  |
| 164 | 4 | 5 | 364 | 3 | 5 | 564 | 5 | 5 | 764 | 8 | 9 | 964 | 8 | 9 |  |  |  |
| 165 | 3 | 4 | 365 | 8 | 9 | 565 | 9 | 8 | 765 | 7 | 9 | 965 | 8 | 9 |  |  |  |
| 166 | 2 | 4 | 366 | 5 | 5 | 566 | 4 | 5 | 766 | 3 | 4 | 966 | 6 | 8 |  |  |  |
| 167 | 6 | 7 | 367 | 6 | 5 | 567 | 3 | 4 | 767 | 4 | 5 | 967 | 2 | 4 |  |  |  |
| 168 | 8 | 8 | 368 | 5 | 6 | 568 | 7 | 7 | 768 | 1 | 4 | 968 | 1 | 4 |  |  |  |
| 169 | 5 | 5 | 369 | 5 | 6 | 569 | 9 | 9 | 769 | 2 | 6 | 969 | 8 | 9 |  |  |  |
| 170 | 7 | 7 | 370 | 6 | 8 | 570 | 9 | 9 | 770 | 5 | 9 | 970 | 5 | 6 |  |  |  |
| 171 | 5 | 7 | 371 | 6 | 8 | 571 | 3 | 4 | 771 | 4 | 6 | 971 | 5 | 8 |  |  |  |
| 172 | 5 | 6 | 372 | 6 | 5 | 572 | 3 | 4 | 772 | 5 | 7 | 972 | 6 | 7 |  |  |  |
| 173 | 6 | 5 | 373 | 5 | 5 | 573 | 8 | 9 | 773 | 4 | 9 | 973 | 8 | 9 |  |  |  |
| 174 | 4 | 5 | 374 | 6 | 6 | 574 | 6 | 6 | 774 | 3 | 4 | 974 | 5 | 8 |  |  |  |
| 175 | 5 | 5 | 375 | 3 | 5 | 575 | 6 | 5 | 775 | 4 | 5 | 975 | 5 | 5 |  |  |  |
| 176 | 4 | 4 | 376 | 9 | 9 | 576 | 4 | 5 | 776 | 2 | 4 | 976 | 9 | 9 |  |  |  |
| 177 | 4 | 5 | 377 | 5 | 7 | 577 | 7 | 8 | 777 | 3 | 5 | 977 | 6 | 8 |  |  |  |
| 178 | 1 | 3 | 378 | 5 | 7 | 578 | 4 | 4 | 778 | 1 | 4 | 978 | 3 | 4 |  |  |  |
| 179 | 3 | 5 | 379 | 8 | 8 | 579 | 3 | 3 | 779 | 3 | 4 | 979 | 6 | 8 |  |  |  |
| 180 | 4 | 4 | 380 | 8 | 8 | 580 | 3 | 3 | 780 | 3 | 4 | 980 | 8 | 9 |  |  |  |
| 181 | 3 | 4 | 381 | 8 | 7 | 581 | 7 | 6 | 781 | 1 | 3 | 981 | 4 | 5 |  |  |  |
| 182 | 3 | 5 | 382 | 6 | 9 | 582 | 8 | 8 | 782 | 3 | 4 | 982 | 7 | 9 |  |  |  |
| 183 | 5 | 5 | 383 | 6 | 8 | 583 | 4 | 5 | 783 | 3 | 5 | 983 | 8 | 9 |  |  |  |
| 184 | 4 | 5 | 384 | 7 | 7 | 584 | 1 | 3 | 784 | 6 | 6 | 984 | 5 | 8 |  |  |  |
| 185 | 3 | 3 | 385 | 4 | 5 | 585 | 2 | 4 | 785 | 9 | 9 | 985 | 8 | 9 |  |  |  |
| 186 | 4 | 4 | 386 | 6 | 8 | 586 | 6 | 6 | 786 | 6 | 5 | 986 | 9 | 9 |  |  |  |
| 187 | 4 | 4 | 387 | 5 | 6 | 587 | 1 | 4 | 787 | 6 | 7 | 987 | 7 | 9 |  |  |  |
| 188 | 5 | 5 | 388 | 4 | 5 | 588 | 3 | 4 | 788 | 5 | 5 | 988 | 9 | 9 |  |  |  |
| 189 | 4 | 5 | 389 | 7 | 9 | 589 | 3 | 4 | 789 | 6 | 9 | 989 | 9 | 9 |  |  |  |
| 190 | 4 | 5 | 390 | 7 | 7 | 590 | 1 | 4 | 790 | 5 | 5 | 990 | 9 | 9 |  |  |  |
| 191 | 4 | 4 | 391 | 7 | 8 | 591 | 2 | 4 | 791 | 5 | 6 | 991 | 9 | 9 |  |  |  |
| 192 | 3 | 4 | 392 | 4 | 4 | 592 | 3 | 5 | 792 | 7 | 9 | 992 | 8 | 9 |  |  |  |
| 193 | 4 | 5 | 393 | 4 | 6 | 593 | 3 | 5 | 793 | 7 | 9 | 993 | 8 | 8 |  |  |  |
| 194 | 4 | 5 | 394 | 3 | 4 | 594 | 8 | 8 | 794 | 4 | 6 | 994 | 7 | 9 |  |  |  |
| 195 | 4 | 4 | 395 | 5 | 5 | 595 | 5 | 5 | 795 | 6 | 8 | 995 | 8 | 9 |  |  |  |
| 196 | 3 | 4 | 396 | 3 | 7 | 596 | 2 | 4 | 796 | 8 | 9 | 996 | 7 | 8 |  |  |  |
| 197 | 4 | 5 | 397 | 5 | 6 | 597 | 5 | 9 | 797 | 5 | 8 | 997 | 9 | 9 |  |  |  |
| 198 | 3 | 4 | 398 | 6 | 6 | 598 | 7 | 8 | 798 | 5 | 6 | 998 | 6 | 9 |  |  |  |
| 199 | 3 | 5 | 399 | 7 | 5 | 599 | 4 | 4 | 799 | 4 | 7 | 999 | 9 | 9 |  |  |  |
| 200 | 2 | 3 | 400 | 7 | 8 | 600 | 1 | 4 | 800 | 2 | 4 | 1000 | 9 | 9 |  |  |  |

**Supplementary figure 8 (next page).** **Changes in disorder probabilities predicted by PONDR for single amino acid substitutions.** **(a)** Disorder probabilities for WT CIITA sequence positions 1-160; red and blue vertical lines mark the experimentally-studied positions. **(b)‑(h)** At each of the experimentally-tested positions, substituitons with all 19 other amino acids were made computation­ally and disorder probabilities were calculated with PONDR. Plots are shown for the region containing CIITA 42-112; no changes in intrinsic propensity were observed outside of the region shown. In these panels, the propensities for the WT sequence is indicated with black dashed lines; the substituted position is indicated with the verticle line (**(e)** 62, **(f)** 65, **(g)** 70, **(b)**74, **(c)** 77, **(d)** 78, **(h)** 91); and colours for each of the probability curves correspond to the substituted amino acid type: acidic (D, E; red), basic (R, H, K; blue), polar uncharged (S, T, N, C, Q; yellow), hydrophobic (M, F, W, Y, A, V, I, L; green), and structure breakers (P, G; gray). The horizontal dotted lines on each panel mark the signficance threshold of 0.5.

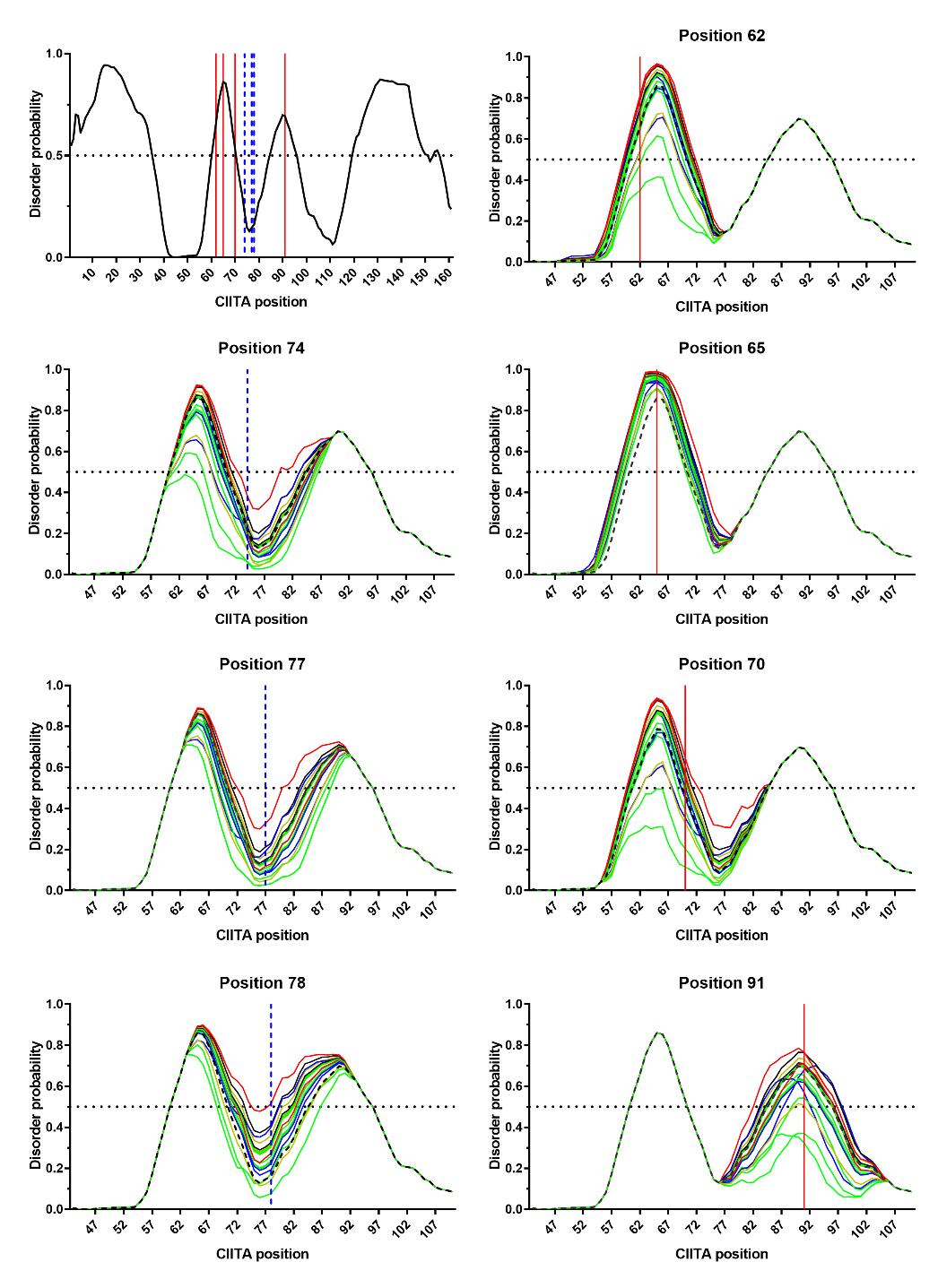

**(g)**

**(c)**

**(d)**

**(h)**

**(b)**

**(f)**

**(e)**

**(a)**

**Supplementary figure 9. Substitution-induced changes in disorder predictions are not unique to the CIITA region or to PONDR calculations**. On each panel, the horizontal dotted lines mark the signficance threshold of 0.5 and propensities for the substituted sequences are coloured as in Supplementary figure 8. Disorder propensities of the WT sequence are indicated by black dashed lines. No changes in intrinsic propensity were observed outside the regions shown on the x axes. **(a)** Predictions using PONDR’s VL-XT subroutine for all 19 possible amino acid substitutions at position 142, which is located outside of the experimentally-tested region. This control PONDR calculation shows that the substitution sensitivity of intrinsic propensity calculations is not a special property of the experimentally-tested CIITA region. **(b-c)** Predictions using metapredictV2^6^ for all 19 possible substitutions at positions **(b)** 62 and **(c)** 77, which were experimentally tested in the current work. Computations with metapredictV2 affect a wider sequence window than PONDR, which is expected from the different mathematical windows used by the two algorithms. These control computations illustrate that the sensitivity of intrinsic propensity computations to single substitutions is not unique to PONDR (Supplementary figure 8).

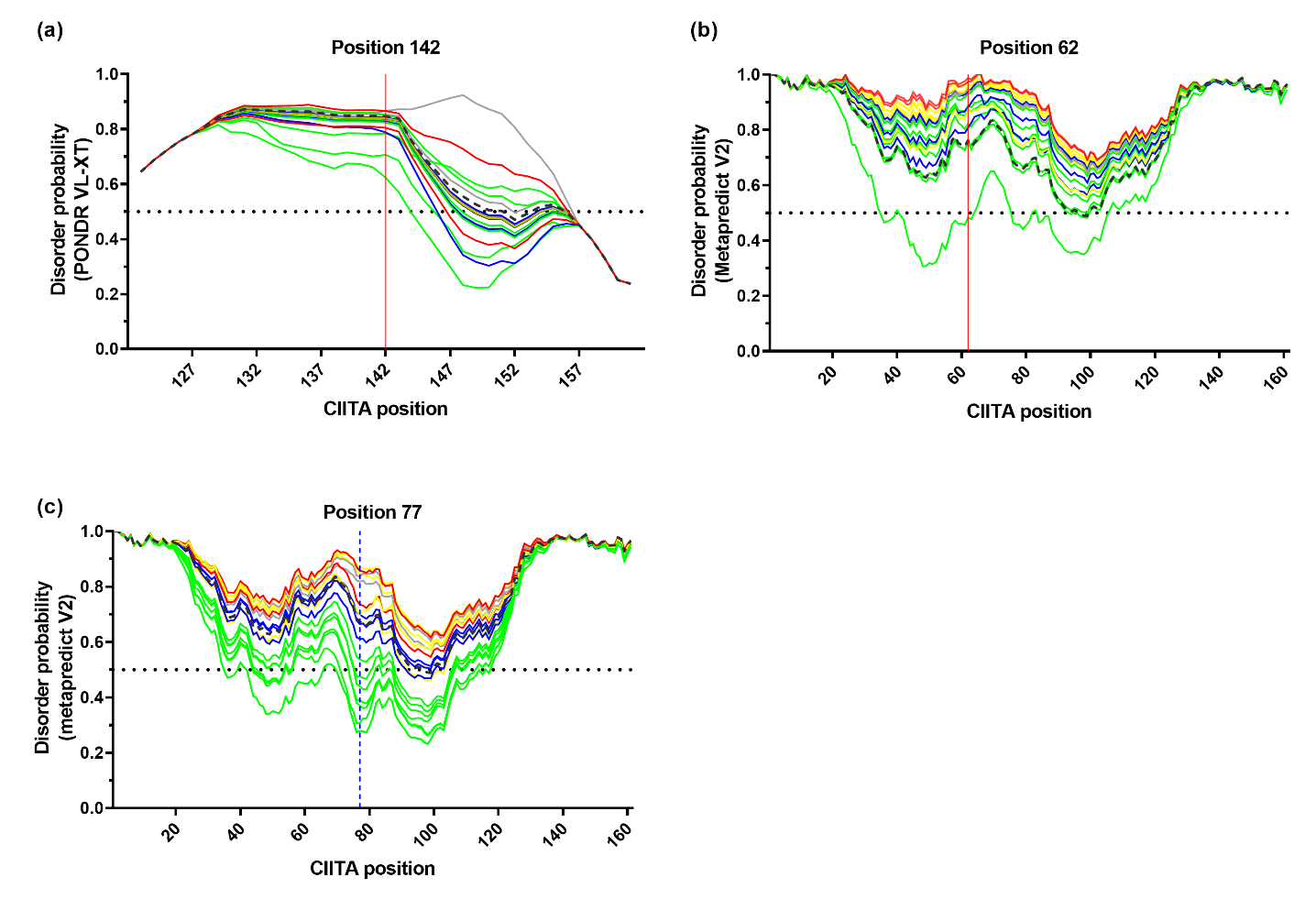

**Supplementary figure 10. Comparison of CIITA cellular concentration and side chain size**. For each of the substitutions experimentally assessed in this study, the side chain size was obtained from the relative, solvent-accessible amino acid side chain surface area in Å^2^ in a G-X-G tripeptide, where X is the amino acid ^9^. These values were compared to cellular concentrations for the assessed substitutions at positions with low (<0.5) **(a)** 74, **(b)** 77, **(c)** 78, and high disorder probabilities for the WT residue (>0.5) **(d)** 62, **(e)** 65, **(f)** 70, **(g)** 91.

**(d)**

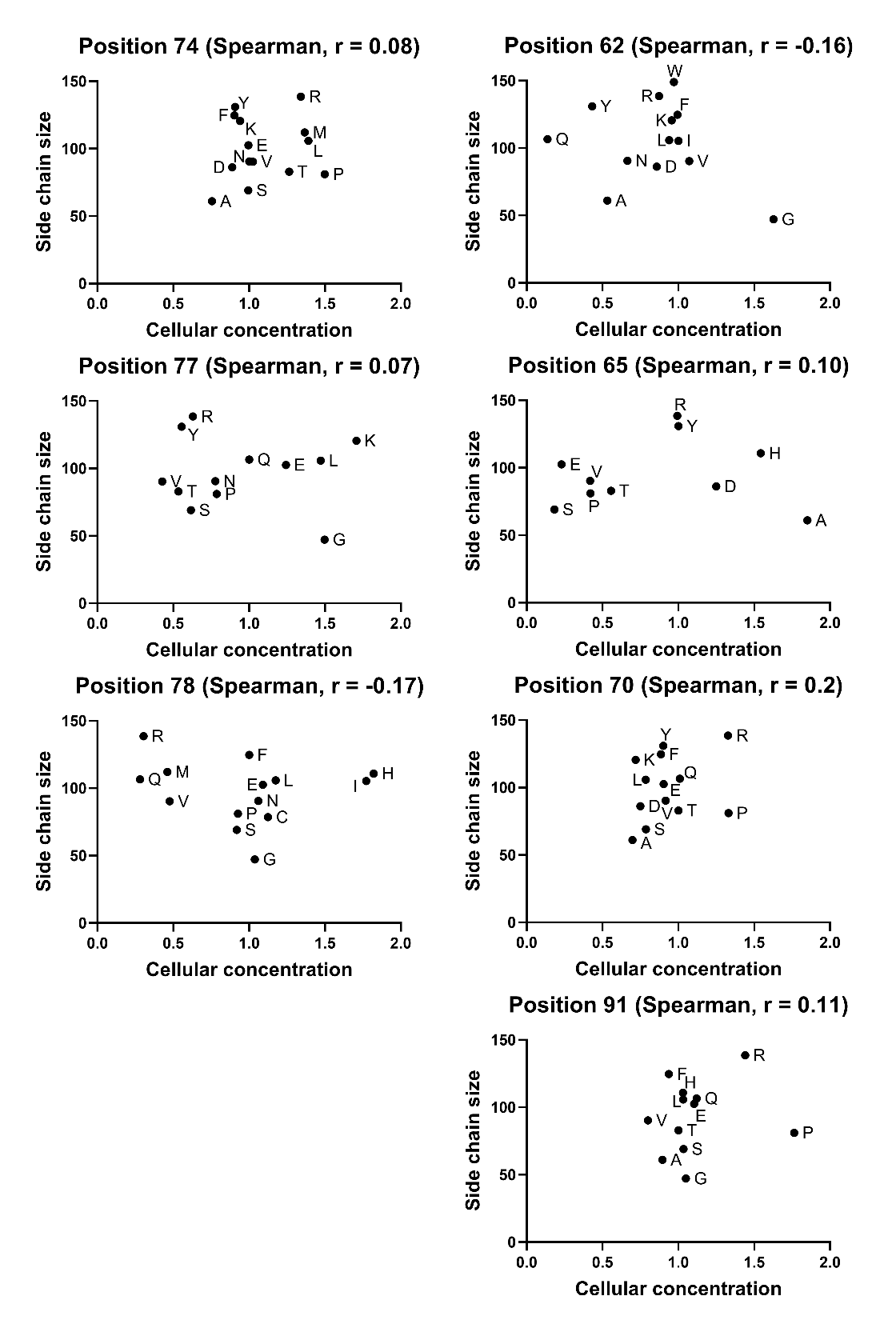

**(a)**

**(g)**

**(f)**

**(c)**

**(e)**

**(b)**

**Supplementary figure 11.** **Comparison of CIITA cellular concentration and helical propensity.** Helical propensities obtained host-guest peptide studies are reported in units of kcal/mol ^10^; lower values correspond to higher propensities. For each of the substitutions experimentally assessed in this study, helical propensities were compared to cellular concentration for positions with low (<0.5) **(a)** 74, **(b)** 77, **(c)** 78, and high disorder probabilities for the WT residue (>0.5) **(d)** 62, **(e)** 65, **(f)** 70, **(g)** 91.

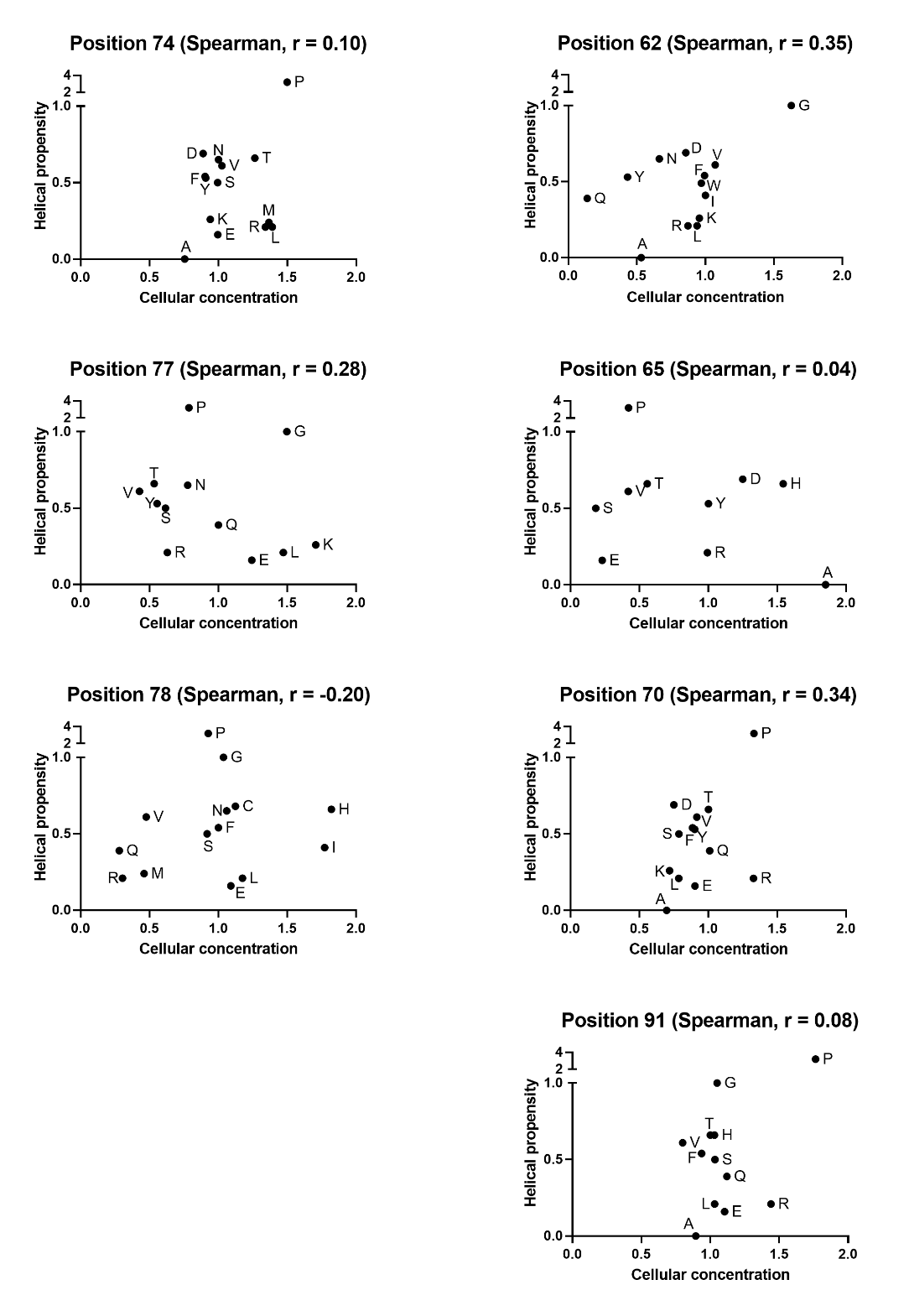

**(a)**

**(b)**

**(c)**

**(g)**

**(f)**

**(e)**

**(d)**

**Supplementary figure 12.** **Comparison of CIITA cellular concentration and disorder probabilities from PONDR**. PONDR scores were taken from the plots shown in Supplementary figure 8. For each of the substitutions assessed in this study, these values were compared to cellular concentration for positions with low (<0.5) **(a)** 74, **(b)** 77, **(c)** 78, and high disorder probabilities for the WT residue (>0.5) **(d)** 62, **(e)** 65, **(f)** 70, **(g)** 91.

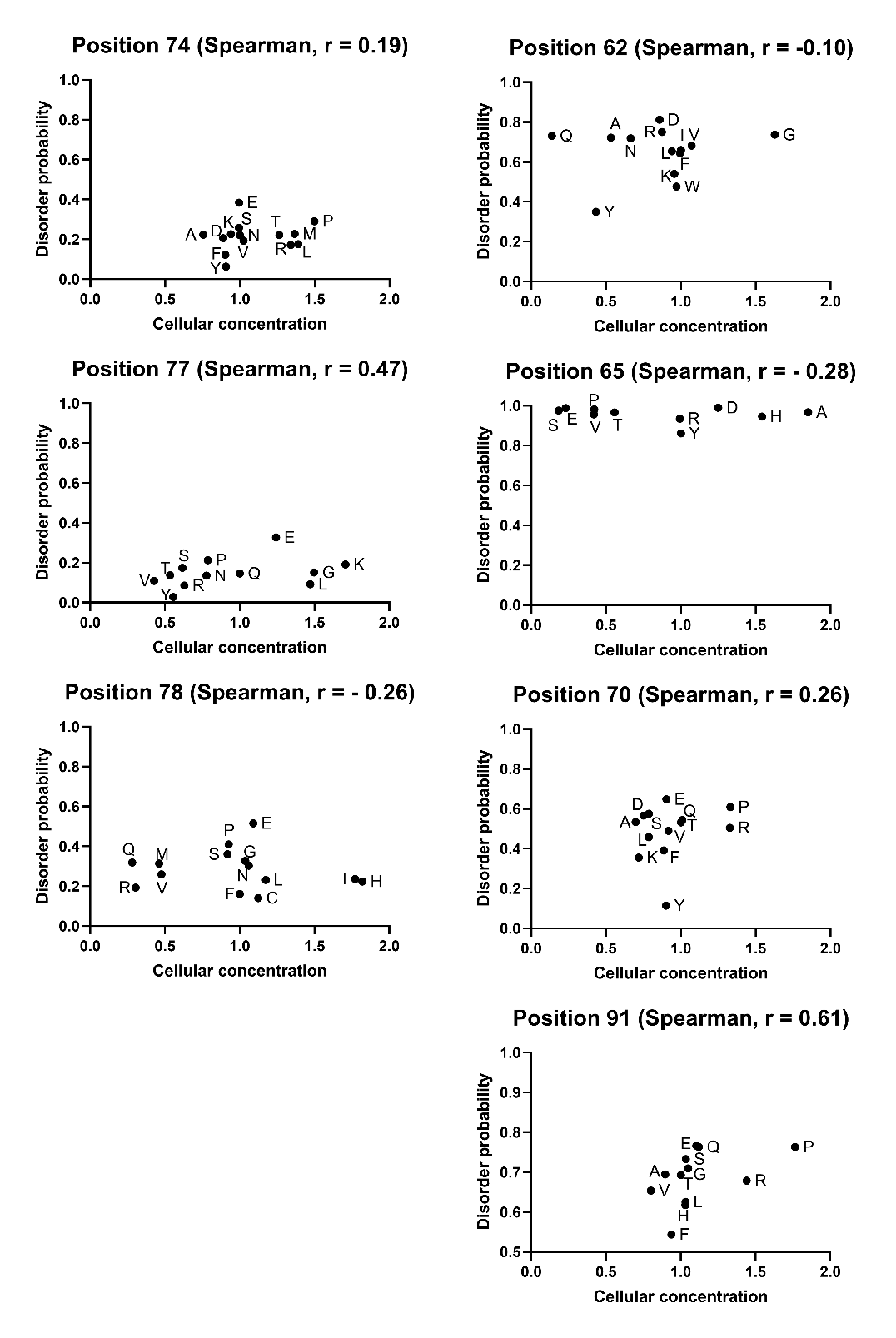

**(d)**

**(a)**

**(g)**

**(f)**

**(c)**

**(e)**

**(b)**

**Supplementary figure 13.** **Comparison of CIITA transcriptional activation and amino acid side chain size.** For each of the substitutions experimentally assessed in this study, the side chain size was obtained from a the relative, solvent-accessible amino acid side chain surface area in Å^2^ in a G-X-G tripeptide ^9^. These values were compared to transcriptional activation for the assessed substitutions at positions with low (<0.5) **(a)** 74, **(b)** 77 **(c)** 78, and high disorder probabilities for the WT residue (>0.5) **(d)** 62, **(e)** 65, **(f)** 70, **(g)** 91.

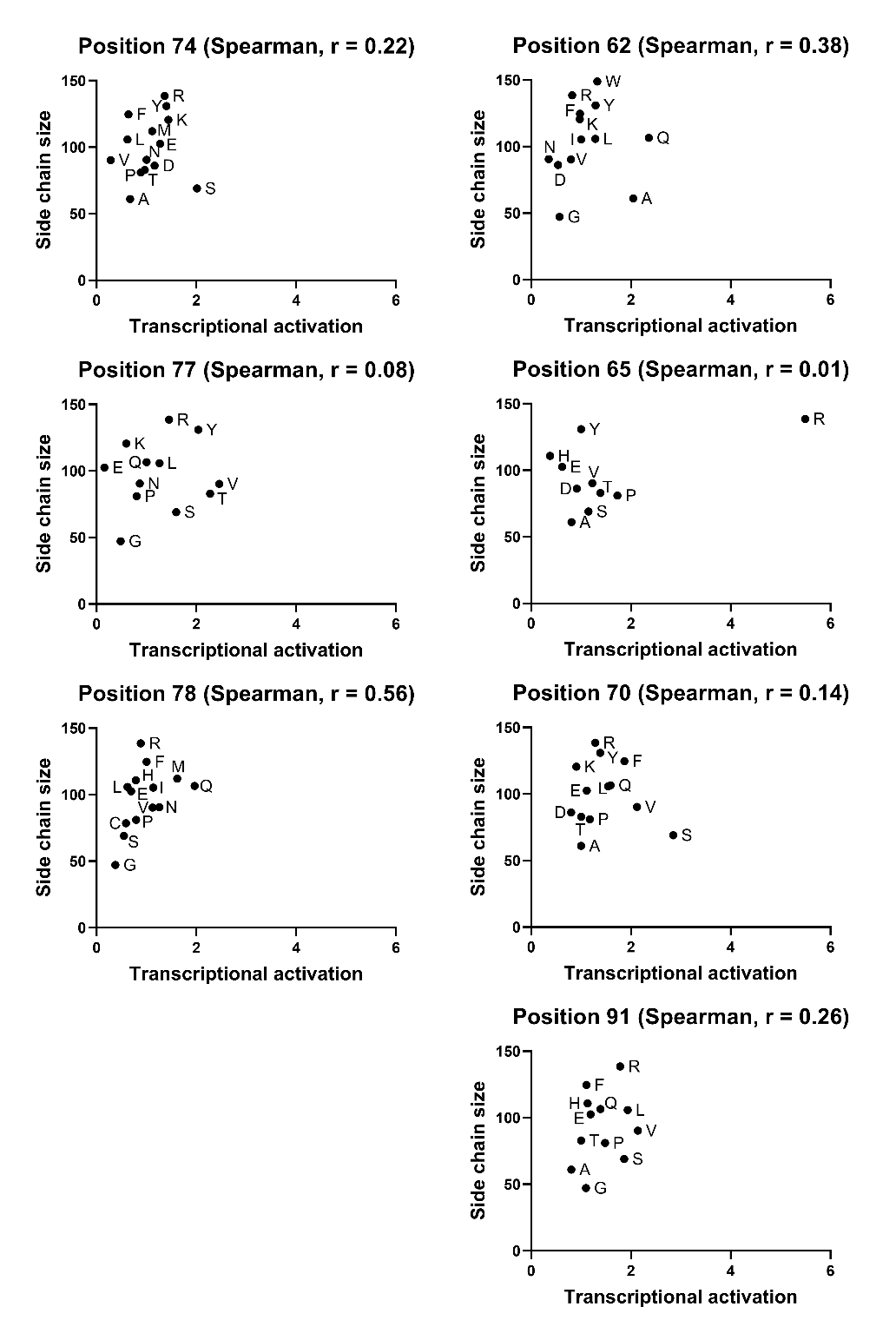

**(d)**

**(a)**

**(g)**

**(e)**

**(f)**

**(c)**

**(b)**

**Supplementary figure 14.** **Comparison of CIITA transcriptional activation by CIITA and helical propensity.** Helical propensities obtained from host-guest peptide studies are reported in units of kcal/mol ^10^; lower values correspond to higher propensities. For each of the substitutions experimentally assessed in this study, helical propensities were compared to transcriptional activation for positions with low (<0.5) **(a)** 74, **(b)** 77, **(c)** 78, and high disorder probabilities for the WT residue (>0.5) **(d)** 62, **(e)** 65, **(f)** 70, **(g)** 91.

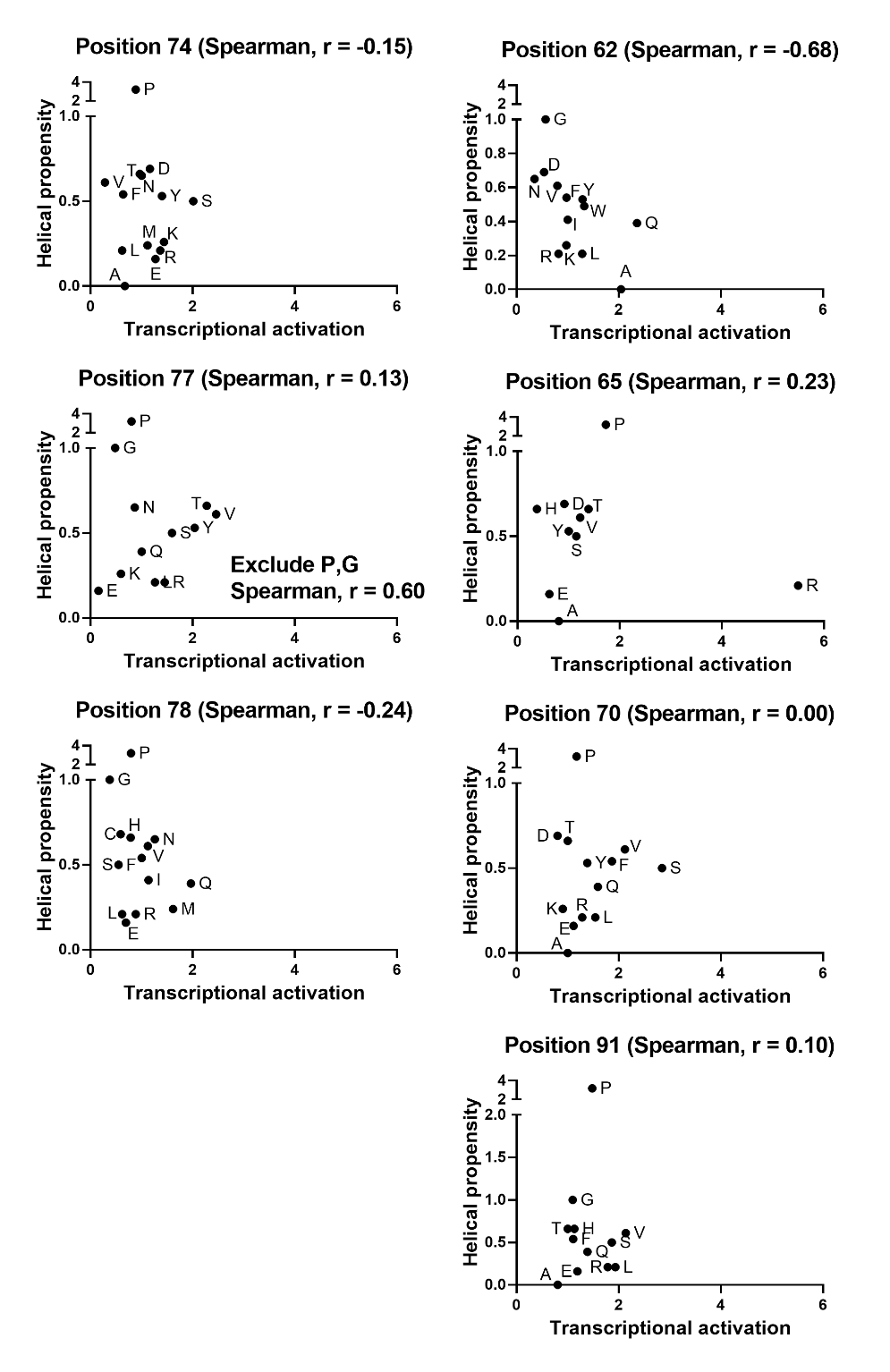

**(d)**

**(a)**

**(g)**

**(f)**

**(e)**

**(c)**

**(b)**

**Supplementary figure 15. Comparison of CIITA transcriptional activation by CIITA and disorder probabilities from PONDR and metapredictV2. (a-b, d-f)** PONDR scores were taken from the plots shown in Supplementary figure 8. For each of the substitutions assessed in this study, the values were compared to transcriptional activation for positions with low (<0.5) **(a)** 74, **(b)** 77, **(d)** 78, and high disorder probabilities for the WT residue (>0.5) **(e)** 62, **(f)** 65, **(g)** 70, **(h)** 91. In **(c),** the experimentally determined transcriptional activation levels were compared to metapredictV2’s propensities for experimentally-assessed substitutions (Supplementary figure 9c). The similarity of panels b and c suggests the correlation is not an artefact of PONDR.

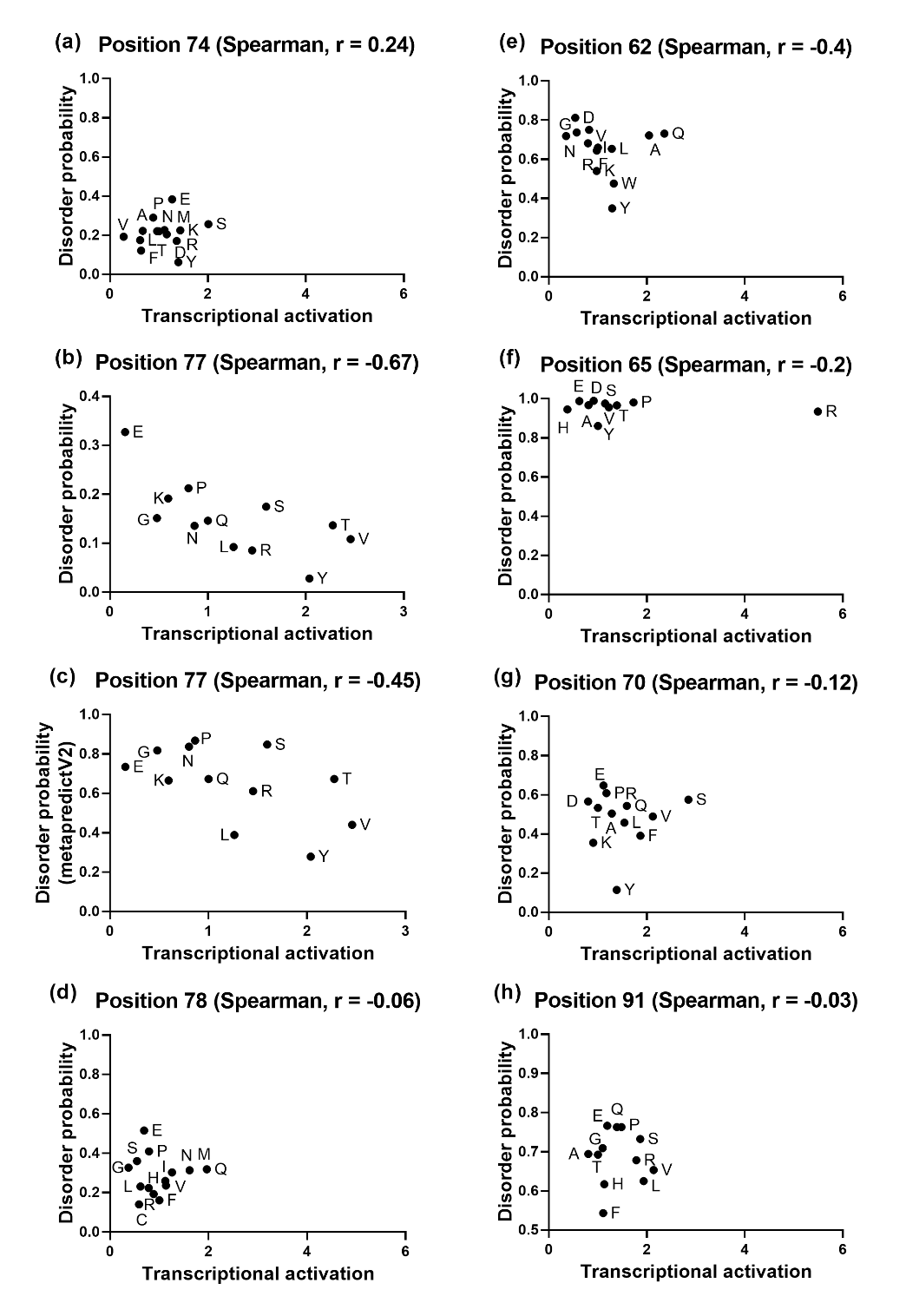

**Supplementary figure 16**. **Correlation of ConSurf raw scores for various CIITA MSAs and MSA subsets.** ConSurf calculates the relative evolutionary conservation score at each position in an MSA and then Z-normalizes all scores to the average score. The lowest score is assigned to the most conserved position, and the highest score is assigned to the most variable position. **(a)-(i)** Nine representative correlations for ConSurf raw scores of 19 random sequence subsets extracted from “ConSurf_manual_MSA” with corresponding Spearman coefficients. Agreement between the scores (dots along the diagonal) shows that the subsets adequately sample the CIITA family; since subsamples with 250 sequences showed less agreement, average ConSurf values for this MSA were calculated from subsamples containing >300 CIITA sequences.

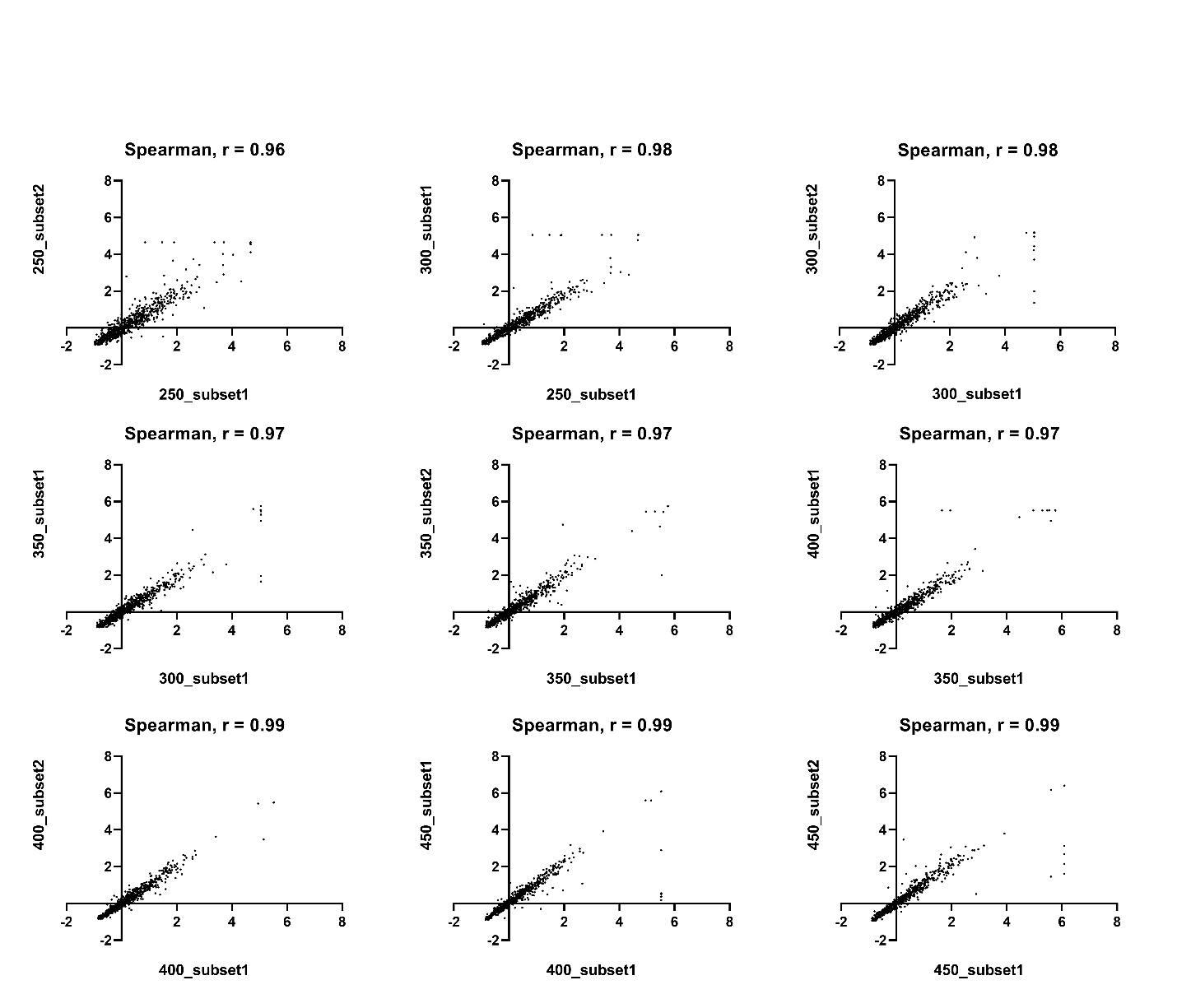

**(c)**

**(b)**

**(a)**

**(f)**

**(e)**

**(d)**

**(h)**

**(g)**

**(i)**

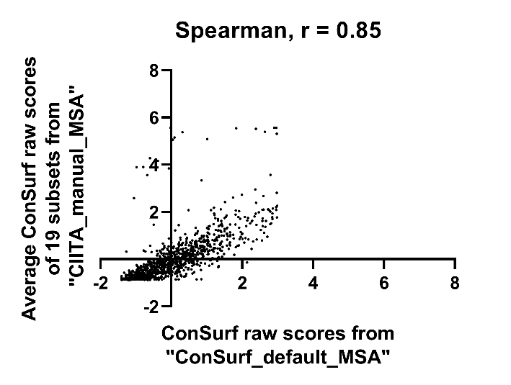
**(j)** ConSurf raw scores were averaged from 19 subsets of “CIITA_manual_MSA” and plotted against raw scores determined for “CIITA_default_MSA”. The two score sets show reasonable agreement across most positions, suggesting that the two MSAs are comparable. The values for these scores are in Supplementary table 2. Most of the observed differences are in the first 20 positions of CIITA.

**Supplementary table 5.** **One way ANOVA.** Statistical analyses for measured experimental outcomes for amino acid variants relative to the WT sample measured in parallel. **(a)**-**(g)** cellular concentration **(h)-(n)** transcriptional activation **(o)-(u)** combined effects.

**Supplementary table 5a. Position 62 (Cellular CIITA concentration)**

| Dunnett's multiple  comparisons test | Mean Diff. | 95.00% CI of diff. | Summary | Adjusted P Value |
| --- | --- | --- | --- | --- |
| WT-I62 vs. I62G | -0.159 | -0.3511 to 0.03319 | ns | 0.178 |
| WT-I62 vs. I62V | -0.02454 | -0.2382 to 0.1891 | ns | 0.9996 |
| WT-I62 vs. I62D | 0.09218 | -0.1095 to 0.2938 | ns | 0.873 |
| WT-I62 vs. I62R | 0.04965 | -0.2016 to 0.3009 | ns | 0.9993 |
| WT-I62 vs. I62F | 0.02118 | -0.1710 to 0.2133 | ns | 0.9997 |
| WT-I62 vs. I62K | 0.01762 | -0.2336 to 0.2688 | ns | 0.9998 |
| WT-I62 vs. I62L | 0.01929 | -0.2319 to 0.2705 | ns | 0.9997 |
| WT-I62 vs. I62Y | 0.4707 | 0.2691 to 0.6724 | **** | <0.0001 |
| WT-I62 vs. I62W | 0.08644 | -0.1057 to 0.2786 | ns | 0.8837 |
| WT-I62 vs. I62A | 0.4291 | 0.2369 to 0.6213 | **** | <0.0001 |
| WT-I62 vs. I62Q | 0.8519 | 0.6006 to 1.103 | **** | <0.0001 |

**Supplementary table 5b. Position 65 (Cellular CIITA concentration)**

| Dunnett's multiple comparisons test | Mean Diff. | 95.00% CI of diff. | Summary | Adjusted P Value |
| --- | --- | --- | --- | --- |
| WT - Y65 vs. Y65E | 0.6392 | 0.3304 to 0.9480 | **** | <0.0001 |
| WT - Y65 vs. Y65P | 0.4884 | 0.1921 to 0.7848 | *** | 0.0002 |
| WT - Y65 vs. Y65A | -0.2558 | -0.5522 to 0.04054 | ns | 0.1239 |
| WT - Y65 vs. Y65H | -0.1224 | -0.4471 to 0.2023 | ns | 0.8966 |
| WT - Y65 vs. Y65D | -0.08845 | -0.3848 to 0.2079 | ns | 0.9689 |
| WT - Y65 vs. Y65R | 0.03609 | -0.2886 to 0.3608 | ns | 0.9996 |
| WT - Y65 vs. Y65S | 0.7642 | 0.4396 to 1.089 | **** | <0.0001 |
| WT - Y65 vs. Y65T | 0.3316 | 0.02276 to 0.6404 | * | 0.0293 |

**Supplementary table 5c. Position 70 (Cellular CIITA concentration)**

| Dunnett's multiple comparisons test | Mean Diff. | 95.00% CI of diff. | Summary | Adjusted P Value |
| --- | --- | --- | --- | --- |
| WT - T70 vs. T70D | 0.1984 | 0.03992 to 0.3568 | ** | 0.0048 |
| WT - T70 vs. T70K | 0.153 | 0.01136 to 0.2947 | * | 0.025 |
| WT - T70 vs. T70E | 0.04412 | -0.09754 to 0.1858 | ns | 0.9914 |
| WT - T70 vs. T70A | 0.1577 | 0.01608 to 0.2994 | * | 0.0185 |
| WT - T70 vs. T70P | -0.07148 | -0.2131 to 0.07018 | ns | 0.8139 |
| WT - T70 vs. T70R | -0.105 | -0.2467 to 0.03666 | ns | 0.3118 |
| WT - T70 vs. T70Y | 0.1046 | -0.03705 to 0.2463 | ns | 0.3169 |
| WT - T70 vs. T70L | 0.1378 | -0.003870 to 0.2795 | ns | 0.0626 |
| WT - T70 vs. T70Q | 0.01565 | -0.1260 to 0.1573 | ns | 0.9997 |
| WT - T70 vs. T70F | 0.07238 | -0.08608 to 0.2308 | ns | 0.8914 |
| WT - T70 vs. T70V | 0.04363 | -0.1148 to 0.2021 | ns | 0.999 |
| WT - T70 vs. T70S | 0.1208 | -0.06682 to 0.3084 | ns | 0.5105 |

**Supplementary table 5d. Position 74 (Cellular CIITA concentration)**

| Dunnett's multiple comparisons test | Mean Diff. | 95.00% CI of diff. | Summary | Adjusted P Value |
| --- | --- | --- | --- | --- |
| WT - N74 vs. N74V | -0.0104 | -0.1450 to 0.1242 | Ns | 0.9998 |
| WT - N74 vs. N74L | -0.1443 | -0.2569 to -0.03168 | ** | 0.0033 |
| WT - N74 vs. N74F | 0.0506 | -0.06202 to 0.1632 | Ns | 0.9211 |
| WT - N74 vs. N74A | 0.144 | 0.03138 to 0.2566 | ** | 0.0034 |
| WT - N74 vs. N74P | -0.1602 | -0.2454 to -0.07496 | **** | <0.0001 |
| WT - N74 vs. N74T | -0.0999 | -0.2 to -6.774e-005 | * | 0.0497 |
| WT - N74 vs. N74M | -0.1317 | -0.2231 to -0.04039 | *** | 0.0005 |
| WT - N74 vs. N74D | 0.05193 | -0.04792 to 0.1518 | ns | 0.8127 |
| WT - N74 vs. N74R | -0.0915 | -0.1767 to -0.006260 | * | 0.026 |
| WT - N74 vs. N74E | 0.02063 | -0.09199 to 0.1333 | ns | 0.9994 |
| WT - N74 vs. N74Y | 0.04963 | -0.05023 to 0.1495 | ns | 0.8537 |
| WT - N74 vs. N74K | 0.03557 | -0.07705 to 0.1482 | ns | 0.9924 |
| WT - N74 vs. N74S | 0.04624 | -0.04696 to 0.1394 | ns | 0.8552 |

**Supplementary table 5e. Position 77 (Cellular CIITA concentration)**

| Dunnett's multiple comparisons test | Mean Diff. | 95.00% CI of diff. | Summary | Adjusted P Value |
| --- | --- | --- | --- | --- |
| WT - Q77 vs. Q77E | -0.05972 | -0.3225 to 0.2030 | ns | 0.9991 |
| WT - Q77 vs. Q77G | -0.173 | -0.4492 to 0.1031 | ns | 0.5134 |
| WT - Q77 vs. Q77K | -0.1625 | -0.4253 to 0.1002 | ns | 0.5308 |
| WT - Q77 vs. Q77P | 0.1061 | -0.1567 to 0.3689 | ns | 0.9304 |
| WT - Q77 vs. Q77N | 0.1138 | -0.1490 to 0.3765 | ns | 0.8956 |
| WT - Q77 vs. Q77L | -0.1169 | -0.3796 to 0.1459 | ns | 0.8793 |
| WT - Q77 vs. Q77R | 0.286 | 0.02327 to 0.5488 | * | 0.0241 |
| WT - Q77 vs. Q77S | 0.2702 | -0.005959 to 0.5464 | ns | 0.0592 |
| WT - Q77 vs. Q77Y | 0.3293 | 0.05314 to 0.6055 | ** | 0.0096 |
| WT - Q77 vs. Q77T | 0.3334 | 0.07069 to 0.5962 | ** | 0.0047 |
| WT - Q77 vs. Q77V | 0.4317 | 0.1689 to 0.6944 | **** | <0.0001 |

**Supplementary table 5f. Position 78 (Cellular CIITA concentration)**

| Dunnett's multiple comparisons test | Mean Diff. | 95.00% CI of diff. | Summary | Adjusted P Value |
| --- | --- | --- | --- | --- |
| WT - F78 vs. F78S | 0.07881 | -0.09006 to 0.2477 | ns | 0.8764 |
| WT - F78 vs. F78C | -0.05095 | -0.2241 to 0.1221 | ns | 0.9932 |
| WT - F78 vs. F78L | -0.08922 | -0.2623 to 0.08388 | ns | 0.7922 |
| WT - F78 vs. F78E | -0.00803 | -0.1769 to 0.1608 | ns | 0.9999 |
| WT - F78 vs. F78H | -0.2364 | -0.4142 to -0.05861 | ** | 0.0021 |
| WT - F78 vs. F78R | 0.5383 | 0.3694 to 0.7072 | **** | <0.0001 |
| WT - F78 vs. F78P | 0.06067 | -0.1082 to 0.2295 | ns | 0.9789 |
| WT - F78 vs. F78I | -0.2042 | -0.3820 to -0.02634 | * | 0.0133 |
| WT - F78 vs. F78V | 0.3647 | 0.1958 to 0.5336 | **** | <0.0001 |
| WT - F78 vs. F78N | -0.01164 | -0.1847 to 0.1615 | ns | 0.9998 |
| WT - F78 vs. F78M | 0.326 | 0.1571 to 0.4948 | **** | <0.0001 |
| WT - F78 vs. F78Q | 0.6003 | 0.4314 to 0.7692 | **** | <0.0001 |

**Supplementary table 5g. Position 91 (Cellular CIITA concentration)**

| Dunnett's multiple comparisons test | Mean Diff. | 95.00% CI of diff. | Summary | Adjusted P Value |
| --- | --- | --- | --- | --- |
| WT - T91 vs. T91G | -0.03171 | -0.2298 to 0.1664 | ns | 0.9994 |
| WT - T91 vs. T91F | 0.04317 | -0.1549 to 0.2413 | ns | 0.9972 |
| WT - T91 vs. T91H | -0.0013 | -0.1994 to 0.1968 | ns | >0.9999 |
| WT - T91 vs. T91E | -0.03201 | -0.2301 to 0.1661 | ns | 0.9994 |
| WT - T91 vs. T91Q | -0.03549 | -0.2336 to 0.1626 | ns | 0.9994 |
| WT - T91 vs. T91P | -0.2057 | -0.4038 to -0.007655 | * | 0.0368 |
| WT - T91 vs. T91R | -0.1476 | -0.3457 to 0.05048 | ns | 0.2791 |
| WT - T91 vs. T91S | 0.01704 | -0.1810 to 0.2151 | ns | 0.9997 |
| WT - T91 vs. T91L | 0.0596 | -0.1385 to 0.2577 | ns | 0.9886 |
| WT - T91 vs. T91V | 0.12 | -0.07807 to 0.3181 | ns | 0.5456 |

**Supplementary table 5h. Position 62 (Transcriptional activation)**

| Dunnett's multiple comparisons test | Mean Diff. | 95.00% CI of diff. | Summary | Adjusted P Value |
| --- | --- | --- | --- | --- |
| WT-I62 vs. I62N | 0.4573 | 0.2725 to 0.6421 | **** | <0.0001 |
| WT-I62 vs. I62G | 0.2595 | 0.1048 to 0.4143 | **** | <0.0001 |
| WT-I62 vs. I62V | 0.127 | -0.04511 to 0.2990 | ns | 0.3152 |
| WT-I62 vs. I62D | 0.2867 | 0.1243 to 0.4491 | **** | <0.0001 |
| WT-I62 vs. I62R | 0.08209 | -0.1202 to 0.2844 | ns | 0.9466 |
| WT-I62 vs. I62F | 0.03683 | -0.1179 to 0.1916 | ns | 0.9991 |
| WT-I62 vs. I62K | 0.000832 | -0.2015 to 0.2031 | ns | >0.9999 |
| WT-I62 vs. I62L | -0.1069 | -0.3092 to 0.09542 | ns | 0.7631 |
| WT-I62 vs. I62Y | -0.09405 | -0.2610 to 0.07289 | ns | 0.6884 |
| WT-I62 vs. I62W | -0.08644 | -0.2448 to 0.07194 | ns | 0.7268 |
| WT-I62 vs. I62A | -0.2815 | -0.4362 to -0.1267 | **** | <0.0001 |
| WT-I62 vs. I62Q | -0.3813 | -0.5836 to -0.1790 | **** | <0.0001 |

**Supplementary table 5i. Position 65 (Transcriptional activation)**

| Dunnett's multiple comparisons test | Mean Diff. | 95.00% CI of diff. | Summary | Adjusted P Value |
| --- | --- | --- | --- | --- |
| WT - Y65 vs. Y65V | -0.1248 | -0.3977 to 0.1482 | ns | 0.8217 |
| WT - Y65 vs. Y65E | 0.1939 | -0.06365 to 0.4514 | ns | 0.2559 |
| WT - Y65 vs. Y65P | -0.2649 | -0.5102 to -0.01959 | * | 0.0269 |
| WT - Y65 vs. Y65A | 0.0589 | -0.1864 to 0.3042 | ns | 0.9962 |
| WT - Y65 vs. Y65H | 0.4196 | 0.1742 to 0.6649 | **** | <0.0001 |
| WT - Y65 vs. Y65D | 0.01866 | -0.2267 to 0.2640 | ns | 0.9997 |
| WT - Y65 vs. Y65R | -0.7852 | -1.058 to -0.5122 | **** | <0.0001 |
| WT - Y65 vs. Y65S | -0.06495 | -0.3379 to 0.2080 | ns | 0.9964 |
| WT - Y65 vs. Y65T | -0.1518 | -0.4093 to 0.1057 | ns | 0.5552 |

**Supplementary table 5j. Position 70 (Transcriptional activation)**

| Dunnett's multiple comparisons test | Mean Diff. | 95.00% CI of diff. | Summary | Adjusted P Value |
| --- | --- | --- | --- | --- |
| WT - T70 vs. T70D | 0.1593 | -0.05450 to 0.3731 | Ns | 0.3074 |
| WT - T70 vs. T70K | 0.07569 | -0.07577 to 0.2271 | Ns | 0.8261 |
| WT - T70 vs. T70E | 0.02153 | -0.1721 to 0.2151 | Ns | 0.9997 |
| WT - T70 vs. T70A | 0.004874 | -0.1553 to 0.1651 | Ns | >0.9999 |
| WT - T70 vs. T70P | -0.04649 | -0.2348 to 0.1418 | Ns | 0.9991 |
| WT - T70 vs. T70R | -0.09513 | -0.2486 to 0.05830 | Ns | 0.5686 |
| WT - T70 vs. T70Y | -0.03362 | -0.1964 to 0.1292 | Ns | 0.9993 |
| WT - T70 vs. T70L | -0.1438 | -0.3321 to 0.04450 | Ns | 0.2757 |
| WT - T70 vs. T70Q | -0.1766 | -0.3650 to 0.01168 | Ns | 0.0825 |
| WT - T70 vs. T70F | -0.2729 | -0.4867 to -0.05914 | ** | 0.0037 |
| WT - T70 vs. T70V | -0.3095 | -0.5157 to -0.1033 | *** | 0.0003 |
| WT - T70 vs. T70S | -0.4265 | -0.6717 to -0.1813 | **** | <0.0001 |

**Supplementary table 5k. Position 74 (Transcriptional activation)**

| Dunnett's multiple comparisons test | Mean Diff. | 95.00% CI of diff. | Summary | Adjusted P Value |
| --- | --- | --- | --- | --- |
| WT - N74 vs. N74V | 0.5329 | 0.2883 to 0.7776 | **** | <0.0001 |
| WT - N74 vs. N74L | 0.2666 | 0.05065 to 0.4825 | ** | 0.0053 |
| WT - N74 vs. N74F | 0.1998 | -0.01613 to 0.4157 | ns | 0.0925 |
| WT - N74 vs. N74A | 0.1678 | -0.05638 to 0.3919 | ns | 0.314 |
| WT - N74 vs. N74P | 0.07199 | -0.1033 to 0.2473 | ns | 0.9579 |
| WT - N74 vs. N74T | 0.04631 | -0.1453 to 0.2379 | ns | 0.9992 |
| WT - N74 vs. N74M | 0.02543 | -0.1499 to 0.2007 | ns | 0.9996 |
| WT - N74 vs. N74D | -0.02206 | -0.2187 to 0.1746 | ns | 0.9997 |
| WT - N74 vs. N74R | -0.06897 | -0.2410 to 0.1031 | ns | 0.9648 |
| WT - N74 vs. N74E | -0.04402 | -0.2599 to 0.1719 | ns | 0.9993 |
| WT - N74 vs. N74Y | -0.1517 | -0.3432 to 0.03991 | ns | 0.2408 |
| WT - N74 vs. N74K | -0.111 | -0.3269 to 0.1049 | ns | 0.8239 |
| WT - N74 vs. N74S | -0.2181 | -0.3970 to -0.03924 | ** | 0.0062 |

**Supplementary table 5l. Position 77 (Transcriptional activation)**

| Dunnett's multiple comparisons test | Mean Diff. | 95.00% CI of diff. | Summary | Adjusted P Value |
| --- | --- | --- | --- | --- |
| WT - Q77 vs. Q77E | 0.8114 | 0.5889 to 1.034 | **** | <0.0001 |
| WT - Q77 vs. Q77G | 0.3176 | 0.08371 to 0.5514 | ** | 0.002 |
| WT - Q77 vs. Q77K | 0.271 | 0.04846 to 0.4935 | ** | 0.0076 |
| WT - Q77 vs. Q77P | 0.214 | -0.008450 to 0.4365 | ns | 0.0671 |
| WT - Q77 vs. Q77N | 0.1108 | -0.1117 to 0.3333 | ns | 0.7897 |
| WT - Q77 vs. Q77L | -0.06003 | -0.2825 to 0.1625 | ns | 0.9958 |
| WT - Q77 vs. Q77R | -0.09686 | -0.3194 to 0.1256 | ns | 0.8925 |
| WT - Q77 vs. Q77S | -0.1866 | -0.4205 to 0.04727 | ns | 0.2088 |
| WT - Q77 vs. Q77Y | -0.3102 | -0.5441 to -0.07635 | ** | 0.0027 |
| WT - Q77 vs. Q77T | -0.3229 | -0.5454 to -0.1004 | *** | 0.0007 |
| WT - Q77 vs. Q77V | -0.3843 | -0.6068 to -0.1618 | **** | <0.0001 |

**Supplementary table 5m. Position 78 (Transcriptional activation)**

| Dunnett's multiple comparisons test | Mean Diff. | 95.00% CI of diff. | Summary | Adjusted P Value |
| --- | --- | --- | --- | --- |
| WT - F78 vs. F78G | 0.4111 | 0.2131 to 0.6092 | **** | <0.0001 |
| WT - F78 vs. F78S | 0.3036 | 0.08133 to 0.5258 | ** | 0.0013 |
| WT - F78 vs. F78C | 0.2252 | 0.02714 to 0.4232 | * | 0.0144 |
| WT - F78 vs. F78L | 0.2479 | 0.03914 to 0.4567 | ** | 0.0088 |
| WT - F78 vs. F78E | 0.1697 | -0.02834 to 0.3678 | ns | 0.1529 |
| WT - F78 vs. F78P | 0.09269 | -0.1054 to 0.2907 | ns | 0.8916 |
| WT - F78 vs. F78H | 0.08563 | -0.1295 to 0.3007 | ns | 0.9638 |
| WT - F78 vs. F78R | 0.08005 | -0.1287 to 0.2888 | ns | 0.9729 |
| WT - F78 vs. F78I | 0.01858 | -0.1902 to 0.2274 | ns | 0.9997 |
| WT - F78 vs. F78V | 0.02022 | -0.1778 to 0.2183 | ns | 0.9997 |
| WT - F78 vs. F78N | -0.04116 | -0.2443 to 0.1620 | ns | 0.9993 |
| WT - F78 vs. F78M | -0.2014 | -0.3995 to -0.003350 | * | 0.0432 |
| WT - F78 vs. F78Q | -0.3043 | -0.5194 to -0.08917 | *** | 0.0008 |

**Supplementary table 5n. Position 91 (Transcriptional activation)**

| Dunnett's multiple comparisons test | Mean Diff. | 95.00% CI of diff. | Summary | Adjusted P Value |
| --- | --- | --- | --- | --- |
| WT - T91 vs. T91A | 0.1735 | -0.03441 to 0.3813 | ns | 0.166 |
| WT - T91 vs. T91G | -0.02546 | -0.2333 to 0.1824 | ns | 0.9996 |
| WT - T91 vs. T91F | -0.02204 | -0.2299 to 0.1858 | ns | 0.9997 |
| WT - T91 vs. T91H | -0.05485 | -0.2627 to 0.1530 | ns | 0.9961 |
| WT - T91 vs. T91E | -0.07868 | -0.2865 to 0.1292 | ns | 0.9543 |
| WT - T91 vs. T91Q | -0.1265 | -0.3344 to 0.08137 | ns | 0.5547 |
| WT - T91 vs. T91P | -0.1656 | -0.3735 to 0.04222 | ns | 0.2108 |
| WT - T91 vs. T91R | -0.241 | -0.4488 to -0.03310 | * | 0.013 |
| WT - T91 vs. T91S | -0.2608 | -0.4687 to -0.05297 | ** | 0.0053 |
| WT - T91 vs. T91L | -0.2856 | -0.5042 to -0.06697 | ** | 0.0032 |
| WT - T91 vs. T91V | -0.2973 | -0.5052 to -0.08942 | *** | 0.0009 |

**Supplementary table 5o. Position 62 (*in vivo* function)**

| Dunnett's multiple comparisons test | Mean Diff. | 95.00% CI of diff. | Summary | Adjusted P Value |
| --- | --- | --- | --- | --- |
| WT-I62 vs. I62N | 0.215 | -0.01448 to 0.4445 | ns | 0.0827 |
| WT-I62 vs. I62G | -0.08409 | -0.3083 to 0.1402 | ns | 0.9694 |
| WT-I62 vs. I62V | -0.5274 | -0.7767 to -0.2780 | **** | <0.0001 |
| WT-I62 vs. I62D | 0.3475 | 0.1122 to 0.5828 | *** | 0.0004 |
| WT-I62 vs. I62R | 0.344 | 0.05087 to 0.6372 | * | 0.0106 |
| WT-I62 vs. I62F | -0.0742 | -0.2984 to 0.1501 | ns | 0.9885 |
| WT-I62 vs. I62K | 0.09184 | -0.2013 to 0.3850 | ns | 0.991 |
| WT-I62 vs. I62L | -0.07062 | -0.3638 to 0.2225 | ns | 0.9991 |
| WT-I62 vs. I62Y | 0.2856 | 0.006324 to 0.5650 | * | 0.0414 |
| WT-I62 vs. I62W | 0.1229 | -0.1066 to 0.3524 | ns | 0.749 |
| WT-I62 vs. I62A | 0.6291 | 0.3797 to 0.8784 | **** | <0.0001 |
| WT-I62 vs. I62Q | 0.3918 | 0.09864 to 0.6850 | ** | 0.002 |

**Supplementary table 5p. Position 65 (*in vivo* function)**

| Dunnett's multiple comparisons test | Mean Diff. | 95.00% CI of diff. | Summary | Adjusted P Value |
| --- | --- | --- | --- | --- |
| WT - Y65 vs. Y65V | 0.4441 | 0.08845 to 0.7998 | ** | 0.0069 |
| WT - Y65 vs. Y65E | 1.051 | 0.6950 to 1.406 | **** | <0.0001 |
| WT - Y65 vs. Y65P | 0.7207 | 0.3794 to 1.062 | **** | <0.0001 |
| WT - Y65 vs. Y65A | 1.072 | 0.7311 to 1.414 | **** | <0.0001 |
| WT - Y65 vs. Y65H | 1.199 | 0.8576 to 1.540 | **** | <0.0001 |
| WT - Y65 vs. Y65D | 0.9428 | 0.5871 to 1.298 | **** | <0.0001 |
| WT - Y65 vs. Y65R | 0.1986 | -0.1753 to 0.5725 | ns | 0.6452 |
| WT - Y65 vs. Y65S | 1.005 | 0.6313 to 1.379 | **** | <0.0001 |
| WT - Y65 vs. Y65T | 0.5949 | 0.2392 to 0.9505 | *** | 0.0001 |

**Supplementary table 5q. Position 70 (*in vivo* function)**

| Dunnett's multiple comparisons test | Mean Diff. | 95.00% CI of diff. | Summary | Adjusted P Value |
| --- | --- | --- | --- | --- |
| WT - T70 vs. T70D | 0.3386 | 0.1188 to 0.5584 | *** | 0.0002 |
| WT - T70 vs. T70K | 0.2716 | 0.07689 to 0.4664 | ** | 0.001 |
| WT - T70 vs. T70E | 0.2435 | 0.05340 to 0.4336 | ** | 0.0036 |
| WT - T70 vs. T70A | 0.2556 | 0.06083 to 0.4503 | ** | 0.0026 |
| WT - T70 vs. T70P | -0.1162 | -0.3110 to 0.07855 | ns | 0.6162 |
| WT - T70 vs. T70R | 0.007149 | -0.1928 to 0.2071 | ns | >0.9999 |
| WT - T70 vs. T70Y | 0.06366 | -0.1363 to 0.2636 | ns | 0.9906 |
| WT - T70 vs. T70L | -0.08882 | -0.2836 to 0.1059 | ns | 0.891 |
| WT - T70 vs. T70Q | -0.04656 | -0.2413 to 0.1482 | ns | 0.9991 |
| WT - T70 vs. T70F | -0.1054 | -0.3252 to 0.1145 | ns | 0.8561 |
| WT - T70 vs. T70V | -0.2143 | -0.4266 to -0.001981 | * | 0.0462 |
| WT - T70 vs. T70S | 0.05532 | -0.1570 to 0.2676 | ns | 0.999 |

**Supplementary table 5r. Position 74 (*in vivo* function)**

| Dunnett's multiple comparisons test | Mean Diff. | 95.00% CI of diff. | Summary | Adjusted P Value |
| --- | --- | --- | --- | --- |
| WT - N74 vs. N74V | 0.7221 | 0.5351 to 0.9091 | **** | <0.0001 |
| WT - N74 vs. N74L | 0.4317 | 0.2447 to 0.6187 | **** | <0.0001 |
| WT - N74 vs. N74F | 0.3236 | 0.1367 to 0.5106 | **** | <0.0001 |
| WT - N74 vs. N74A | 0.2782 | 0.08410 to 0.4723 | *** | 0.0006 |
| WT - N74 vs. N74P | -0.06557 | -0.2119 to 0.08081 | ns | 0.9207 |
| WT - N74 vs. N74T | 0.008916 | -0.1570 to 0.1748 | ns | 0.9999 |
| WT - N74 vs. N74M | -0.02686 | -0.1708 to 0.1171 | ns | 0.9994 |
| WT - N74 vs. N74D | 0.1497 | -0.02060 to 0.3200 | ns | 0.1312 |
| WT - N74 vs. N74R | -0.08519 | -0.2291 to 0.05875 | ns | 0.6552 |
| WT - N74 vs. N74E | 0.09119 | -0.09579 to 0.2782 | ns | 0.866 |
| WT - N74 vs. N74Y | -0.06963 | -0.2355 to 0.09625 | ns | 0.9491 |
| WT - N74 vs. N74K | 0.00698 | -0.1270 to 0.1409 | ns | 0.9999 |
| WT - N74 vs. N74S | 0.04698 | -0.1079 to 0.2019 | ns | 0.9955 |

**Supplementary table 5s. Position 77 (*in vivo* function)**

| Dunnett's multiple comparisons test | Mean Diff. | 95.00% CI of diff. | Summary | Adjusted P Value |
| --- | --- | --- | --- | --- |
| WT - Q77 vs. Q77E | 0.8575 | 0.5885 to 1.126 | **** | <0.0001 |
| WT - Q77 vs. Q77G | -0.1032 | -0.3859 to 0.1795 | ns | 0.9632 |
| WT - Q77 vs. Q77K | 0.02805 | -0.2409 to 0.2970 | ns | 0.9997 |
| WT - Q77 vs. Q77P | 0.0143 | -0.2547 to 0.2833 | ns | 0.9999 |
| WT - Q77 vs. Q77N | -0.08912 | -0.3581 to 0.1798 | ns | 0.9819 |
| WT - Q77 vs. Q77L | -0.2487 | -0.5177 to 0.02028 | ns | 0.0887 |
| WT - Q77 vs. Q77R | 0.4072 | 0.1382 to 0.6762 | *** | 0.0004 |
| WT - Q77 vs. Q77S | 0.1323 | -0.1367 to 0.4013 | ns | 0.8007 |
| WT - Q77 vs. Q77Y | 0.04385 | -0.2389 to 0.3266 | ns | 0.9995 |
| WT - Q77 vs. Q77T | -0.02399 | -0.2930 to 0.2450 | ns | 0.9997 |
| WT - Q77 vs. Q77V | 0.009306 | -0.2597 to 0.2783 | ns | >0.9999 |

**Supplementary table 5t. Position 78 (*in vivo* function)**

| Dunnett's multiple comparisons test | Mean Diff. | 95.00% CI of diff. | Summary | Adjusted P Value |
| --- | --- | --- | --- | --- |
| WT - F78 vs. F78G | -0.09847 | -0.3461 to 0.1491 | ns | 0.9635 |
| WT - F78 vs. F78S | -0.00549 | -0.2664 to 0.2555 | ns | >0.9999 |
| WT - F78 vs. F78C | -0.03308 | -0.3019 to 0.2358 | ns | 0.9996 |
| WT - F78 vs. F78L | -0.2673 | -0.5212 to -0.01336 | * | 0.0316 |
| WT - F78 vs. F78E | -0.1734 | -0.4210 to 0.07426 | ns | 0.4019 |
| WT - F78 vs. F78P | -0.1105 | -0.3581 to 0.1372 | ns | 0.9187 |
| WT - F78 vs. F78H | -0.2197 | -0.4736 to 0.03422 | ns | 0.1439 |
| WT - F78 vs. F78R | 0.1828 | -0.07112 to 0.4367 | ns | 0.3618 |
| WT - F78 vs. F78I | 0.06819 | -0.1857 to 0.3221 | ns | 0.999 |
| WT - F78 vs. F78V | -0.05766 | -0.3053 to 0.1899 | ns | 0.9992 |
| WT - F78 vs. F78N | -0.1423 | -0.3962 to 0.1116 | ns | 0.7187 |
| WT - F78 vs. F78M | -0.1948 | -0.4424 to 0.05285 | ns | 0.2444 |
| WT - F78 vs. F78Q | 0.3402 | 0.08624 to 0.5941 | ** | 0.0018 |

**Supplementary table 5u. Position 91 (*in vivo* function)**

| Dunnett's multiple comparisons test | Mean Diff. | 95.00% CI of diff. | Summary | Adjusted P Value |
| --- | --- | --- | --- | --- |
| T91G vs. T91A | 0.06305 | -0.1827 to 0.3088 | ns | 0.9914 |
| T91G vs. WT - T91 | -0.2126 | -0.4143 to -0.01089 | * | 0.0336 |
| T91G vs. T91F | -0.1461 | -0.3918 to 0.09964 | ns | 0.4988 |
| T91G vs. T91H | 0.005836 | -0.2399 to 0.2515 | ns | >0.9999 |
| T91G vs. T91E | -0.08713 | -0.3328 to 0.1586 | ns | 0.9368 |
| T91G vs. T91Q | -0.1099 | -0.3557 to 0.1358 | ns | 0.8 |
| T91G vs. T91P | -0.1297 | -0.3754 to 0.1160 | ns | 0.6367 |
| T91G vs. T91R | -0.1458 | -0.3915 to 0.09990 | ns | 0.501 |
| T91G vs. T91S | -0.2166 | -0.4624 to 0.02907 | ns | 0.1113 |
| T91G vs. T91L | -0.2057 | -0.4600 to 0.04866 | ns | 0.1734 |
| T91G vs. T91V | -0.04327 | -0.2890 to 0.2024 | ns | 0.9993 |

**Supplementary Table 6.** **False discovery rate** analyses for measured experimental outcomes for amino acid variants relative to the WT sample measured in parallel. **(a)**-**(g)** cellular concentration **(h)-(n)** transcriptional activation **(o)-(u)** combined effects.

**Supplementary table 6a. Position 62 (Cellular CIITA concentration)**

| Two-stage linear step-up procedure of Benjamini | Krieger and Yekutieli | Mean Diff. | Discovery? | q value |
| --- | --- | --- | --- | --- |
| WT-I62 vs. I62G | -0.159 | Yes | 0.0412 | 0.0196 |
| WT-I62 vs. I62V | -0.02454 | No | 0.6429 | 0.7439 |
| WT-I62 vs. I62D | 0.09218 | No | 0.2825 | 0.1946 |
| WT-I62 vs. I62R | 0.04965 | No | 0.6429 | 0.5741 |
| WT-I62 vs. I62F | 0.02118 | No | 0.6429 | 0.7539 |
| WT-I62 vs. I62K | 0.01762 | No | 0.6429 | 0.8419 |
| WT-I62 vs. I62L | 0.01929 | No | 0.6429 | 0.8271 |
| WT-I62 vs. I62Y | 0.4707 | Yes | <0.0001 | <0.0001 |
| WT-I62 vs. I62W | 0.08644 | No | 0.2825 | 0.2018 |
| WT-I62 vs. I62A | 0.4291 | Yes | <0.0001 | <0.0001 |
| WT-I62 vs. I62Q | 0.8519 | Yes | <0.0001 | <0.0001 |

**Supplementary table 6b. Position 65 (Cellular CIITA concentration)**

| Two-stage linear step-up procedure of Benjamini | Krieger and Yekutieli | Mean Diff. | Discovery? | q value |
| --- | --- | --- | --- | --- |
| WT - Y65 vs. Y65E | 0.6392 | Yes | <0.0001 | <0.0001 |
| WT - Y65 vs. Y65P | 0.4884 | Yes | <0.0001 | <0.0001 |
| WT - Y65 vs. Y65A | -0.2558 | Yes | 0.0124 | 0.0197 |
| WT - Y65 vs. Y65H | -0.1224 | No | 0.1572 | 0.2994 |
| WT - Y65 vs. Y65D | -0.08845 | No | 0.1848 | 0.4106 |
| WT - Y65 vs. Y65R | 0.03609 | No | 0.2987 | 0.7586 |
| WT - Y65 vs. Y65S | 0.7642 | Yes | <0.0001 | <0.0001 |
| WT - Y65 vs. Y65T | 0.3316 | Yes | 0.0033 | 0.0042 |

**Supplementary table 6c. Position 70 (Cellular CIITA concentration)**

| Two-stage linear step-up procedure of Benjamini | Krieger and Yekutieli | Mean Diff. | Discovery? | q value |
| --- | --- | --- | --- | --- |
| WT - T70 vs. T70D | 0.1984 | Yes | 0.0034 | 0.0004 |
| WT - T70 vs. T70K | 0.153 | Yes | 0.0061 | 0.0022 |
| WT - T70 vs. T70E | 0.04412 | No | 0.3127 | 0.3722 |
| WT - T70 vs. T70A | 0.1577 | Yes | 0.0061 | 0.0016 |
| WT - T70 vs. T70P | -0.07148 | No | 0.1564 | 0.1489 |
| WT - T70 vs. T70R | -0.105 | Yes | 0.0493 | 0.0345 |
| WT - T70 vs. T70Y | 0.1046 | Yes | 0.0493 | 0.0352 |
| WT - T70 vs. T70L | 0.1378 | Yes | 0.012 | 0.0057 |
| WT - T70 vs. T70Q | 0.01565 | No | 0.526 | 0.7514 |
| WT - T70 vs. T70F | 0.07238 | No | 0.1784 | 0.1912 |
| WT - T70 vs. T70V | 0.04363 | No | 0.3285 | 0.4302 |
| WT - T70 vs. T70S | 0.1208 | No | 0.0792 | 0.066 |

**Supplementary table 6d. Position 74 (Cellular CIITA concentration)**

| Two-stage linear step-up procedure of Benjamini | Krieger and Yekutieli | Mean Diff. | Discovery? | q value |
| --- | --- | --- | --- | --- |
| WT - N74 vs. N74V | -0.01039 | No | 0.4654 | 0.8232 |
| WT - N74 vs. N74L | -0.1443 | Yes | 0.0005 | 0.0003 |
| WT - N74 vs. N74F | 0.0506 | No | 0.1427 | 0.1941 |
| WT - N74 vs. N74A | 0.144 | Yes | 0.0005 | 0.0003 |
| WT - N74 vs. N74P | -0.1602 | Yes | <0.0001 | <0.0001 |
| WT - N74 vs. N74T | -0.09992 | Yes | 0.005 | 0.0041 |
| WT - N74 vs. N74M | -0.1317 | Yes | 0.0001 | <0.0001 |
| WT - N74 vs. N74D | 0.05193 | No | 0.124 | 0.1331 |
| WT - N74 vs. N74R | -0.09146 | Yes | 0.0031 | 0.0021 |
| WT - N74 vs. N74E | 0.02063 | No | 0.3651 | 0.596 |
| WT - N74 vs. N74Y | 0.04963 | No | 0.124 | 0.1511 |
| WT - N74 vs. N74K | 0.03557 | No | 0.2413 | 0.3611 |
| WT - N74 vs. N74S | 0.04624 | No | 0.124 | 0.1519 |

**Supplementary table 6e. Position 77 (Cellular CIITA concentration)**

| Two-stage linear step-up procedure of Benjamini | Krieger and Yekutieli | Mean Diff. | Discovery? | q value |
| --- | --- | --- | --- | --- |
| WT - Q77 vs. Q77E | -0.05972 | No | 0.2953 | 0.5157 |
| WT - Q77 vs. Q77G | -0.173 | No | 0.071 | 0.0752 |
| WT - Q77 vs. Q77K | -0.1625 | No | 0.071 | 0.0789 |
| WT - Q77 vs. Q77P | 0.1061 | No | 0.157 | 0.2493 |
| WT - Q77 vs. Q77N | 0.1138 | No | 0.1518 | 0.2168 |
| WT - Q77 vs. Q77L | -0.1169 | No | 0.1518 | 0.2047 |
| WT - Q77 vs. Q77R | 0.286 | Yes | 0.0037 | 0.0024 |
| WT - Q77 vs. Q77S | 0.2702 | Yes | 0.0076 | 0.006 |
| WT - Q77 vs. Q77Y | 0.3293 | Yes | 0.0019 | 0.0009 |
| WT - Q77 vs. Q77T | 0.3334 | Yes | 0.0014 | 0.0004 |
| WT - Q77 vs. Q77V | 0.4317 | Yes | <0.0001 | <0.0001 |

**Supplementary table 6f. Position 78 (Cellular CIITA concentration)**

| Two-stage linear step-up procedure of Benjamini | Krieger and Yekutieli | Mean Diff. | Discovery? | q value |
| --- | --- | --- | --- | --- |
| WT - F78 vs. F78S | 0.07881 | No | 0.1434 | 0.1821 |
| WT - F78 vs. F78C | -0.05095 | No | 0.2517 | 0.3995 |
| WT - F78 vs. F78L | -0.08922 | No | 0.1267 | 0.1408 |
| WT - F78 vs. F78E | -0.00803 | No | 0.4681 | 0.8917 |
| WT - F78 vs. F78H | -0.2364 | Yes | 0.0002 | 0.0002 |
| WT - F78 vs. F78R | 0.5383 | Yes | <0.0001 | <0.0001 |
| WT - F78 vs. F78P | 0.06067 | No | 0.2128 | 0.304 |
| WT - F78 vs. F78I | -0.2042 | Yes | 0.0012 | 0.0012 |
| WT - F78 vs. F78V | 0.3647 | Yes | <0.0001 | <0.0001 |
| WT - F78 vs. F78N | -0.01164 | No | 0.4681 | 0.8472 |
| WT - F78 vs. F78M | 0.326 | Yes | <0.0001 | <0.0001 |
| WT - F78 vs. F78Q | 0.6003 | Yes | <0.0001 | <0.0001 |

**Supplementary table 6g. Position 91 (Cellular CIITA concentration)**

| Two-stage linear step-up procedure of Benjamini | Krieger and Yekutieli | Mean Diff. | Discovery? | q value |
| --- | --- | --- | --- | --- |
| WT - T91 vs. T91G | -0.03171 | No | 0.7679 | 0.6501 |
| WT - T91 vs. T91F | 0.04317 | No | 0.7679 | 0.5371 |
| WT - T91 vs. T91H | -0.0013 | No | 0.931 | 0.9852 |
| WT - T91 vs. T91E | -0.03201 | No | 0.7679 | 0.647 |
| WT - T91 vs. T91Q | -0.03549 | No | 0.7679 | 0.6118 |
| WT - T91 vs. T91P | -0.2057 | Yes | 0.0377 | 0.004 |
| WT - T91 vs. T91R | -0.1476 | No | 0.174 | 0.0368 |
| WT - T91 vs. T91S | 0.01704 | No | 0.8477 | 0.8073 |
| WT - T91 vs. T91L | 0.0596 | No | 0.7679 | 0.3945 |
| WT - T91 vs. T91V | 0.12 | No | 0.2782 | 0.0883 |

**Supplementary table 6h. Position 62 (Transcriptional activation)**

| Two-stage linear step-up procedure of Benjamini | Krieger and Yekutieli | Mean Diff. | Discovery? | q value |
| --- | --- | --- | --- | --- |
| WT-I62 vs. I62N | 0.4573 | Yes | <0.0001 | <0.0001 |
| WT-I62 vs. I62G | 0.2595 | Yes | <0.0001 | <0.0001 |
| WT-I62 vs. I62V | 0.127 | Yes | 0.0432 | 0.0353 |
| WT-I62 vs. I62D | 0.2867 | Yes | <0.0001 | <0.0001 |
| WT-I62 vs. I62R | 0.08209 | No | 0.18 | 0.2448 |
| WT-I62 vs. I62F | 0.03683 | No | 0.3305 | 0.4946 |
| WT-I62 vs. I62K | 0.000832 | No | 0.6067 | 0.9906 |
| WT-I62 vs. I62L | -0.1069 | No | 0.1065 | 0.1305 |
| WT-I62 vs. I62Y | -0.09405 | No | 0.1065 | 0.107 |
| WT-I62 vs. I62W | -0.08644 | No | 0.1065 | 0.1183 |
| WT-I62 vs. I62A | -0.2815 | Yes | <0.0001 | <0.0001 |
| WT-I62 vs. I62Q | -0.3813 | Yes | <0.0001 | <0.0001 |

**Supplementary table 6i. Position 65 (Transcriptional activation)**

| Two-stage linear step-up procedure of Benjamini | Krieger and Yekutieli | Mean Diff. | Discovery? | q value |
| --- | --- | --- | --- | --- |
| WT - Y65 vs. Y65V | -0.1248 | No | 0.2114 | 0.2013 |
| WT - Y65 vs. Y65E | 0.1939 | No | 0.0583 | 0.037 |
| WT - Y65 vs. Y65P | -0.2649 | Yes | 0.0067 | 0.0032 |
| WT - Y65 vs. Y65A | 0.0589 | No | 0.397 | 0.5003 |
| WT - Y65 vs. Y65H | 0.4196 | Yes | <0.0001 | <0.0001 |
| WT - Y65 vs. Y65D | 0.01866 | No | 0.5815 | 0.8307 |
| WT - Y65 vs. Y65R | -0.7852 | Yes | <0.0001 | <0.0001 |
| WT - Y65 vs. Y65S | -0.06495 | No | 0.397 | 0.5041 |
| WT - Y65 vs. Y65T | -0.1518 | No | 0.1267 | 0.1006 |

**Supplementary table 6j. Position 70 (Transcriptional activation)**

| Two-stage linear step-up procedure of Benjamini | Krieger and Yekutieli | Mean Diff. | Discovery? | q value |
| --- | --- | --- | --- | --- |
| WT - T70 vs. T70D | 0.1593 | Yes | 0.0471 | 0.0336 |
| WT - T70 vs. T70K | 0.07569 | No | 0.1608 | 0.1531 |
| WT - T70 vs. T70E | 0.02153 | No | 0.5728 | 0.75 |
| WT - T70 vs. T70A | 0.004874 | No | 0.6514 | 0.9305 |
| WT - T70 vs. T70P | -0.04649 | No | 0.4477 | 0.4796 |
| WT - T70 vs. T70R | -0.09513 | No | 0.092 | 0.0766 |
| WT - T70 vs. T70Y | -0.03362 | No | 0.4655 | 0.5542 |
| WT - T70 vs. T70L | -0.1438 | Yes | 0.0471 | 0.0295 |
| WT - T70 vs. T70Q | -0.1766 | Yes | 0.016 | 0.0076 |
| WT - T70 vs. T70F | -0.2729 | Yes | 0.0009 | 0.0003 |
| WT - T70 vs. T70V | -0.3095 | Yes | 0.0001 | <0.0001 |
| WT - T70 vs. T70S | -0.4265 | Yes | <0.0001 | <0.0001 |

**Supplementary table 6k. Position 74 (Transcriptional activation)**

| Two-stage linear step-up procedure of Benjamini | Krieger and Yekutieli | Mean Diff. | Discovery? | q value |
| --- | --- | --- | --- | --- |
| WT - N74 vs. N74V | 0.5329 | Yes | <0.0001 | <0.0001 |
| WT - N74 vs. N74L | 0.2666 | Yes | 0.0015 | 0.0004 |
| WT - N74 vs. N74F | 0.1998 | Yes | 0.0185 | 0.0078 |
| WT - N74 vs. N74A | 0.1678 | Yes | 0.0488 | 0.031 |
| WT - N74 vs. N74P | 0.07199 | No | 0.2588 | 0.2351 |
| WT - N74 vs. N74T | 0.04631 | No | 0.4576 | 0.4842 |
| WT - N74 vs. N74M | 0.02543 | No | 0.5313 | 0.6746 |
| WT - N74 vs. N74D | -0.02206 | No | 0.5418 | 0.7454 |
| WT - N74 vs. N74R | -0.06897 | No | 0.2588 | 0.2465 |
| WT - N74 vs. N74E | -0.04402 | No | 0.4769 | 0.5552 |
| WT - N74 vs. N74Y | -0.1517 | Yes | 0.0427 | 0.0226 |
| WT - N74 vs. N74K | -0.111 | No | 0.1857 | 0.1376 |
| WT - N74 vs. N74S | -0.2181 | Yes | 0.0015 | 0.0005 |

**Supplementary table 6l. Position 77 (Transcriptional activation)**

| Two-stage linear step-up procedure of Benjamini | Krieger and Yekutieli | Mean Diff. | Discovery? | q value |
| --- | --- | --- | --- | --- |
| WT - Q77 vs. Q77E | 0.8114 | Yes | <0.0001 | <0.0001 |
| WT - Q77 vs. Q77G | 0.3176 | Yes | 0.0001 | 0.0002 |
| WT - Q77 vs. Q77K | 0.271 | Yes | 0.0004 | 0.0007 |
| WT - Q77 vs. Q77P | 0.214 | Yes | 0.0031 | 0.0069 |
| WT - Q77 vs. Q77N | 0.1108 | No | 0.0547 | 0.1562 |
| WT - Q77 vs. Q77L | -0.06003 | No | 0.1261 | 0.4405 |
| WT - Q77 vs. Q77R | -0.09686 | No | 0.0675 | 0.2144 |
| WT - Q77 vs. Q77S | -0.1866 | Yes | 0.0095 | 0.0242 |
| WT - Q77 vs. Q77Y | -0.3102 | Yes | 0.0002 | 0.0002 |
| WT - Q77 vs. Q77T | -0.3229 | Yes | <0.0001 | <0.0001 |
| WT - Q77 vs. Q77V | -0.3843 | Yes | <0.0001 | <0.0001 |

**Supplementary table 6m. Position 78 (Transcriptional activation)**

| Two-stage linear step-up procedure of Benjamini | Krieger and Yekutieli | Mean Diff. | Discovery? | q value |
| --- | --- | --- | --- | --- |
| WT - F78 vs. F78G | 0.4111 | Yes | <0.0001 | <0.0001 |
| WT - F78 vs. F78S | 0.3036 | Yes | 0.0002 | 0.0001 |
| WT - F78 vs. F78C | 0.2252 | Yes | 0.0014 | 0.0011 |
| WT - F78 vs. F78L | 0.2479 | Yes | 0.0011 | 0.0007 |
| WT - F78 vs. F78E | 0.1697 | Yes | 0.0124 | 0.0138 |
| WT - F78 vs. F78P | 0.09269 | No | 0.1391 | 0.1767 |
| WT - F78 vs. F78H | 0.08563 | No | 0.1688 | 0.2502 |
| WT - F78 vs. F78R | 0.08005 | No | 0.1688 | 0.268 |
| WT - F78 vs. F78I | 0.01858 | No | 0.3861 | 0.7968 |
| WT - F78 vs. F78V | 0.02022 | No | 0.3861 | 0.7677 |
| WT - F78 vs. F78N | -0.04116 | No | 0.3195 | 0.5579 |
| WT - F78 vs. F78M | -0.2014 | Yes | 0.0037 | 0.0036 |
| WT - F78 vs. F78Q | -0.3043 | Yes | 0.0002 | <0.0001 |

**Supplementary table 6n. Position 91 (Transcriptional activation)**

| Two-stage linear step-up procedure of Benjamini | Krieger and Yekutieli | Mean Diff. | Discovery? | q value |
| --- | --- | --- | --- | --- |
| WT - T91 vs. T91A | 0.1735 | Yes | 0.0195 | 0.0185 |
| WT - T91 vs. T91G | -0.02546 | No | 0.3635 | 0.726 |
| WT - T91 vs. T91F | -0.02204 | No | 0.3635 | 0.7616 |
| WT - T91 vs. T91H | -0.05485 | No | 0.2629 | 0.4508 |
| WT - T91 vs. T91E | -0.07868 | No | 0.1838 | 0.28 |
| WT - T91 vs. T91Q | -0.1265 | No | 0.0629 | 0.0839 |
| WT - T91 vs. T91P | -0.1656 | Yes | 0.0213 | 0.0244 |
| WT - T91 vs. T91R | -0.241 | Yes | 0.0016 | 0.0012 |
| WT - T91 vs. T91S | -0.2608 | Yes | 0.0009 | 0.0005 |
| WT - T91 vs. T91L | -0.2856 | Yes | 0.0008 | 0.0003 |
| WT - T91 vs. T91V | -0.2973 | Yes | 0.0004 | <0.0001 |

**Supplementary table 6o. Position 62 (*in vivo* function)**

| Two-stage linear step-up procedure of Benjamini | Krieger and Yekutieli | Mean Diff. | Discovery? | q value |
| --- | --- | --- | --- | --- |
| WT-I62 vs. I62N | 0.215 | Yes | 0.0058 | 0.0077 |
| WT-I62 vs. I62G | -0.08409 | No | 0.1646 | 0.2822 |
| WT-I62 vs. I62V | -0.5274 | Yes | <0.0001 | <0.0001 |
| WT-I62 vs. I62D | 0.3475 | Yes | <0.0001 | <0.0001 |
| WT-I62 vs. I62R | 0.344 | Yes | 0.001 | 0.0009 |
| WT-I62 vs. I62F | -0.0742 | No | 0.1759 | 0.3424 |
| WT-I62 vs. I62K | 0.09184 | No | 0.1759 | 0.3686 |
| WT-I62 vs. I62L | -0.07062 | No | 0.214 | 0.4892 |
| WT-I62 vs. I62Y | 0.2856 | Yes | 0.0033 | 0.0037 |
| WT-I62 vs. I62W | 0.1229 | No | 0.0821 | 0.1251 |
| WT-I62 vs. I62A | 0.6291 | Yes | <0.0001 | <0.0001 |
| WT-I62 vs. I62Q | 0.3918 | Yes | 0.0002 | 0.0002 |

**Supplementary Table 6p. Position 65 (*in vivo* function)**

| Two-stage linear step-up procedure of Benjamini | Krieger and Yekutieli | Mean Diff. | Discovery? | q value |
| --- | --- | --- | --- | --- |
| WT - Y65 vs. Y65V | 0.4441 | Yes | 0.0001 | 0.0008 |
| WT - Y65 vs. Y65E | 1.051 | Yes | <0.0001 | <0.0001 |
| WT - Y65 vs. Y65P | 0.7207 | Yes | <0.0001 | <0.0001 |
| WT - Y65 vs. Y65A | 1.072 | Yes | <0.0001 | <0.0001 |
| WT - Y65 vs. Y65H | 1.199 | Yes | <0.0001 | <0.0001 |
| WT - Y65 vs. Y65D | 0.9428 | Yes | <0.0001 | <0.0001 |
| WT - Y65 vs. Y65R | 0.1986 | Yes | 0.0165 | 0.141 |
| WT - Y65 vs. Y65S | 1.005 | Yes | <0.0001 | <0.0001 |
| WT - Y65 vs. Y65T | 0.5949 | Yes | <0.0001 | <0.0001 |

**Supplementary Table 6q. Position 70 (*in vivo* function)**

| Two-stage linear step-up procedure of Benjamini | Krieger and Yekutieli | Mean Diff. | Discovery? | q value |
| --- | --- | --- | --- | --- |
| WT - T70 vs. T70D | 0.3386 | Yes | 0.0001 | <0.0001 |
| WT - T70 vs. T70K | 0.2716 | Yes | 0.0003 | <0.0001 |
| WT - T70 vs. T70E | 0.2435 | Yes | 0.0006 | 0.0003 |
| WT - T70 vs. T70A | 0.2556 | Yes | 0.0005 | 0.0002 |
| WT - T70 vs. T70P | -0.1162 | No | 0.108 | 0.0882 |
| WT - T70 vs. T70R | 0.007149 | No | 0.5625 | 0.9183 |
| WT - T70 vs. T70Y | 0.06366 | No | 0.2953 | 0.3616 |
| WT - T70 vs. T70L | -0.08882 | No | 0.1762 | 0.1918 |
| WT - T70 vs. T70Q | -0.04656 | No | 0.3295 | 0.4932 |
| WT - T70 vs. T70F | -0.1054 | No | 0.1762 | 0.1703 |
| WT - T70 vs. T70V | -0.2143 | Yes | 0.0061 | 0.0042 |
| WT - T70 vs. T70S | 0.05532 | No | 0.3295 | 0.4552 |

**Supplementary Table 6r. Position 74 (*in vivo* function)**

| Two-stage linear step-up procedure of Benjamini | Krieger and Yekutieli | Mean Diff. | Discovery? | q value |
| --- | --- | --- | --- | --- |
| WT - N74 vs. N74V | 0.7221 | Yes | <0.0001 | <0.0001 |
| WT - N74 vs. N74L | 0.4317 | Yes | <0.0001 | <0.0001 |
| WT - N74 vs. N74F | 0.3236 | Yes | <0.0001 | <0.0001 |
| WT - N74 vs. N74A | 0.2782 | Yes | <0.0001 | <0.0001 |
| WT - N74 vs. N74P | -0.06557 | No | 0.2058 | 0.196 |
| WT - N74 vs. N74T | 0.008916 | No | 0.5688 | 0.8765 |
| WT - N74 vs. N74M | -0.02686 | No | 0.4503 | 0.5896 |
| WT - N74 vs. N74D | 0.1497 | Yes | 0.0194 | 0.0115 |
| WT - N74 vs. N74R | -0.08519 | No | 0.1231 | 0.0879 |
| WT - N74 vs. N74E | 0.09119 | No | 0.1912 | 0.1593 |
| WT - N74 vs. N74Y | -0.06963 | No | 0.2105 | 0.2256 |
| WT - N74 vs. N74K | 0.00698 | No | 0.5688 | 0.8803 |
| WT - N74 vs. N74S | 0.04698 | No | 0.3199 | 0.3809 |

**Supplementary Table 6s. Position 77 (*in vivo* function)**

| Two-stage linear step-up procedure of Benjamini | Krieger and Yekutieli | Mean Diff. | Discovery? | q value |
| --- | --- | --- | --- | --- |
| WT - Q77 vs. Q77E | 0.8575 | Yes | <0.0001 | <0.0001 |
| WT - Q77 vs. Q77G | -0.1032 | No | 0.4816 | 0.2973 |
| WT - Q77 vs. Q77K | 0.02805 | No | 0.7034 | 0.7654 |
| WT - Q77 vs. Q77P | 0.0143 | No | 0.7034 | 0.879 |
| WT - Q77 vs. Q77N | -0.08912 | No | 0.4816 | 0.344 |
| WT - Q77 vs. Q77L | -0.2487 | Yes | 0.0261 | 0.0093 |
| WT - Q77 vs. Q77R | 0.4072 | Yes | 0.0001 | <0.0001 |
| WT - Q77 vs. Q77S | 0.1323 | No | 0.3388 | 0.1613 |
| WT - Q77 vs. Q77Y | 0.04385 | No | 0.7034 | 0.6573 |
| WT - Q77 vs. Q77T | -0.02399 | No | 0.7034 | 0.7985 |
| WT - Q77 vs. Q77V | 0.009306 | No | 0.7034 | 0.9211 |

**Supplementary Table 6t. Position 78 (*in vivo* function)**

| Two-stage linear step-up procedure of Benjamini | Krieger and Yekutieli | Mean Diff. | Discovery? | q value |
| --- | --- | --- | --- | --- |
| WT - F78 vs. F78G | -0.09847 | No | 0.322 | 0.2509 |
| WT - F78 vs. F78S | -0.00549 | No | 0.8454 | 0.9515 |
| WT - F78 vs. F78C | -0.03308 | No | 0.695 | 0.7221 |
| WT - F78 vs. F78L | -0.2673 | Yes | 0.0149 | 0.0026 |
| WT - F78 vs. F78E | -0.1734 | No | 0.0844 | 0.0439 |
| WT - F78 vs. F78P | -0.1105 | No | 0.2858 | 0.1979 |
| WT - F78 vs. F78H | -0.2197 | Yes | 0.0499 | 0.0129 |
| WT - F78 vs. F78R | 0.1828 | No | 0.0844 | 0.0383 |
| WT - F78 vs. F78I | 0.06819 | No | 0.5057 | 0.4378 |
| WT - F78 vs. F78V | -0.05766 | No | 0.526 | 0.501 |
| WT - F78 vs. F78N | -0.1423 | No | 0.1752 | 0.1062 |
| WT - F78 vs. F78M | -0.1948 | No | 0.0685 | 0.0237 |
| WT - F78 vs. F78Q | 0.3402 | Yes | 0.0016 | 0.0001 |

**Supplementary Table 6u. Position 91 (*in vivo* function)**

| Two-stage linear step-up procedure of Benjamini | Krieger and Yekutieli | Mean Diff. | Discovery? | q value |
| --- | --- | --- | --- | --- |
| T91G vs. T91A | 0.06305 | No | 0.5556 | 0.4762 |
| T91G vs. WT - T91 | -0.2126 | Yes | 0.0433 | 0.0041 |
| T91G vs. T91F | -0.1461 | No | 0.2128 | 0.1007 |
| T91G vs. T91H | 0.005836 | No | 0.9043 | 0.9474 |
| T91G vs. T91E | -0.08713 | No | 0.4271 | 0.3254 |
| T91G vs. T91Q | -0.1099 | No | 0.323 | 0.2153 |
| T91G vs. T91P | -0.1297 | No | 0.2526 | 0.1443 |
| T91G vs. T91R | -0.1458 | No | 0.2128 | 0.1013 |
| T91G vs. T91S | -0.2166 | No | 0.0827 | 0.0158 |
| T91G vs. T91L | -0.2057 | No | 0.0925 | 0.0264 |
| T91G vs. T91V | -0.04327 | No | 0.6559 | 0.6246 |

**Supplementary table 7. Calculations for the lower limit of the tunable range (hypothetical “dead” variant) for each measured experimental outcome, on the WT-normalized scale.**

|  | Average, unnormalized background signal in the absence of CIITA | Estimated threshold multiplier^b^ | Lower limit of the tunable range (hypothetical “dead” variant)^c^ |
| --- | --- | --- | --- |
| Cellular concentration | 0.1^a^ | ------ | ------ |
| Transcriptional activation | 0.8 | 0.1 | **0.08** |
| Combined effects | 3 | 0.05 | **0.15** |

^a^ Concentration: For a sample with no CIITA, the background signal on the western blot for an area comparable to that of the WT band had a signal that was ~10% of the WT signal.

^b^ Average for variant with smallest normalized value of each parameter. This value was used to identify the highest possible threshold below which activity is not detected. The “dead” variant must be smaller than that of the least detected activity (This threshold could actually be lower).

^c^ Product of the unnormalized signal and the multiplier
